# Fibroblast-derived Osteoglycin Promotes Epithelial Cell Repair

**DOI:** 10.1101/2024.11.14.623606

**Authors:** Luke van der Koog, Manon E. Woest, Iris C. Gorter, Vicky Verschut, Robin A.B. Elferink, Annet B. Zuidhof, Dyan F. Nugraha, Maunick L. Koloko Ngassie, Sophie I.T. Bos, Deepesh Dhakad, Justina C. Wolters, Peter L. Horvatovich, Y.S. Prakash, Wim Timens, Önder A. Yildirim, Corry-Anke Brandsma, Henderik W. Frijlink, Anika Nagelkerke, Reinoud Gosens

## Abstract

There is an urgent need for innovative pharmacological treatments targeting defective epithelial repair in chronic diseases, such as chronic obstructive pulmonary disease. The mesenchymal niche is a critical regulator in epithelial stem cell activation during repair. We hypothesized that secreted factors in this interaction are potent drug targets. Utilizing a cutting-edge proteomics-guided drug discovery strategy, we explored the lung fibroblast secretome to uncover impactful drug targets. Our lung organoid assays identified several regenerative ligands, with the secreted matrix protein osteoglycin (OGN) surprisingly showing the most profound effects. Transcriptomic analyses revealed that OGN enhances alveolar progenitor cell differentiation, boosts reactive oxygen species detoxification, reduces cellular senescence, and strengthens fibroblast-epithelial crosstalk. Critically, OGN expression was diminished in COPD patients and smoke-exposed mice. An active fragment of OGN, encompassing leucine-rich repeat regions 4-7, demonstrated regenerative potential akin to full-length OGN. This fragment significantly ameliorated elastase-induced lung injury precision-cut lung slices and improved lung function *in vivo*. These findings highlight lung fibroblast-derived OGN as a pivotal secreted protein for alveolar epithelial growth, positioning its active fragment as a promising therapeutic for epithelial repair in individuals with accelerated lung tissue damage.

## INTRODUCTION

There is a growing interest in therapeutic strategies aimed at restoring defective tissue repair in chronic diseases to cure or slow down the decline in organ function. In the lung, regenerative medicine has primarily focused on transplantation, tissue engineering, stem or progenitor cell therapy, or a combination of these approaches.^1,2^ A regenerative pharmacological approach holds considerable promise due to its potential for large-scale application and early intervention in the disease process.^3^ Currently, no approved drugs support lung tissue repair.

Within respiratory medicine, chronic obstructive pulmonary disease (COPD) is the most common chronic lung disease with a need for better therapies.^2,4^ The underlying pathogenetic mechanism of COPD involves a complex interplay between enhanced destruction of alveolar septa and a diminished capacity of alveolar epithelial progenitor cells to support epithelial tissue repair.^5–7^ The alveolar epithelium comprises alveolar type (AT)1 cells, which facilitate gas exchange, and AT2 cells, which secrete pulmonary surfactants and possess stem cell-like characteristics essential for lung repair.^8,9^ Cell types within the alveolar niche, such as distal lung fibroblasts, provide structural support, secrete extracellular matrix, and provide paracrine molecular cues that direct proper proliferation and differentiation of the alveolar epithelium.^9–13^

In COPD, aberrant fibroblast-epithelial signaling contributes to remodeling and impaired epithelial repair.^14–18^ Recent findings show that the progenitor cell niche is a driver of defective epithelial repair and that targeting the niche pharmacologically is feasible.^3,19^ The importance of this process is illustrated by distal lung alveolar epithelial progenitors, which differentiate into ATI and II cells in organoids. These require mesenchymal support cells and only form if these mesenchymal cells are in contact with epithelial progenitors.^9,17^ Additionally, we have previously demonstrated that lung fibroblast-derived secretome ameliorated lung tissue injury.^20^ Exposure of these mesenchymal cells to cytokines and factors (e.g., transforming growth factor (TGF)-β, cigarette smoke (CS)) that drive the disease process in COPD strongly impairs their promotive role.^17,18,21^

Lung fibroblasts support alveolar epithelial regeneration through several mechanisms, including paracrine signaling, by releasing extracellular vesicles (EVs) and soluble factors.^22^ Here, we aimed to identify factors in the lung fibroblast secretome with regenerative potential for therapeutic development. Using a proteomics-guided drug discovery strategy, we identified osteoglycin (OGN) and its active fragment to have significant potential in supporting alveolar epithelial repair. We also demonstrate reduced expression of OGN in lung tissue from current smokers and patients with smoking-associated COPD. Additionally, we demonstrate that treatment with OGN and its active fragment ameliorates elastase-induced injury in precision-cut lung slices (PCLS) and restores lung function in mice.

## MATERIALS AND METHODS

Full details of materials and methods are described in the online supplementary material.

### Animals

Lung harvesting from mice for organoid experiments and PCLS was performed at the Central Animal Facility (CDP) of the University Medical Center Groningen (UMCG) per the national guidelines and upon approval of the experimental procedures by CDP and the Institutional Animal Care and Use Committee (IACUC) of the University of Groningen, under AVD1050020209205. Animals were housed conventionally under a 12-h light-dark cycle and received food and water ad libitum.

### Human material

Human lung tissue was obtained from lung transplant donors in strict adherence to the Research Code of the UMCG, as stated on https://umcgresearch.org/w/research-code-umcg as well as national ethical and professional guidelines Code of Conduct for Health Research (https://www.coreon.org/wp-content/uploads/2023/06/Code-of-Conduct-for-Health-Research-2022.pdf). The use of left-over lung tissue in this study was not subject to the Medical Research Human Subjects Act in the Netherlands, as confirmed by a statement of the Medical Ethical Committee of the UMCG, and therefore exempt from consent according to national laws (Dutch laws: Medical Treatment Agreement Act (WGBO) art 458 / GDPR art 9/ UAVG art 24). Human lung tissue was acquired from extra tissue left over after lung surgeries, which exceed the amount needed for clinical care purposes. All samples and clinical information were blinded. before experiments.

#### Primary alveolar epithelial isolation and organoid culture

Isolation of murine primary alveolar epithelial cells, in brief, Epcam^+^ cells (CD31^-^/CD45^-^/CD326^+^), was based on previously published protocols.^3^ We have used a total of 54 mice (male and female (1:1), eight to fourteen weeks old). For human organoids, adult human donor tissue was isolated from histologically normal regions of lung tissue specimens obtained at the UMCG from N = 11 patients with GOLD stage IV COPD (see Table S7 for clinical information of donors). For organoid cultures, freshly isolated Epcam^+^ / EpCAM^+^ cells were combined with lung fibroblasts at a 1:1 ratio (10,000 cells each) in DMEM/F12 containing 10% (v/v) FBS. The cell suspension was then diluted 1:1.5 (v/v) with Corning® Matrigel® Membrane Matrix (Corning, 356234) and was then seeded into transwell inserts (Greiner, 662641) in 24-well plates (100 µL/insert). Organoid cultures were cultured at 37 °C with 5% CO_2_. The medium was refreshed every 2-3 days. The total number and diameters of organoid structures (>50 µm) was measured on day 14 using NIS-Elements with bright field microscopy (20x magnification).

#### Organoid resorting to regain fibroblasts and epithelial cells

For organoid resorting, a mixture of 300,000 Epcam^+^ cells and 300.000 CCL206 fibroblasts was seeded in a 1 mL solution of Matrigel diluted 1:1.5 (v/v) with DMEM/F12 (supplemented with 10% FBS) and added to one well of a 6-well plate. After the Matrigel solidified for an hour, 2 mL of organoid culture medium was added on top of the Matrigel, including EVs (10^9^ EVs/mL), SFs (30 µg/mL), or OGN (10 µg/mL). After three days, dispase (Corning, 354235) was added to each well for 30 mins at 37 °C to dissociate the Matrigel. MACS buffer (MACS rinsing solution (Milteny Biotec, 130-091-222) premixed with BSA (Milteny Biotech, 130-091-376)) was added to stop the dispase activity. Organoids were collected and centrifuged at 300 g for 5 mins. The pellets were resuspended in 5 mL diluted trypsin (1:5 in PBS, v/v) (T7409, Sigma-Aldrich) for 5 mins at 37 °C, after which 9 mL DMEM/F-12 supplemented with 10% FBS was added to neutralize trypsin action. The cell pellets were incubated with CD326 microbeads for 20 mins and resuspended in MACS buffer. The cell suspensions were introduced to the QuadroMACS™ Separator system to obtain CD326^-^ fibroblasts and CD326+ (Epcam^+^) epithelial cells derived from organoids, which were then used for further experimentation.

#### Proteomic analysis

Proteomic data from MRC5-derived EVs and SFs were obtained from 4 biological replicates. Liquid chromatography-tandem mass spectrometry (LC-MS) was employed for unbiased shotgun proteomic analysis. In-gel digestion was performed as described previously^23^ using 1 µg total protein in the EV-enriched samples and 75 µg total protein for the SF-enriched samples. LC-MS-based proteomics analyses were performed as described previously^24^ using 0.5 µg digested total protein starting material from the EVs or 1 µg digested total protein starting material from the SFs. LC-MS raw data were processed with Spectronaut (version 15.4.210913, Biognosys) using the standard settings of the directDIA workflow with a human SwissProt database (www.uniprot.org, 20350 entries). For the quantification, local normalization was applied, and the Q-value filtering was adjusted to the classical setting without imputing.

#### Immunohistochemical staining on human lung slides

Control and COPD human lung tissue was obtained from leftover material at the UMCG and St. Mary’s Hospital, Mayo Clinic Rochester, MN. This staining was part of the HOLLAND (HistopathOlogy of Lung Aging aNd COPD) cohort.^25^ Immunohistochemical (IHC) staining on human lung slides was performed as described previously.^25^

#### Precision-cut lung slices

PCLS were obtained from naive C57BL/6J mice (eight to fourteen weeks old) and prepared as described previously.^20^ A total of twelve mice (male and female, ratio 1:1) were used for PCLS experiments. Mouse lungs were filled with 1.5% low-melting point agarose (Gerbu Biotechnik GmbH, Wieblingen, Germany) solution. Subsequently, lung lobes were cut into precision-cut lung slices (PCLS) of 250 µm in thickness. To induce emphysematous changes, matched slices were treated with 2.5 µg/mL elastase for 16h. Slices were treated with 10 µg/mL OGN or 4.48 µg/mL OGN active fragment for 40h, overlapping with the 16h elastase treatment.

#### Elastase in vivo model

Pulmonary emphysematous changes were induced by intratracheal instillation of pancreatic porcine elastase (40 U/kg body weight) in 40 µL sterile PBS on day zero, as previously described.^20^ Animals were treated every other day from day zero till day nine (5 treatments in total) with OGN fragment (6.75 µg or 20.25 µg), and sacrificed on day ten.

#### Statistical analysis

Results are shown as mean ± standard error, with sample size and repeats in figure legends. Four normality tests (D’Agostino-Pearson, Anderson-Darling, Shapiro-Wilk, Kolmogorov-Smirnov) were conducted; if three indicated normality, parametric tests were applied. One-Way ANOVA with Dunnett’s multiple comparisons was used for group comparisons, and unpaired/paired two-tailed Student’s t-test or Mann-Whitney and Wilcoxon tests for two-group comparisons. Significance was set at p < 0.05. Analyses were done in GraphPad Prism 10 and RNA sequencing analysis in R Studio.

## RESULTS

### Lung fibroblast-derived EVs and SFs support alveolar epithelial growth

To assess the regenerative potential of the secretome of lung fibroblasts, we used an established *in vitro* murine lung organoid model.^3^ We assessed whether fibroblasts exert their preventive function in a paracrine manner through the secretion of extracellular vesicles and soluble factors (Fig. 1a). MRC5 human lung fibroblast-derived EV-enriched fractions (EVs) and SFs-enriched fractions (SFs) were purified using a combination of ultrafiltration and size exclusion chromatography, and previously characterized according to MISEV guidelines.^20,26^ Morphological assessment of EV-enriched fractions revealed intact, spherical vesicles with a diameter of around 110 nm (Fig. S1). The colony forming efficiency (CFE) of organoids established by coculturing murine lung tissue-derived CD31^-^/CD45^-^/Epcam^+^ cells and CCL-206 fibroblasts was evaluated. Freshly isolated CD31^-^/CD45^-^/Epcam^+^ cells are mainly composed of AT2 cells (around 80-85%) as we have demonstrated previously using FACS and RNAseq analysis.^17,27,28^ The CFE was significantly increased by treating cultures with EVs (10^9^ particles/mL, 50,000 particles per cell) or SFs (30 µg/mL) (Fig. 1a-c), without affecting organoid size (Fig. 1d). In addition, EV or SF treatment increased organoid formation in a concentration-dependent manner (Fig. S2a-d). Interestingly, reducing the number of fibroblasts yielded significantly fewer murine organoids, underscoring the supportive function of fibroblasts in epithelial organoid formation (Fig. S2e-f). We compared treatment effects with EVs or SFs in the presence of varying fibroblast numbers: 10,000, 5,000, or 2,500 CCL206 fibroblasts (Fig. S2g-j). As the number of fibroblasts decreased, both treatments with EVs (10^9^ particles/mL) or SFs (30 µg/mL) exhibited a more pronounced increase in CFE (Fig S1g-i). To specifically analyze the impact of treatment with EVs or SFs on organoids derived from alveolar epithelial progenitors, we used immunofluorescence staining to distinguish between airway-type organoids (ACT^+^) and alveolar-type organoids (SPC^+^). This analysis revealed that alveolar epithelial progenitors were more prone to induce CFE to treatment with EVs or SFs compared to control, yielding significantly more SPC^+^ organoids (Fig. 1e) without affecting the number of ACT^+^ organoids (Fig. S2k).

**Fig. 1.**
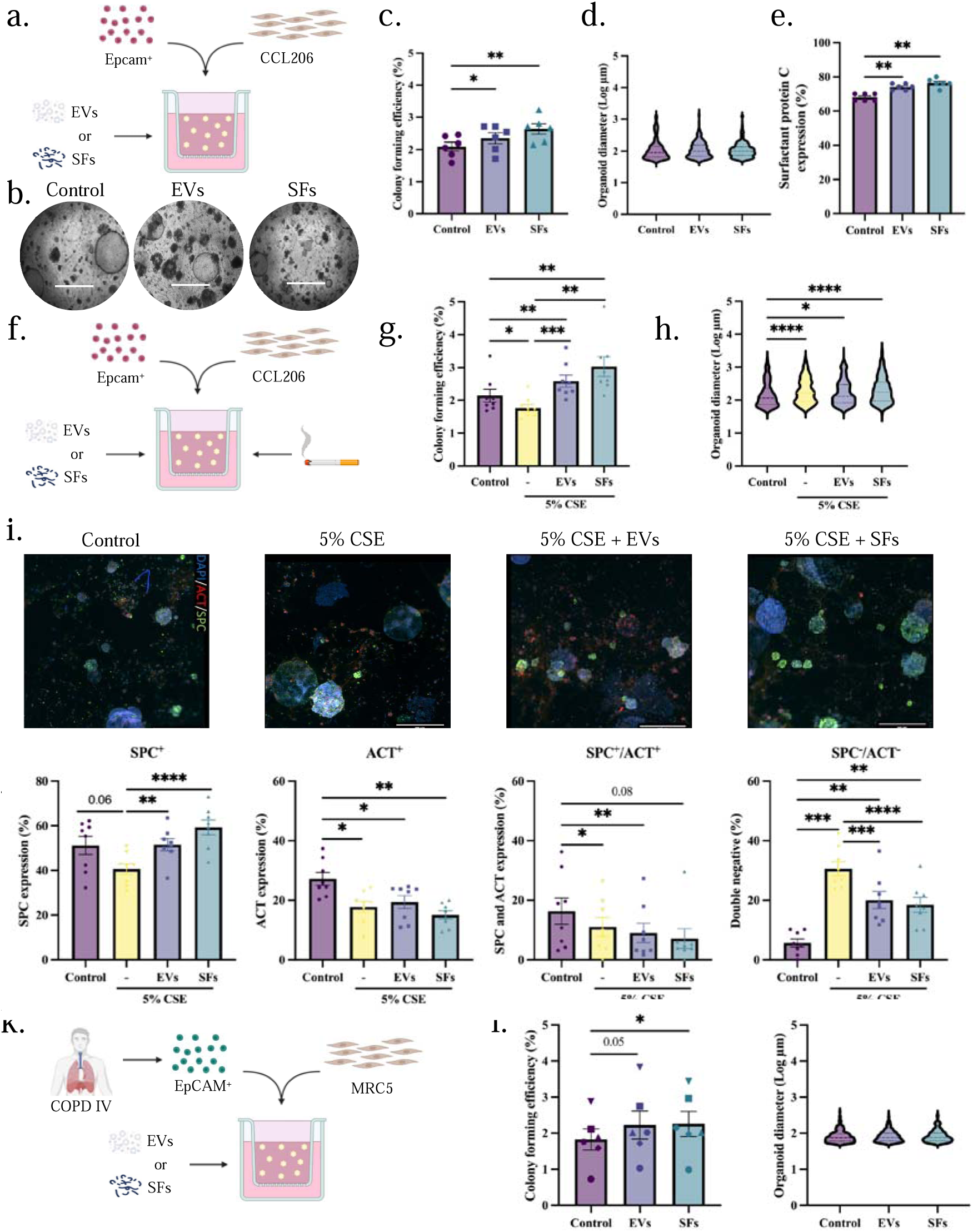
Lung fibroblast-derived EVs and SFs support alveolar organoid formation. **a** Schematic of in vitro murine organoid design treated with EVs or SFs. **b** Brightfield images of murine lung organoids on day 14 (scale = 700 µm). **c** CFE of murine organoids (mean ± SEM, N=6, one-way ANOVA, Dunnett test). **d** Log of murine organoid diameter (median, N=6, Kolmogorov-Smirnov test, α = 0.0025). **e** Immunohistochemistry quantification for SPC in murine organoids (mean ± SEM, N=6, one-way ANOVA, Dunnett test). **f** Schematic of in vitro CSE-exposed murine organoid design. **g** CFE of CSE-exposed murine organoids (mean ± SEM, N=8, one-way ANOVA, Dunnett test). **h** Log of CSE-exposed murine organoid diameter (median, N=8, Kolmogorov-Smirnov test, α = 0.0167). **i** Immunofluorescence images of organoids for airway-type (ACT, red), alveolar-type (SPC, green), and DAPI (nuclei, blue) (scale = 500 µm). **j** Immunohistochemistry quantification for SPC^+^, ACT^+^, SPC^+^/ACT^+^, and SPC^-^/ACT^-^ organoids (mean ± SEM, N=6, one-way ANOVA, Dunnett test). **k** Schematic of in vitro human organoid design. **l** CFE of human organoids (mean ± SEM, N=6, one-way ANOVA, Dunnett test). **m** Log of human organoid diameter (median, N=6, Kolmogorov-Smirnov test, α = 0.025). Statistically significant comparisons: *p < 0.05, **p < 0.01, ***p < 0.001, ****p < 0.0001.

During COPD development, the alveolar epithelial cells are exposed to a hostile microenvironment, which changes their composition, phenotype, and responses to molecules. To simulate this hostile microenvironment in our organoid model, we exposed organoid cultures to 5% CS extract (CSE; Fig. 1f), which yielded, in accordance with previous findings, a significantly lower organoid number (Fig. 1g) and increased organoid diameter (Fig. 1h).^3^ Concurrent treatment with either EVs or SFs was able to counteract the negative effects of CSE in respect of CFE number. In contrast, organoid size was increased compared to control (Fig. 1g-h). Immunofluorescence staining revealed that the number of SPC^+^ and ACT^+^ organoids decreased in the presence of 5% CSE (Fig. 1i-j), and the proportion of organoids negative for SPC and ACT increased (Fig. 1j). Treatment with EVs and SFs, however, increased the number of SPC^+^ organoids (Fig. 1j) and lowered the number of double-negative (SPC^-^/ACT^-^) (Fig. 1j). For a translational perspective, we further investigated lung fibroblast-derived EVs and SFs on human organoid formation derived from EpCAM^+^ epithelial cells isolated from COPD IV patient lung tissue (Fig. 1k). EV and SF treatment increased the CFE of human organoids, whereas diameter was unaffected by both treatments (Fig. 1l-m).

#### A proteomics-guided drug discovery strategy identifies osteoglycin to induce organoid formation

We used a proteomics-guided drug discovery strategy (Fig. 2a) to identify potential therapeutic factors within the fibroblast secretome that may restore defective lung repair. Analysis of lung fibroblast-derived EVs and SFs (N=4) revealed 1262 proteins and 2090 proteins that were identified in at least one of the replicates, respectively. To demonstrate the nature of the EVs, we analyzed the presence of proteins stipulated following the MISEV guidelines.^26^ EVs showed enrichment of transmembrane and GPI-anchored proteins associated with EVs, in contrast to the relatively low abundance in SFs (Table S1). Additionally, CD90, a marker for mesenchymal stromal cells, was present in all EV samples. In summary, the proteomic analysis collectively demonstrates an enrichment of EV-associated proteins in EVs when compared to SFs.

**Fig. 2.**
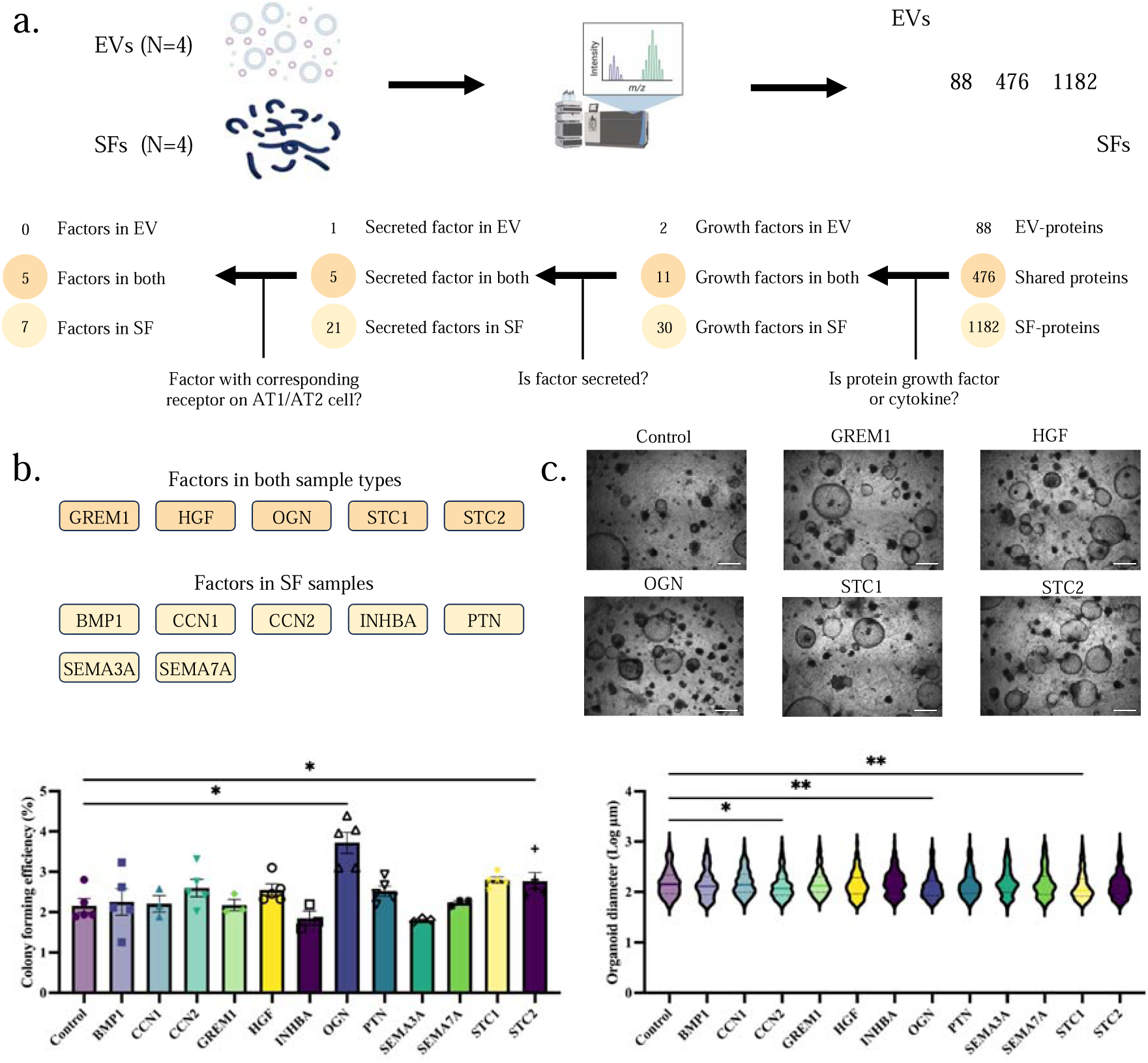
A proteomics-guided drug discovery strategy identifies osteoglycin to induce organoid formation. **a** Schematic outline of the proteomics-guided drug discovery strategy using human lung fibroblasts (MRC5). **b** Overview of included factors in the secretome of human lung fibroblasts. **c** Representative brightfield image of murine organoid cultures for proteomics-guided drug discovery strategy with factors in both EVs and SFs. **d** CFE of murine organoids in proteomics-guided drug discovery strategy (mean ± SEM, N=3-5, paired one-way ANOVA, Dunnett test). **e** Log of murine organoid diameter in proteomics-guided drug discovery strategy (median, N=3-5, Kolmogorov-Smirnov test, Bonferroni correction: α = 0.004). Statistically significant comparisons: *p < 0.05, **p < 0.01.

For subsequent analysis, we focused on 564 EV-proteins and 1658 SF-proteins that were consistently present in all four replicates (Fig. 2a). Notably, 476 of these proteins were found in both the EVs and SFs (Fig. 2a, Table S2). Since growth factors and cytokines are known to affect epithelial progenitor cell behavior strongly, we analyzed the number of growth factors and cytokines using the Gene Set Enrichment Analysis tool (www.gsea-msigdb.org) (Fig. 2a). We identified a total of 43 growth factors, of which two factors were exclusively present in EVs, 30 factors in SFs, and 11 factors in both (Fig. 2a). Since our focus was on paracrine interactions, we narrowed down the number of candidates based on predictions for secretion of the factors using Signal IP (https://services.healthtech.dtu.dk/services/SignalP-6.0/) and Phobius (https://phobius.sbc.su.se/), resulting in a total of 32 secreted factors (Fig. 2a, Table S3). Lastly, we investigated whether the secreted factor interacts with a receptor expressed in the target recipient cells (AT1 and AT2 cells) (Fig. 2a). In total, we identified twelve factors for evaluation in lung organoids: seven factors within the SF-protein group and five factors present in both sample types (Fig. 2a-b, Table S3).

Using murine lung organoid cultures, we assessed the efficacy of these factors in inducing organoid formation (Fig. 2c-d). All information about the recombinant factors used in this study is provided in Table S4. Stanniocalcin (STC)-2 increased organoid numbers without affecting size, while bone morphogenetic (BMP)1, hepatocyte growth factor (HGF), pleiotrophin (PTN), and STC1 tended to increase CFE. CCN2 and STC1 significantly reduced organoid size (Fig. 2e). Notably, osteoglycin (OGN) emerged as the most promising factor identified in the secretome of lung fibroblasts, inducing additional organoid formation by approximately 75% compared to controls (Fig. 2c-d). Interestingly, OGN was present in both EVs and SFs (Fig. 2b). Given our previous observations of the supportive effects of both EVs and SFs on alveolar organoid formation (Fig. 1), we decided to further explore the potential of OGN as a therapeutic candidate for lung repair.

#### Osteoglycin expression is reduced in response to cigarette smoke

To assess cell type-specific expression of OGN in the lungs, we performed single-cell RNA sequencing (scRNA-Seq) analysis of naive murine lung tissue, which showed predominant OGN expression in fibroblasts and, to a lesser extent, in mesothelial cells (Fig. 3a). Confirming these results in human lung tissue, analysis of a public scRNA-Seq dataset (https://lungmap.nl/) showed that 59.06% of alveolar fibroblasts (type 2) express OGN (Fig. 3b, Fig. S3a). Furthermore, OGN gene expression was observed in both MRC5 cells and in primary human lung fibroblasts (Fig. S3b). These data suggest alveolar fibroblasts as the primary endogenous source of OGN in the lungs. In human lung tissue from never-smoker controls, OGN staining was most pronounced around larger arteries, interlobular septa, and pleura, correlating with areas rich in high collagen. (Fig. 3c, Fig. S4a). This suggests that fibroblasts, known to secrete important components of the extracellular matrix, such as collagens, are a likely endogenous source for OGN.

**Fig. 3.**
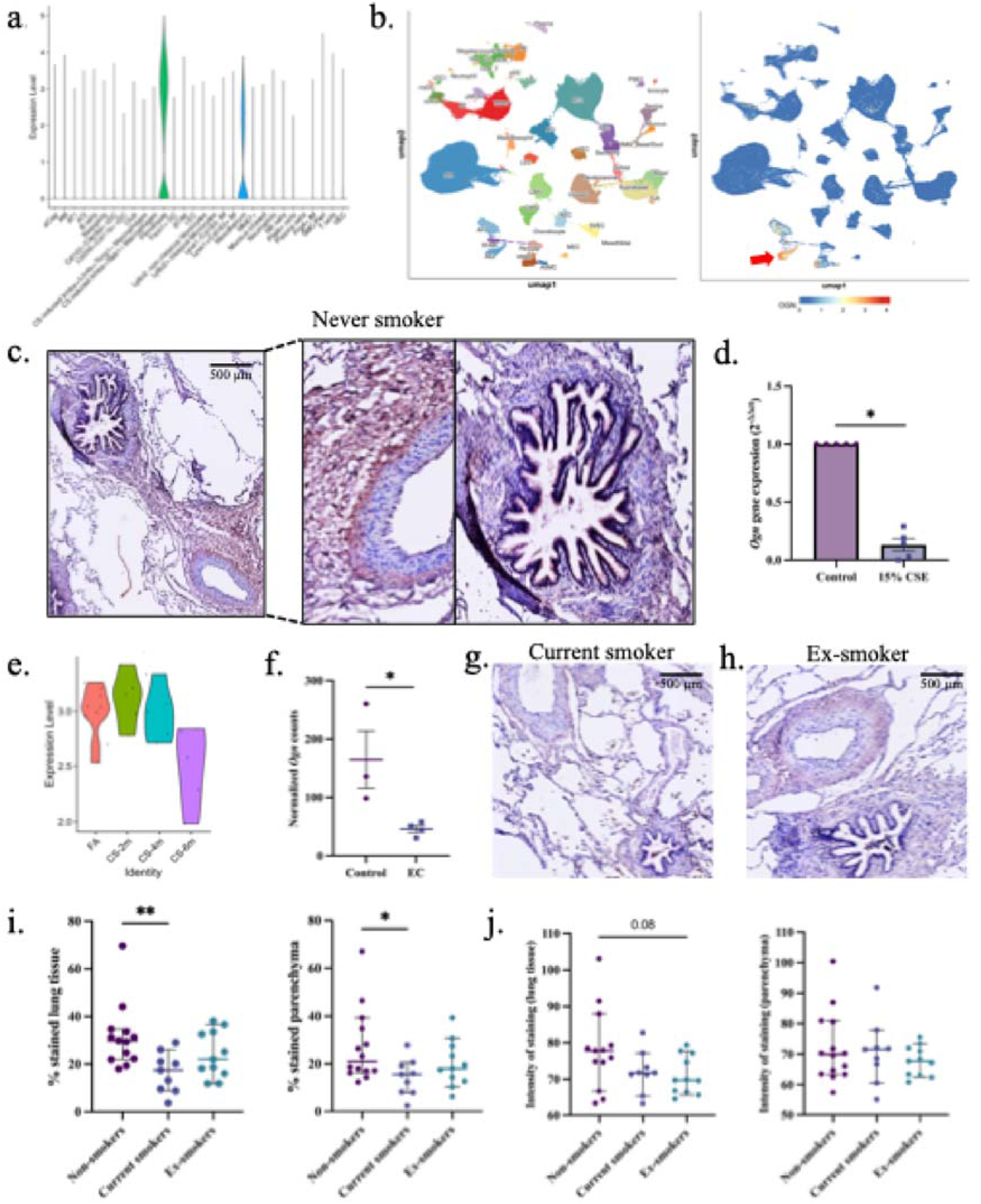
Cigarette smoke reduces fibroblast-derived osteoglycin expression in the lungs. **a** ScRNA-Seq analysis of *Ogn* expression by cell type in murine lungs. **b** ScRNA-Seq analysis showing cell type specificity of *OGN* in the human lung, with red arrow indicating OGN in alveolar fibroblasts. **c** Image of OGN staining in lung tissue of never-smoker (scale = 500 µm), with close-up showing an artery and an airway. **d** *Ogn* gene expression in PCLS treated with 15% CSE (mean ± SEM, N=5, paired t-test on log-transformed ΔCt values). **e** Pseudobulk ScRNA-Seq of *Ogn* in fibroblasts from lung tissue of mice exposed to two, four, and six months of CS compared to filtered air (FA). **f** Normalized counts from DESeq2 analysis of *Ogn* in fibroblasts upon treatment with a COPD-related exacerbation cocktail (mean ± SEM, N=3-4). **g-h** OGN staining images in lung tissue of current smoker and ex-smoker donor (scale = 500 µm). **i** Percentage (%) of positively stained area for OGN in whole lung tissue and parenchyma in never, current, and ex-smokers. **j** Intensity of positive staining for OGN in whole lung tissue and parenchyma of never, current, and ex-smokers. Statistically significant comparisons: *p < 0.05, **p < 0.01.

We then investigated the impact of CS on *Ogn* expression by exposing murine PCLS to 15% CSE for 24 hours and analyzed *Ogn* expression. CSE treatment significantly reduced *Ogn* expression in PCLS (Fig. 3d). Additionally, in vivo mice exposed to CS for two, four, and six months^29^ showed progressively diminished *Ogn* expression in lung tissue, as revealed by scRNA-Seq analysis (Fig. 3e). Furthermore, *Ogn* was significantly downregulated in murine fibroblasts from resorted organoids treated for 72 hours with an exacerbation cocktail (EC) containing COPD-related cytokines (300 pg/mL IL-1β, 300 pg/mL IL-6, 20 ng/mL keratinocyte chemoattractant, and 200 pg/mL TNFα) (Fig. 3f).^27^

To study the impact of smoking on OGN expression, we compared OGN protein expression in lung tissue of never, current, and ex-smokers (Fig. 3g-h, Table S5a). OGN was more abundant in never-smoker lung tissue compared to current and ex-smoker tissue (Fig. 3g-i). Image analysis showed a lower percentage area of OGN-positive staining in current smokers compared to never smokers (Fig. 3i). Interestingly, OGN expression in ex-smokers was not different from current smokers or never smokers, with mean expression values intermediate between these groups, implying a persistent impact of cigarette smoke on OGN expression (Fig. 3i). OGN staining intensity showed no significant differences between never, current, and ex-smokers (Fig. 3j). Next, we assessed whether OGN expression changed in patients with moderate-severe COPD (II/III) or severe-early onset (SEO-)COPD compared to ex-smoker control lung tissue (Fig. S4b-c). In Table S5B, the clinical parameters of donors included in this analysis are summarized. In patients with COPD II/III, no significant differences were observed in the percentage area and the average intensity of the OGN staining in both whole lung and parenchyma (Fig. S4d-e). Interestingly, we did observe a tendency for a lower percentage area of OGN staining and higher average staining intensity in the parenchyma of SEO-COPD compared to the control (p=0.09) (Fig. S4f-g). These data suggest that OGN expression is mainly affected by current smoking, corroborating our findings in a CS-exposed mouse model.

#### Osteoglycin promotes alveolar epithelial growth

OGN, also known as mimecan, is an endogenous small leucine-rich proteoglycan (SLRP) with diverse roles in human biology, including tissue development and fibrosis regulation.^30^ We investigated the effects of OGN on alveolar epithelial progenitor cell activation using murine lung organoids. Human recombinant OGN protein provided concentration-dependent growth for lung organoids (Fig. 4a-b) without altering organoid size (Fig. 4c). Furthermore, there was an enhanced number of SPC^+^ organoids with OGN treatment (Fig. 4d). In contrast, the proportion of ACT^+^ organoids remained unchanged (Fig. S5a).

**Fig. 4.**
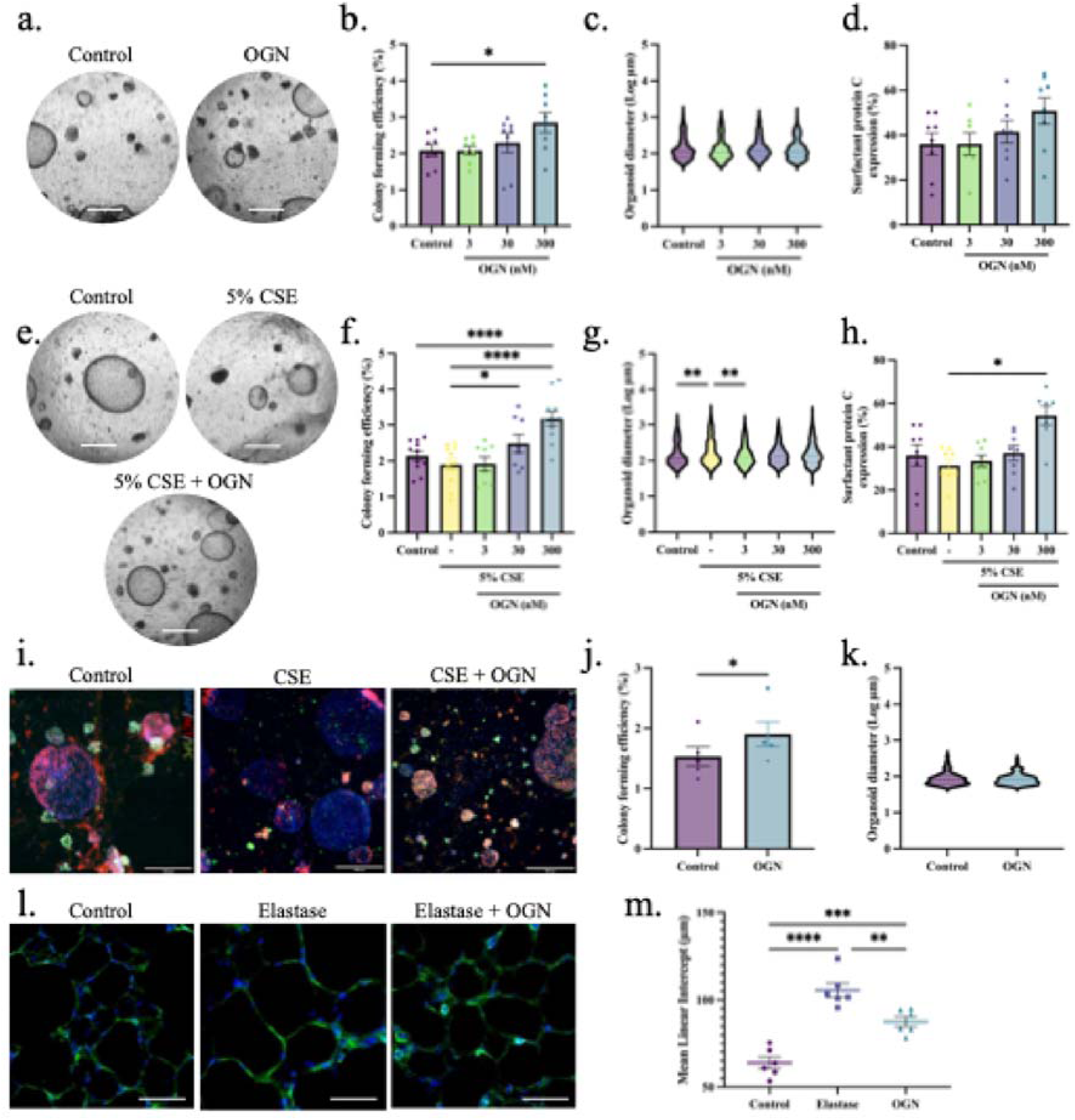
Osteoglycin supports alveolar organoid formation. **a** Brightfield images of murine lung organoids with OGN (300 nM) (scale = 500 µm). **b** CFE of murine organoids (mean ± SEM, N=8, paired one-way ANOVA, Dunnett test). **c** Log of murine organoid diameter (median, N=8, Kolmogorov-Smirnov test, Bonferroni correction: α = 0.0167). **d** Immunohistochemistry for pro-surfactant protein C in murine organoids on day 14 (mean ± SEM, N=6, paired one-way ANOVA, Dunnett test). **e** Brightfield images of CSE-exposed murine lung organoids with OGN (300 nM) (scale = 500 µm). **f** CFE of CSE-exposed murine organoids (mean ± SEM, N=8-11, paired one-way ANOVA, Dunnett test). **g** Log of CSE-exposed murine organoid diameter (median, N=8-11, Kolmogorov-Smirnov test, Bonferroni correction: α = 0.0125). **h** Immunohistochemistry for SPC in CSE-exposed murine organoids (mean ± SEM, N=7-8, paired one-way ANOVA, Dunnett test). **i** Immunofluorescence images of organoids for airway-type (ACT, red), alveolar-type (SPC, green), and DAPI (blue) (scale = 500 µm). OGN is 300 nM. **j** CFE of human organoids with 300 nM OGN (mean ± SEM, N=5, paired Student T-test). **k** Log of human organoid diameter with 300 nM OGN (median, N=5, Kolmogorov-Smirnov test). **l** PCLS stained for F-actin (green) and DAPI (blue) (scale = 50 µm) after elastase and OGN (300 nM). **m** LMI measurements following treatments (mean ± SEM, N=6, one-way ANOVA, Dunnett test). Significant comparisons: *p < 0.05, **p < 0.01, ***p < 0.001, ****p < 0.0001.

In addition, we exposed the organoid cultures to 5% CSE. OGN maintained its regenerative effects without affecting the organoid size and increased the proportion of SPC^+^ organoids in a concentration-dependent manner (Fig. 4e-h). As illustrated in Fig. S5b, CSE treatment alone increased the proportion of double negative organoids (SPC^-^/ACT^-^). In contrast, additional OGN treatment increased the percentage of alveolar (SPC^+^) organoids, which resulted in a decreased percentage of airway-type (ACT^+^) or double-positive organoids (SPC^+^/ACT^+^) (Fig. 4i). Significantly, in a real disease model, using human COPD IV organoids, OGN treatment increased the CFE number without affecting diameter (Fig. 4j-k).

To confirm the effects of OGN on alveolar epithelial growth, we used a mouse PCLS experiment in which we induced emphysematous changes with elastase enzyme.^20^ After 16h incubation with elastase, parenchymal lung tissue damage indicative of emphysema was induced, as evidenced by a significant increase in mean linear intercept (LMI) (Fig. 4l-m). Interestingly, treatment with OGN for 40h, of which the first 16h were concomitant with elastase exposure, effectively prevented the induction of lung tissue injury (Fig. 4l-m). Taken together, these data demonstrate the profound potential of OGN on the human and mouse alveolar epithelium, positioning it as a promising therapeutic candidate supporting alveolar epithelial repair.

#### Osteoglycin enhances epithelial ROS defense mechanisms and increases fibroblast signaling

To determine the potential mechanism of action, we first studied transcriptomic changes in murine lung organoids treated 72 hours with EVs, SFs, or OGN and subsequently, re-sorted into epithelial cells and fibroblasts for bulk RNA sequencing (Fig. 5a). This time point was chosen to capture transcriptomic changes during the initiation of organoid formation. Principal component analysis (PCA) showed distinct transcriptional profiles for the two cell populations (Fig. 5b). Additionally, OGN-treated Epcam^+^ cells and fibroblasts were transcriptionally different from the other conditions. To confirm cell purity, we compared the proportion of AT2 cells and fibroblasts in all untreated samples based on a cell marker gene signature (Table S6). The gene signature for AT2 cells was significantly higher in the epithelial cell population, while cells with a fibroblast signature were enriched in the fibroblast population (Fig. S6a).

**Fig. 5.**
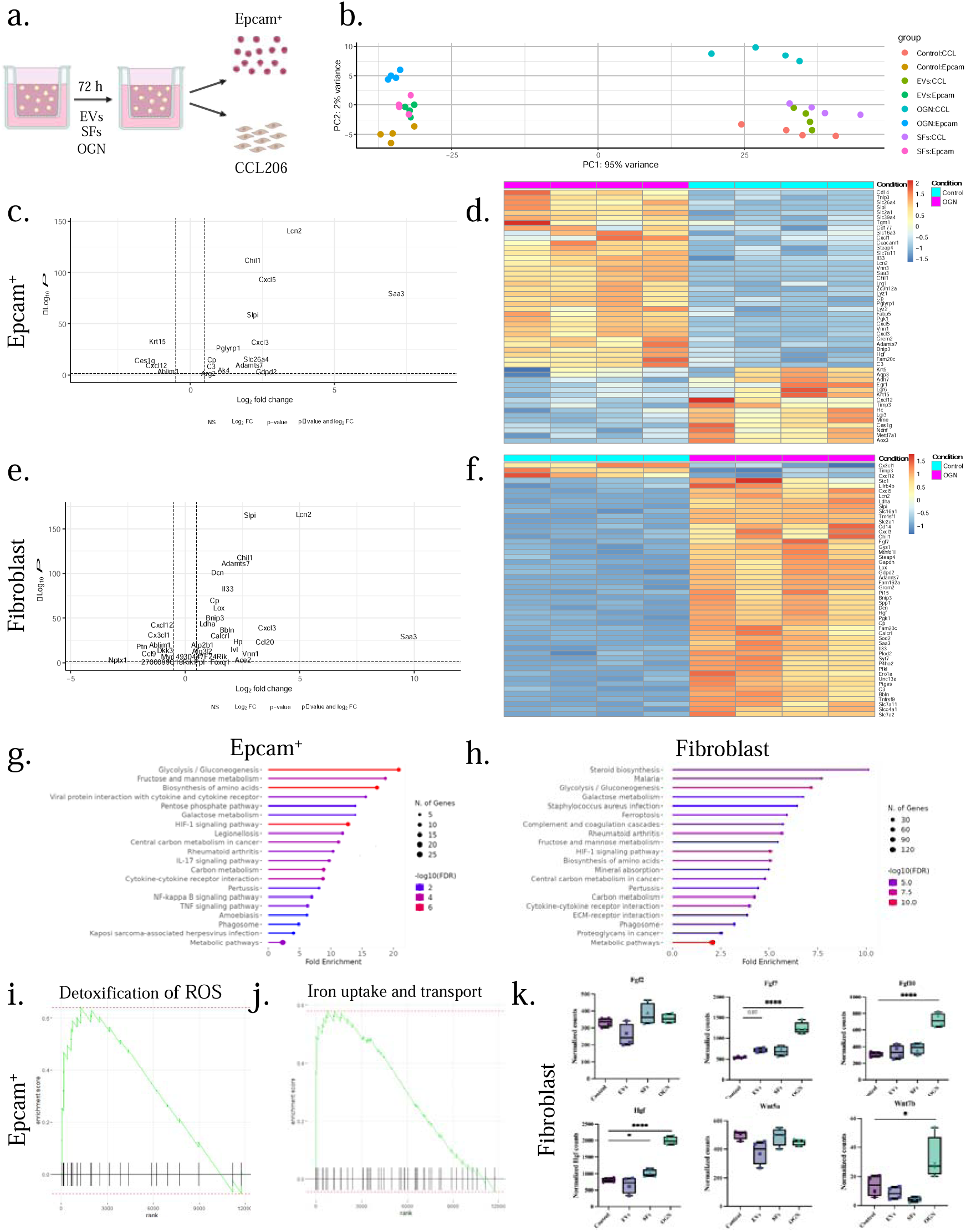
Osteoglycin enhances gene expression for epithelial cell ROS defense mechanisms and increases growth factor expression in fibroblasts. **a** Schematic of organoid RNA sequencing experimental design. **b** PCA plot of epithelial progenitor cells and fibroblasts from murine organoids treated with vehicle, EVs, SFs, or OGN for 72 hours (N=4). **c** Volcano plot illustrating the response to OGN versus control in epithelial progenitor cells (padj<0.05 and logFold>1). **d** Heatmap of top 50 significant genes up- or downregulated in OGN-treated epithelial progenitor cells; shown as z-scores. **e** Volcano plot illustrating the response to OGN versus control in fibroblasts (padj<0.05 and logFold>1). **f** Heatmap of top 50 significant genes up- or downregulated in OGN-treated fibroblasts; shown as z-scores. **g** Top 20 upregulated pathways in epithelial progenitors from organoids treated with OGN. **h** Top 20 upregulated pathways in OGN-treated fibroblasts. **i-j** Reactome enrichment score for “detoxification of ROS” and “iron uptake and transport” in epithelial progenitor cells. **k** Normalized counts of Fgf2, Fgf7, Fgf10, Hgf, Wnt5a, and Wnt7b in fibroblasts with EVs, SFs, or OGN (mean ± min/max, N=4). Statistically significant comparisons: *p < 0.05 and ****p < 0.0001.

Paired differential expression analysis using DESeq2 revealed that EV treatment significantly upregulated 67 genes and downregulated 44 genes in epithelial cells and upregulated 1186 genes and downregulated 983 genes in fibroblasts (Fig. S6b-e). SF treatment had moderate effects, with 27 genes upregulated and 29 downregulated in epithelial cells and 95 genes upregulated and 47 downregulated in fibroblasts (Fig. S6f-i). OGN treatment had more profound effects, with 79 genes downregulated, 118 genes upregulated in epithelial cells, 489 genes downregulated, and 652 upregulated in fibroblasts (Fig. 5c-f). Overall, EVs, SFs, and OGN had the most pronounced transcriptomic effect in fibroblasts compared to epithelial cells.

We then performed GSEA to decipher the observed transcriptomic changes in epithelial cells. Using the KEGG reference database, we identified pathways overrepresented within OGN-modulated genes in Epcam^+^ cells (Fig. 5g) and fibroblasts (Fig. 5h). We also analyzed pathways affected by EVs or SFs (Fig. S7a-c) or OGN (Fig. S7d). OGN treatment upregulated pathways related to Glycolysis/Gluconeogenesis, Biosynthesis of amino acids, HIF-1 signaling, Carbon metabolism, and Metabolic pathways in both cell populations. Reactome analysis showed increased expression of protective pathways, including *Detoxification of reactive oxygen species* and *Iron uptake and transport* in OGN-treated epithelial cells (Fig. 5i-j). Furthermore, OGN treatment enhanced the expression of antiporter subunits *Slc3a2* and *Slc7a11*, facilitating cystine uptake for glutathione synthesis, acting as a co-factor for glutathione peroxidase (Fig. S8a-b). Additionally, OGN increased the expression of superoxide dismutase 2 (*Sod2*), clearing mitochondrial reactive oxygen species (Fig. S8c), while the expression of senescence marker p21 (Cdkn1a) tended to decrease (Fig. S8d). Analysis of AT1 and AT2 markers revealed that the AT1 markers *Emp2*, *Lmo7*, and *Gprc5a* remained unchanged after treatment with EVs, SFs or OGN, and also a composite AT1 gene signature based on the top 10 AT1 marker genes^31^ was unaffected (Fig. S9a). AT2 markers showed a disparate pattern with some (*Sftpb* and *Sftpc*) being unchanged and others (*Sftpa1*) being increased following OGN (but not EVs or SFs) exposure (Fig. S9b). The AT2 signature based on the top 10 AT2 marker genes^31^ was unaffected by OGN (Fig. S9b).

As the transcriptomic changes were most pronounced in fibroblasts, we focused on their expression of growth factors known to be essential mitogens for AT2 cells, including fibroblast growth factor (FGF)2, FGF7, FGF10, HGF, and WNT ligands. We focused on their expression in fibroblasts treated with EVs, SFs, and OGN. OGN notably increased Fgf7 and Fgf10 expression in fibroblasts, with a trend towards upregulation with EV treatment (Fig. 5k). HGF expression significantly increased with SFs and OGN treatment. OGN treatment also increased Wnt7b expression and showed a tendency towards decreased Wnt5a expression (Fig. 5k). Together, these findings suggest enhanced epithelial-fibroblast crosstalk mediated by growth factors upon OGN treatment, potentially contributing to the increased organoid formation.

#### An active fragment of osteoglycin supports alveolar epithelial regeneration

The protein structure of OGN consists of seven leucine-rich repeats (LRRs). LRRs four to seven contain positively charged residues and are part of the C-terminal chain, forming a tail-like structure composed of amino acids 180 to 298.^30^ The specific structure and charge in this protein fragment suggest a role in biological activity or its signaling (Fig. 6a).^30^ Hence, we tested the effects of this OGN fragment on alveolar epithelial progenitor behavior. Similar to full-length OGN, the fragment, which was obtained commercially, increased the CFE of murine organoids in a concentration-dependent manner while organoid size remained unaltered (Fig. S10a and Fig. 6b-c). Higher concentrations of fragment tended to increase the proportion of alveolar (SPC^+^) organoids (Fig. S10b-c) and reduced double positive (SPC^+^ ACT^+^) organoids (Fig. S10c). In the presence of 5% CSE, the fragment increased the CFE and organoid diameter (Fig. 6d-e). Additionally, the fragment significantly increased the proportion of alveolar (SPC^+^) organoids (Fig. S10d) and reduced the proportion of ACT^+^ and double-positive organoids (Fig. S10d). Our next aim was to validate the effect of OGN and its active fragment on the growth of AT2 cells within the organoid model. To achieve this, we enriched the initial cell population for AT2 cells by selecting for major histocompatibility complex class II (MHCII+) cells and then compared the CFE of the MHCII^+^ cells to the broader Epcam^+^ population (Fig. S11a).^27^ Treatment with OGN or its active fragment resulted in a significant increase in the number of organoids formed in both the MHCII^+^ and Epcam^+^ cell populations to a similar extent (Fig. S11b). Notably, in MHCII^+^ cultures, treatment with the OGN fragment led to a significant increase in the formation of larger-diameter organoids (Fig. S11c). Additionally, in human organoid cultures derived from COPD IV donors, the CFE of organoids was significantly increased upon treatment with OGN fragment. In summary, these results indicate that a smaller active fragment of OGN is sufficient to support alveolar epithelial progenitor behavior.

**Fig. 6.**
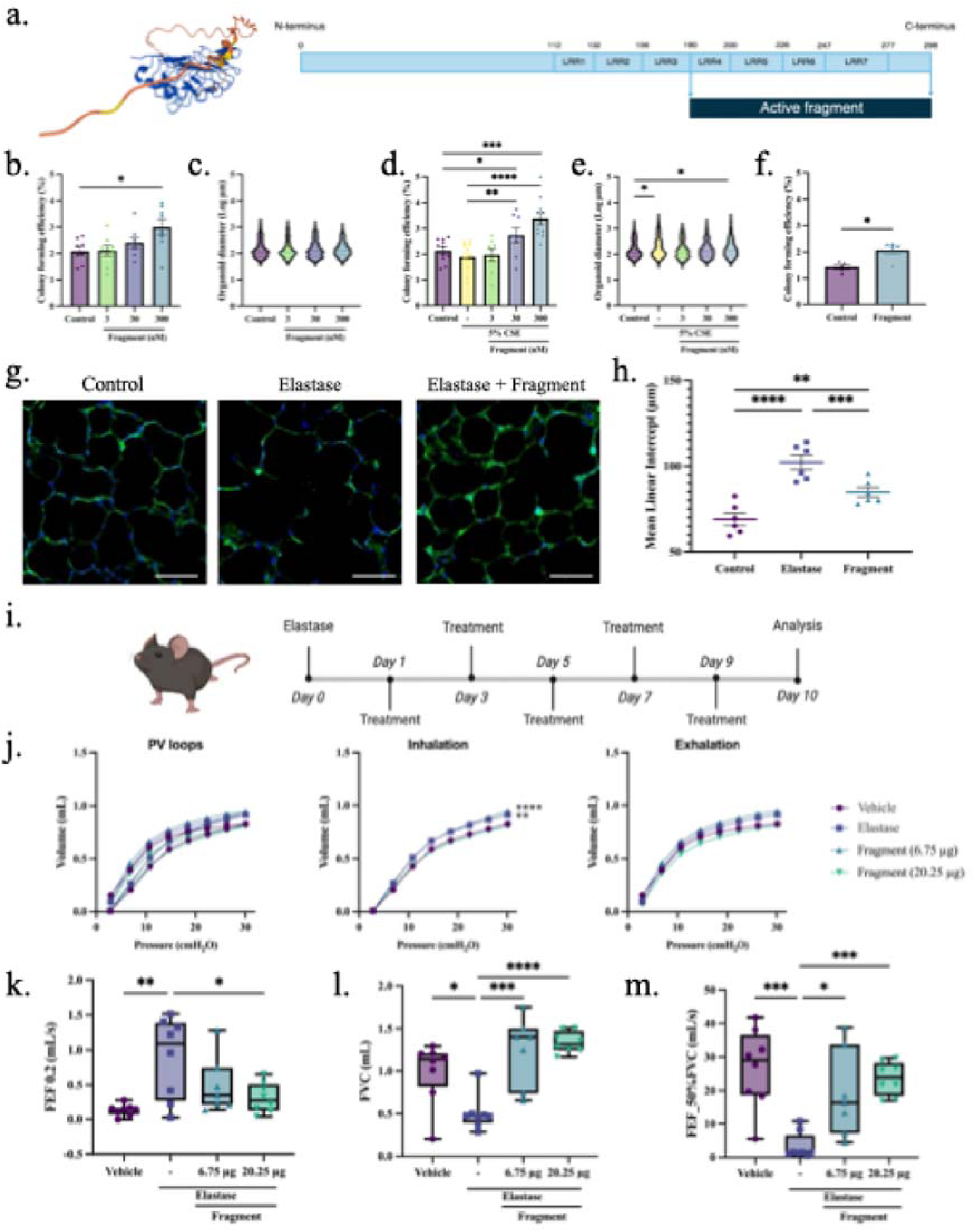
An active fragment of osteoglycin induces alveolar epithelial growth and improves lung function. **a** Predicted alpha-fold structure of osteoglycin, showing the full-length protein comprising seven LRRs and an active fragment (LRR4 - LRR7). **b** CFE of murine organoids treated with OGN fragment (mean ± SEM, N=8, paired one-way ANOVA, Dunnett test). **c** Log of murine organoid diameter (median, N=8, Kolmogorov-Smirnov test, Bonferroni correction: α = 0.0167). **d** CFE of CSE-exposed murine organoids (mean ± SEM, N=8-11, paired one-way ANOVA, Dunnett test). **e** Log of CSE-exposed murine organoid diameter (median, N=8-11, Kolmogorov-Smirnov test, Bonferroni correction: α = 0.0125). **f** CFE of human organoids (mean ± SEM, N=5, paired Student T-test). **g** PCLS stained for F-actin filaments (green) and Dapi (blue) (scale = 50 µm) after treatment with vehicle control, elastase, or elastase + fragment (300 nM). **h** Mean linear intercept measurements following treatments (µm, mean ± SEM, N=6, paired one-way ANOVA, Dunnett test). **i** Schematic of in vivo murine elastase-induced lung injury model. **j** Pressure-volume loops for lung distensibility (N=7-8, two-way ANOVA, Dunnett test). **k-m** Lung function parameters: FEF 0.2, FVC, and FEF_50%FVC measured with FlexiVent (median ± min/max data point, N=7-8, one-way ANOVA, Dunnett test). Controls are identical to Fig. 4. Statistically significant comparisons: *p < 0.05, **p < 0.01, ***p < 0.001, and ****p < 0.0001.

We then investigated the effect of OGN fragment on differentiated epithelial cells across various lung injury models. Murine PCLS exposed to elastase and treated with active fragment showed significantly decreased LMI, indicating reduced lung injury (Fig. 6g-h). Next, we evaluated whether the active fragment of OGN could improve lung function in vivo in a lung injury murine model. Upon inducing lung injury with elastase, mice received low (6.75 µg) or high (20.25 µg) doses of OGN fragment (Fig. 6i). Elastase treatment induced a decrease in elastic recoil, as shown by a decreased tissue elastance and increased compliance (Fig. S12a-c). Although the active fragment of OGN did not elicit significant changes in elastance and compliance, as compared to elastase, we observed a dose-dependent trend indicating improvement in these lung tissue characteristics (Fig. S12a and c). Elastase did not induce any other significant changes in tissue characteristics, such as resistance, tissue damping, and Newtonian resistance (Fig. S12d-f). Additionally, pressure-volume loops of emphysematous mice displayed a typical significant upward shift (Fig. 6j-k), suggesting that elastase diminished the distensibility of the lungs. Diminished distensibility was absent in emphysematous mice treated with OGN fragment. In addition to lung tissue characteristics, we assessed lung function parameters in different experimental groups. The forced expiratory flow at 0.2 seconds (FEF0.2) significantly increased upon elastase exposure, while the forced vital capacity (FVC) and forced expiratory flow at 50% of forced vital capacity (FEF_50%FVC) were significantly decreased (Fig. 6k-m). Treatment with OGN fragment inhibited the harmful effects of elastase on lung function significantly (Fig. 6k-m). We did not observe any significant changes from the control (vehicle) measurements in forced expiratory volume at 0.2 seconds, forced expiratory volume at peak expiratory flow, peak expiratory flow, and time to reach peak expiratory pressure (Fig. S12g-j). Taken together, the active fragment of OGN ameliorates elastase-induced lung injury in PCLS and improves lung function in mice, thereby underscoring its potential as a promising therapeutic approach.

## DISCUSSION

As there are currently no clinically approved pharmacological treatments to prevent the progression of or reverse tissue destruction in the distal lung, we aimed to discover a new potential drug for (re-)activating alveolar epithelial progenitor cells, leveraging the lung fibroblast secretome. Utilizing a proteomics-guided drug discovery strategy, we identified twelve potential factors, with OGN showing the most pronounced effects on alveolar organoid formation. RNA sequencing demonstrated that OGN is related to pathways involving ROS defense mechanisms and enhanced epithelial-fibroblast crosstalk. OGN expression was reduced in the lung tissue of smoke-exposed mice and current smokers, with a similar tendency in COPD patients. Moreover, an active fragment of OGN, containing LRRs 4-7, showed comparable effects on epithelial progenitor activation as the entire OGN protein and improved lung function in a mouse model of elastase-induced lung injury. These findings identify the active fragment of OGN as a promising drug candidate to prevent the progression of or reverse epithelial tissue destruction in the distal lung.

In the lungs, alveolar epithelial repair is predominantly regulated by AT2 cells, which are instructed by meticulously orchestrated molecular cues from neighboring cells within the alveolar niche.^2,15^ The supportive effect of alveolar fibroblasts on epithelial progenitor cells was evident in our organoid assay, where lower organoid numbers were observed when co-cultured with reduced fibroblast numbers. Furthermore, our findings indicate that lung fibroblasts support alveolar organoid formation in a paracrine fashion through the secretion of EVs and SFs. Consistent with these findings, several studies have shown EVs enhance epithelial organoid formation across tissues^32–34^, and that fibroblast-derived SFs are essential for organoid formation.^15,35,36^

Among the candidate factors identified, OGN emerged as the most promising. In the organoid model, an active fragment of OGN (LRRs 4-7) induced alveolar organoid formation, suggesting that this smaller fragment is sufficient for activating alveolar epithelial progenitor cells. This fragment also effectively alleviated elastase-induced lung injury in PCLS and a mouse model. To our knowledge, OGN or its functional fragment has not been studied previously in the context of COPD treatment. While Lee and co-workers revealed that intraperitoneal therapy of mice with OGN improved their whole-body glucose metabolism and insulin homeostasis^37^, most studies on OGN focus on cancer research. To this extent, models of OGN overexpression are usually exploited, which show contradicting effects on epithelial cell proliferation.^38–40^ In our organoid model, no changes in organoid size were observed with OGN or its active fragment, implying no discernible changes in proliferation, which could be context-dependent.

By generating transcriptomic signatures of epithelial progenitor cells, we uncovered the upregulation of several protective pathways, including the detoxification of ROS and iron uptake and transport. We have previously shown that anti-oxidants, such as n-acetyl cysteine and mitoquinone, could reverse the negative impact of diesel exhaust particles on lung epithelial progenitors.^41^ Additionally, other anti-oxidants effectively reduced lung injury, inflammation, and oxidative stress in mice.^42^ Interestingly, the relationship between OGN and the detoxification of ROS has not yet been studied.^43^ Besides building on the identified protective pathways, the focus on iron uptake and transport might be crucial in explaining the protective effects of OGN on epithelial progenitors. Genome-wide association studies have highlighted a pathogenetic role for abnormal iron homeostasis in COPD.^44,45^ Moreover, intracellular accumulation of iron and lipid peroxidation are associated with ferroptosis, a newly identified type of cell death.^46^ Recent studies found that increased ferroptosis and disrupted iron homeostasis were linked to COPD.^46,47^

Lung epithelial progenitor cell behavior is known to be tightly coordinated by alveolar fibroblasts through paracrine signaling of several growth factors, including FGFs and HGF.^48,49^ We found that *Ffg7*, *Ffg10*, and *Hgf* expression was significantly increased in response to OGN treatment on fibroblasts. These growth factors stimulate epithelial cell proliferation and survival, which could contribute to the OGN-induced alterations in organoid formation.^50,51^ Indeed, FGF7, FGF10, and HGF were previously shown to increase lung organoid formation^17,51^ and FGF7 and FGF10 support maturation of lung organoids toward distal lung lineages.^52,53^ In addition, FGF signaling was shown to have antifibrotic effects in experimental pulmonary fibrosis through epithelial progenitor proliferation and inhibition of TGF-β.^54^ As the receptor with which OGN interacts in the lung is unknown, it remains unclear whether OGN directly influences alveolar epithelial cell behavior or indirectly through the intermediary action of fibroblasts. However, the total number of differentially expressed genes in fibroblasts following OGN treatment is significantly higher compared to those in the epithelial cell population, suggesting a predominant effect on fibroblast function. Additionally, while OGN treatment upregulated key pro-regenerative factors such as FGF7, FGF10, and HGF in fibroblasts, the effects of EVs and SFs on these factors appeared somewhat distinct. This difference suggests that the observed regenerative effects may not be solely linked to OGN, indicating that EVs and SFs could exert independent effects mediated through other messenger molecules.

In this study, we describe for the first time the distribution of OGN in human lung tissue on a transcriptomic and protein level. We observed proportionally lower OGN protein expression in the lung tissue of current smokers compared to non-smokers, with ex-smokers showing intermediate levels. This suggests a lasting effect of cigarette smoke exposure on OGN expression. These findings align with Lin and colleagues, who reported reduced OGN mRNA expression in lung tissue of severe COPD patients.^55^ Interestingly, Huang and colleagues report upregulation of OGN in lungs and myofibroblasts obtained from lung fibrosis murine models.^56^ Koloko Ngassie et al. revealed a positive correlation between increasing age and enhanced OGN protein expression in human lung tissue derived from patients with no history of chronic lung disease.^25^ Considering our data, this suggests that OGN expression in the healthy human lung increases with age, whereas this process appears compromised in individuals who smoke. Such impairment could contribute to the hampered alveolar repair and regeneration mechanisms observed in COPD patients. Since this effect is also observed in healthy smokers, this suggests that impaired OGN expression could be an early event towards COPD progression or that a second hit is required for COPD progression. Based on our various disease models, we speculate that therapeutically supplementing COPD patients with OGN or its active fragment could potentially (re-)activate lung epithelial repair or slow down the progression of tissue damage. While alveolar epithelial repair is a crucial step toward regaining functioning alveolar tissue, it is essential to consider that the complete restoration of alveolar function involves addressing the entire alveolar niche, which includes various cell types and extracellular matrix components. To further this research, it is crucial to determine which COPD patients would benefit most from this therapy, at what stage of the disease, and to consider the differences between current and former smokers. Notably, it is essential to acknowledge that OGN is expressed in various tissues, including bone, muscle, and adipose tissue.^30^ Glycosylation and splice variants of OGN can make it cell-type and tissue-specific, which would require further studies.

When contemplating the therapeutic application of OGN as a regenerative agent for (re-)activating alveolar epithelial repair, careful consideration of potential adverse effects associated with exogenous administration of this protein is essential. As a proteoglycan, OGN has been linked to the induction of fibrosis.^57^ For instance, Shi et al. demonstrated that microRNA-140 overexpression inhibits pulmonary fibrosis by downregulating OGN through the Wnt signaling pathway.^58^ The role of OGN in fibrosis regulation extends beyond the lungs, encompassing organs like the heart and kidney.^30^ However, our RNAseq analysis offers a contrasting perspective, indicating a tendency for downregulation of the predicted gene signature for myofibroblasts and fibrosis-related genetic markers upon OGN treatment. It is hypothesized that OGN regulates fibrosis through the crosslinking of collagen via the N-terminus.^57,59^ Following this rationale, therapeutic use of OGN might induce fibrosis by increasing cross-linking. However, in our assays, we show that a fragment encompassing LLRs four through seven exhibits similar effects on epithelial progenitor cell behavior. This may present an attractive approach to circumvent fibrosis initiation in individuals by eliminating the structural component crosslinking activities of the entire OGN protein.

A limitation of this study is the relatively small number of patients included in the different analyses. The variability observed in the data across these studies could be attributed to the heterogeneity of COPD, variations in patient inclusion criteria, and differences in analysis methods. In the context of IHC staining on OGN expression in the lungs, we must acknowledge that our examination is confined to the remaining lung tissue. Particularly in end-stage COPD patients, substantial tissue damage exists, posing challenges to the accurate quantification of OGN expression across different disease stages.

In conclusion, lung fibroblasts support the formation of organoids through the secretion of EVs and SFs. Using a proteomics-guided drug discovery strategy, we identified OGN to activate alveolar epithelial progenitor cells. OGN induces a wide range of transcriptional changes, including the upregulation of protective pathways in epithelial progenitors and growth factor-mediated epithelial-fibroblast crosstalk. Expression of OGN was reduced in response to cigarette smoke and the lung tissue of smokers. Furthermore, an active fragment of OGN was shown to have similar properties on epithelial progenitor activation and improved lung function parameters in mice with lung injury. While this study focused on COPD, potential benefits of OGN could extend to other conditions marked by deficient epithelial progenitor function across organs.

## Acknowledgments

Part of the work has been performed at the UMCG Imaging and Microscopy Center (UMIC). Specifically, we thank Klaas Sjollema for the help provided with confocal imaging. We are grateful for the support provided by the animal facility in Groningen in ensuring the well-being and care of the animals. The figures contained in this article were partially created with BioRender.com.

## Data and materials availability

All data associated with this study are in the paper or the Supplementary Materials. The complete RNA sequencing dataset is available in GEO at GSE271272. The proteomics dataset of the secretome of human lung fibroblasts is available in the PRIDE proteomics identifications database (PXD053766). For material requests, researchers should contact AN or RG.

## Supplementary Information

Supplementary information is available at Experimental and Molecular Medicine’s website.

