## Supplemental Materials for "Fibroblast-derived Osteoglycin Promotes Epithelial Cell Repair"

Luke van der Koog *et al.*

**The PDF includes:**

Material and methods

Figures. S1 to S12

Tables S1 to S7

**Supplementary Materials and Methods**

**Antibodies and reagents**

An anti-pro-SPC antibody (AB3786) was purchased at Sigma Aldrich, and acetylated ⍺ tubulin (sc-23950) was obtained from Santa Cruz. Secondary antibodies for Alexa Fluor™ 488 (A21206) and 568 (A10037) were purchased at Thermo Fisher. Recombinant human OGN full protein was purchased at Biorbyt (orb383003), and the human OGN fragment (amino acid 180-298) was obtained from LifeSpan Biosciences (LS-G15022). Pancreatic porcine elastase for *in vitro* (E0258-5MG) and *in vivo* (324682-250U) experiments were obtained from Sigma Aldrich.

**Cell culture**

Mouse fibroblasts, CCL206 cells (ATCC, Mlg2908), were cultured in DMEM/F12 medium (Gibco) supplemented with 10% (v/v) fetal bovine serum (FBS) (Sigma Aldrich, 12103C), 100 U/ml penicillin/streptomycin (Gibco, 15070-063), 2 mM L-glutamine (Gibco, 25030-024), and 1% amphotericin B (Gibco, 15290026) within a humidified atmosphere under 5% CO_2_/95% air at 37 °C. The human fetal mesenchymal lung fibroblast MRC5 cell line (Sigma Aldrich, 05081101) was cultured in Ham’s F12 medium (Thermo Fisher, 11320033) supplemented with 10% (v/v) FBS, 100 U/mL penicillin/streptomycin, and 2 mM L-glutamine within a humidified atmosphere under 5% CO_2_/95% air at 37 °C.

For organoid experiments, before both fibroblast types (CCL206 and MRC5) were co-cultured with primary epithelial cells, the proliferation of fibroblasts was inactivated by incubation with mitomycin C (10 µg/mL) in growth medium (Sigma Aldrich, M4287-5X2MG) for 2 h. After incubation with mitomycin C, the fibroblasts were washed with warm PBS and then recovered for 1 h in mitomycin C-free growth medium.

**Purification of EVs and SFs using ultrafiltration and size exclusion chromatography**

MRC5 lung fibroblasts were initially expanded from a single T75 flask to 13 T175 flasks through several passaging steps. Once the MRC5 cells reached approximately 80% confluency, the cells were serum starved for a period of 72 hours. The conditioned medium was collected and centrifuged at 4000 g for 15 mins, before filtering using a 0.45 µm bottle top filter. The conditioned medium was then concentrated using 100 kDa Amicon Ultra-15 Centrifugal Filter Units (Merck, UFC900324) by repeated centrifugation at 4000 g at 4 °C to a final volume of ~500 µL. A 30-cm-long, 1-cm-diameter column packed with Sepharose CL-2B (Cytiva, 17-0140-01) was washed with PBS prior to loading the concentrated medium. Thereafter, 24 x 1 mL fractions were collected from the column. Size exclusion column (SEC) fractions 8 till 12 were considered to be the EV-enriched fractions, while fractions 18 till 22 served as the SF-enriched fractions. Both the EV- and SF-fractions were pooled and concentrated using 100 kDa Amicon Ultra-4 Centrifugal Filter Units (Merck, UFC801024) to a final volume of ~600 µL. Subsequently, the EV- and SF-enriched samples were sterilized using Costar Spin-X centrifuge tube filters (Merck, CLS8160-96EA) with a 0.22 µm pore size. Samples were stored at -80 °C until further use. For the treatment of organoid cultures, we used concentrations of EVs and SFs that showed the strongest impact on colony forming efficiency in murine organoids (Fig. S2). Multiple purification batches were applied for each organoid experiment to ensure consistency and reproducibility.

**Primary alveolar epithelial isolation**

For murine organoid cultures, we have used a total of 54 mice (male and female (1:1), eight to fourteen weeks old). In brief, the lungs of mice were flushed through the heart with PBS, instilled with dispase (Corning, 354235), and incubated at room temperature for 45 mins. To obtain a single-cell suspension, lung tissue was digested with DNase 1 (VWR, A3778.0500). Using the QuadroMACS™ Separator (Miltenyi Biotec, 130-091-051) and a mix of antibody-bound magnetic microbeads, the cell suspension was negatively selected for CD31 (Miltenyi Biotec, 130-097-418) and CD45 (Miltenyi Biotec, 130-052-301). Subsequently, to obtain Epcam^+^ cells, CD31^-^/CD45^-^ cells were positively selected with anti-mouse CD326 microbeads (Miltenyi Biotec, 130-105-958).

Human lung tissue was stored in MACS® Tissue Storage Buffer (Miltenyi Biotech, 130-100-008) until further processing. Lung tissue was cut into small pieces (~ 1 mm^3^) and transferred to a dissociation mixture containing 1% penicillin/streptomycin, 1 mg/mL collagenase/dispase (Roche, 11097113001), and 1.8 µg/mL DNase 1 in PBS. The tissue was further dissociated using a gentleMACS™ Octo Dissociator with heaters (130-096-427, Miltenyi Biotec) for 20 mins at 37 °C. The obtained single-cell suspension was washed, and red blood cells were lysed using lysis buffer (ammonium chloride (155 mM), potassium bicarbonate (1 mM), titriplex III (0.001 mM), and 10 µg/mL DNase 1 in ultra-pure water) for 10 mins at 4 °C. Selection for EpCAM^+^ cells was similar to that described above for murine lung tissue.

**Epithelial organoid culture**

For murine organoids, freshly isolated Epcam^+^ cells were combined with CCL206 murine lung fibroblasts at a 1:1 ratio (10,000 cells each) in DMEM/F12 containing 10% (v/v) FBS. The cell suspension was then diluted 1:1.5 (v/v) with Corning® Matrigel® Membrane Matrix (Corning, 356234) and was then seeded into transwell inserts (Greiner, 662641) in 24-well plates (100 µL/insert). Similarly, human organoids were generated by co-culturing freshly isolated EpCAM^+^ cells with proliferation-inactivated MRC5 lung fibroblasts. The Matrigel™ was allowed to solidify at 37 °C for 30 mins. Upon solidification, 410 µL of organoid medium (DMEM/Ham’s F12 supplemented with 5% FBS, 1% penicillin/streptomycin, 1% L-glutamine, 1% amphotericin B, 0.025 ‰ epidermal growth factor (EGF) (Sigma Aldrich, SRP3196-500UG), 1% insulin-transferrin-selenium (Gibco, 51300044), and 1.75 ‰ bovine pituitary extract (Thermo Fisher, 11568866)) was added underneath the insert. On the day of seeding, 10 µM Y-27632 dihydrochloride (Axon, 1683) was added to inhibit Rho-Kinase selectively. Organoid cultures were cultured at 37 °C with 5% CO_2_. The medium was refreshed every 2-3 days. The total number and diameters of organoid structures (>50 µm) was measured on day 14 using NIS-Elements with bright field microscopy (20x magnification).

**Cigarette smoke extract**

To generate 100% cigarette smoke extract (CSE), the smoke from two 3R4F research cigarettes (Tobacco Research Institute) without a filter was introduced into 25 mL of warm fibroblast culture medium.^3^ The smoke was delivered into the medium using a peristaltic pump (Watson Marlow 323 E/D) at a speed of 45 rpm. CSE was freshly prepared before each set of experiments. For organoid experiments, we used 5% CSE in organoid growth medium.

**Immunofluorescence staining organoids**

Organoids were fixed in a 1:1 (v/v) mixture of acetone and methanol for 12 mins at -20 °C.^3^ After fixation, 1 mL of PBS with 0.02% sodium azide (Merck, 6688) was added to the well underneath the insert. Organoids were kept at 4 °C for 1 week after fixation. Organoids were blocked using a blocking solution containing 5% bovine serum albumin (BSA), 2% normal donkey serum (Jackson Immuno Research, 017-000-121), and 0.1% Triton-X100 in PBS overnight at 4 °C. The next day, surfactant protein C (SPC) and acetylated ⍺ tubulin (ACT) primary antibodies diluted 1:200 in 2% BSA, 2% normal donkey serum, and 0.1% Triton-X100 in PBS were added for 48 h. Organoids were then washed three times with PBS for 30 mins, and secondary antibodies Alexa Fluor™ 488 and 568 diluted 1:200 in 2% BSA, 2% normal donkey serum, and 0.1% Triton-X100 in PBS were added for 2 h at room temperature. After washing three times with PBS for 30 mins, the organoids on the insert membrane were transferred to a glass slide with a mounting medium containing DAPI, and a coverslip was applied afterward. The slides were kept at 4 °C, and confocal images were acquired using a Nikon Eclipse Ti2 microscope.

**Bulk RNA sequencing analysis**

Total RNA was extracted from cells resorted from organoids using the NucleoSpin RNA isolation kit (Bioké, 740955.50) according to the manufacturer’s instructions. Bulk RNA sequencing (RNAseq) on resorted Epcam+ and fibroblasts from organoids was performed by GenomeScan (www.genomescan.nl) using an Illumina NovaSeq 6000 sequencer. The analysis procedure comprised several steps, including data quality control, adapter trimming, alignment of short reads, and feature counting. To ensure the integrity of the library preparation, calculations were performed to assess the ribosomal (and globin) content. Additionally, checks were conducted to identify potential samples and barcode contaminations. Quality control tools, such as FastQC v0.34 and FastQA, were employed to establish a set of standard quality metrics for the raw dataset. Prior to alignment, Trimmomatic v0.30 was utilized to remove adapter sequences from the reads. The reads of each sample were aligned against the ensemble mouse reference GRCm38 (patch 6). Principal component analyses were performed in R using the R package DESeq2 in order to visualize the overall effects of experimental covariates as well as batch effects. The same R packages were used to identify differentially expressed genes (DEGs) between control and treated samples following standard normalization procedures. Gene set enrichment analysis (GSEA) of the top 50 differentially regulated genes was performed with ShinyGO 0.77 (www.bioinformatics.sdstate.edu/go/). Kyoto Encyclopedia of Genes and Genomes (KEGG) was used as reference database, and the statistically significant pathway enrichment with FDR q value < 0.05 are reported.

**Immunohistochemical staining on human lung slides**

Lung tissue was embedded in paraffin and cut into 6 µm thick sections. These sections were deparaffinized and rehydrated, after which antigen retrieval was performed with 10 mM citrate buffer (pH 6). Endogenous peroxidase activity was blocked by 0.3% hydrogen peroxidase (H_2_O_2_), followed by overnight incubation at 4 °C with a primary OGN antibody (1:400) (Abcam, ab168348) in 1% BSA-PBS. Subsequently, sections were washed and incubated with horseradish peroxidase (HRP)-conjugated secondary antibody diluted 1:100 in 1% BSA-PBS containing 2% normal human serum. Staining was then visualized upon 5 mins incubation with Vector NovaRED Susbtrate (VectorLaboratories, SK-4800). Sections were then counterstained with hematoxylin, mounted, and scanned using a Hamamatsu NanoZoomer 2.0HT digital slide scanner at 40x magnification. The intensity and area of positive OGN staining in whole lung tissue and parenchyma were analyzed as described previously.^25^

**OGN expression analysis in human lung slides**

Analysis of the intensity and area of positive OGN staining in whole lung tissue and parenchyma were analyzed as described previously.^25^ Briefly, Aperio ImageScope software V.12.4.3 (Leica Biosystems) was used to extract images from the scans. For whole lung tissue analysis, scanned images were used after removal of artefacts. This step was followed by extracting specific areas, including airway wall, bronchial epithelium, and blood vessels, using Adobe Photoshop software (Adobe Inc. CA), to analyze OGN expression in the parenchyma. Fiji/ImageJ software was used to quantify the intensity and area of positive staining of OGN in whole tissue and parenchyma. The formula used to calculate the percentage of area stained positive for the protein is as follows:

$$Area \left( \% \right)= \frac{Area (NovaRed)}{Area (Total)} x 100\%$$

For the quantification of the intensity of the staining, pixel intensities, ranging from 0 to 255 of separated NovaRed images were analyzed with Fiji/ImageJ. A value of 0 corresponds to the darkest shade, while 255 represents the lightest shade. This reciprocal intensity is directly proportional to the quantity of positive NovaRed pixels detected in the image analysis and is calculated with the following equation:

$$Mean intensity=255- \frac{Sum of intensities of pixels positive for NovaRed}{Total number of pixels positive for NovaRed}$$

**Murine precision-cut lung slices**

Murine lungs were inflated with 1.5 mL 1.5% (w/v) low melting agarose solution (Gerbu Biotechnik). After inflation, the agarose was allowed to solidify at 4 °C for 15 mins before the lungs were harvested. A tissue slicer (Leica VT 1000 S Vibratome line) was used to cut lung slices with a thickness of 250 µm. The lung slices were extensively washed and subsequently cultured in DMEM (Gibco, 42430-025) supplemented with sodium pyruvate (1 mM), MEM non-essential amino acids mixture (1:100, Gibco, 11140-050), gentamycin (45 µg/mL, Merck, G1397), penicillin/streptomycin (100 U/mL), and amphotericin B (1.5 µg/mL, Gibco, 15290-026) in 12-well culture plates, using three slices per well. Slices from the same mouse were matched based on lung region (middle region of the left lung lobe or middle region of superior right lung lobe). To induce emphysematous changes, matched slices were treated with 2.5 µg/mL elastase for 16h. Slices were treated with 10 µg/mL OGN or 4.48 µg/mL OGN active fragment for 40h, overlapping with the 16h elastase treatment.

**Immunofluorescence staining PCLS**

PCLS were fixed for 15 mins at 4 °C with 4% paraformaldehyde (Sigma Aldrich, P6148) and then washed with PBS. To visualize the parenchyma of PCLS, actin filaments were stained with Alexa Fluor™ 488 Phalloidin (Thermo Fisher, A12379) for 15 mins at room temperature. After incubation, the slices were washed with PBS and transferred onto a glass slide with two drops of mounting medium containing DAPI (Abcam, 104139). Fluorescence imaging was performed using a confocal laser scanning microscope equipped with a true confocal scanner (SP8 Leica) using a 20x lens. All images were acquired within the linear range, with an image resolution of 1024 x 1024 pixels and a pinhole size of 1 Airy unit to avoid local saturation. The presented images represent a single z-scan. To assess the degree of lung injury in PCLS, we measured the mean linear intercept (LMI), which represents the average distance of free airspace in 5 fields per animal, as previously described.^20^

**Quantification of the degree of lung injury in PCLS**

To assess the degree of lung injury in PCLS, we measured the mean linear intercept (LMI), which represents the average distance of free airspace, in 5 fields per animal, as previously described.^20^ However, due to the ex vivo treatment with elastase, we were unable to follow the standard procedure of fixing the murine lungs at 25 cmH2O for 24h, which is typically used for LMI determination. In this PCLS model, the use of varying pressures to fill the murine lungs with agarose results in variations in LMI measurements between animals. Nevertheless, each animal served as its own control, eliminating the variability caused by different pressure conditions applied to the lungs. However, caution should be exercised when extrapolating these numbers to the in vivo setting.

**Elastase in vivo study**

For this study, 48 C57BL/6J male and female mice (ratio 1:1) were included and randomly allocated into six experimental groups. Pulmonary emphysematous changes were induced by intratracheal instillation of pancreatic porcine elastase (40 U/kg body weight) in 40 µL sterile PBS on day zero. Animals were treated every other day from day zero till day nine (5 treatments in total) with OGN fragment (6.75 µg or 20.25 µg). After ten days, mice were sacrificed by exsanguination under anesthesia, after which the therapeutic effects were examined. To ensure objective analysis, samples for *in vivo* experiments were blinded before analysis.

**Lung function measurements**

Respiratory function was measured using a FlexiVent system module 2 (Scireq). Mice were anesthetized with Dexdomitor® and Ketamine® and a muscle relaxant, rocuronium bromide (Fresenius Kabi, 10 mg/mL) was administered. Mice were ventilated with a tidal volume of 10 mL/kg at a frequency of 150 breaths/min in order to reach a mean lung volume similar to that of spontaneous breathing. Lung function parameters were assessed using pre-installed protocols for SnapShot, Primewave perturbation, and forced expired volume maneuver using the Flexiware V8.3.0 software. Three recordings per animal were taken.

**Supplementary Figures:**

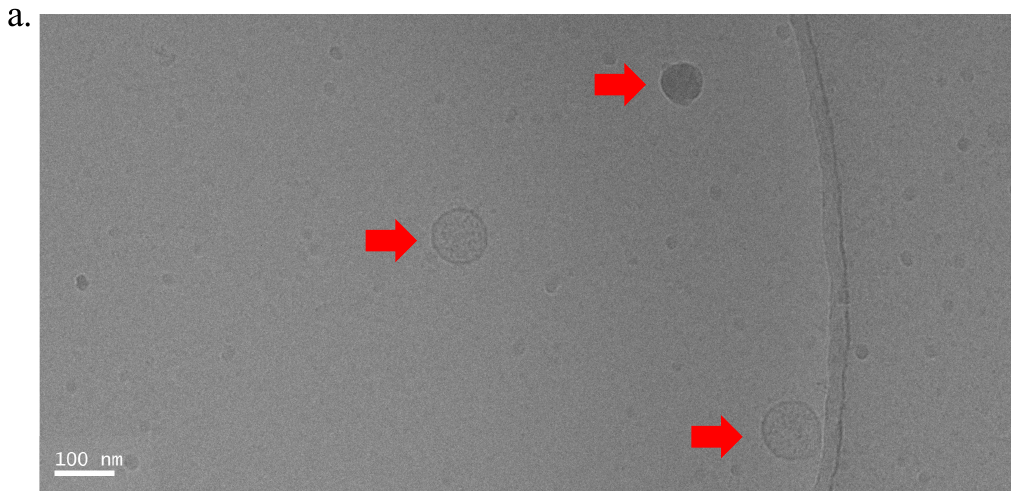

**Fig. S1. Lung fibroblast-derived EVs visualized by Cryo-TEM.**

Cryo-TEM image of pooled lung fibroblast-derived EV-enriched fractions. Red arrows indicate EV-structures. Scale = 100 nm.

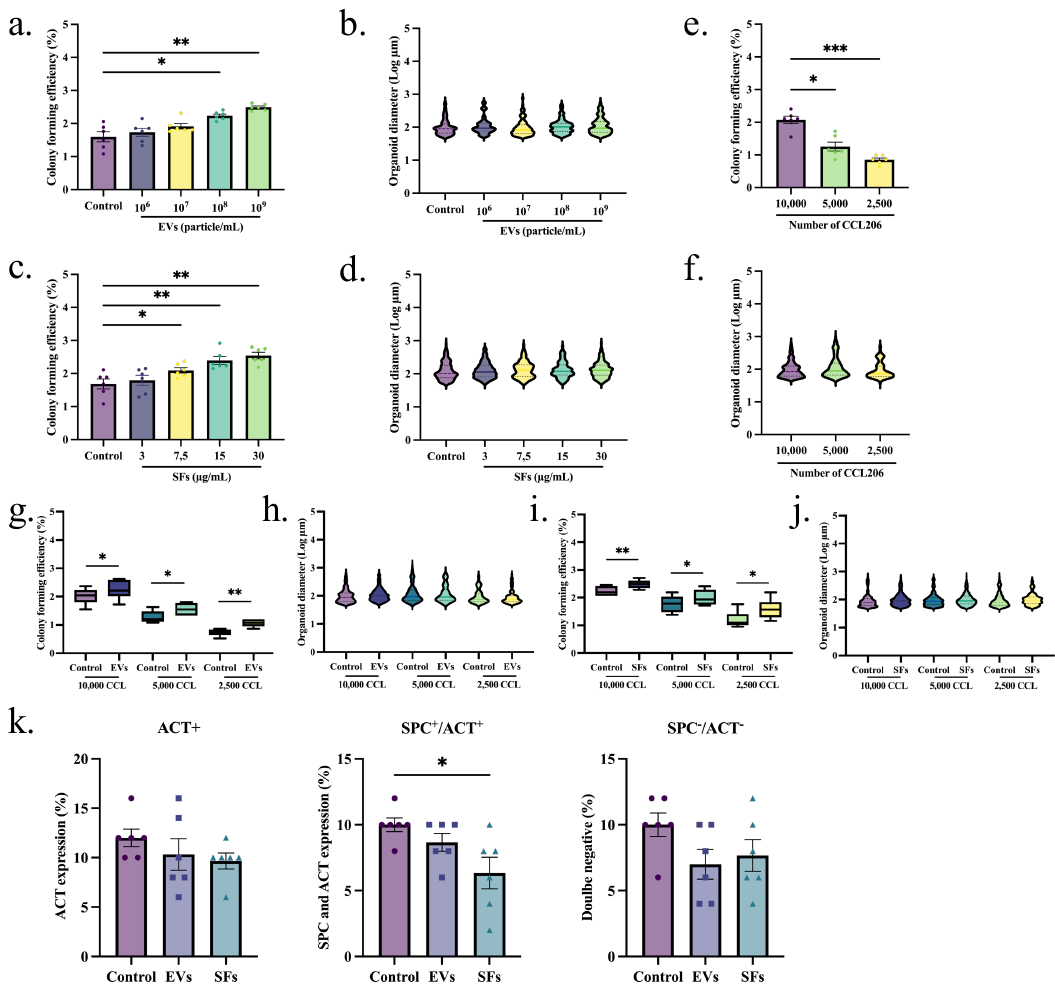

**Fig. S2. Lung fibroblast-derived EVs and SFs support murine alveolar organoid formation.**

**a** Colony forming efficiency of murine organoids with increasing concentrations of EVs (mean ± SEM, N=6, paired Friedman test). **b** Log of murine organoid diameter with increasing concentrations of EVs (median is shown, N=6, Kolmogorov-Smirnov test (after Bonferroni correction: α = 0.0125)). **c** Colony forming efficiency of murine organoids with increasing concentrations of SFs (mean ± SEM, N=6, paired Friedman test). **d** Log of murine organoid diameter with increasing concentrations of SFs (median is shown, N=6, median is shown, N=6, Kolmogorov-Smirnov test (after Bonferroni correction: α = 0.0125). **e** Colony forming efficiency of murine organoids with decreasing lung fibroblasts (CCL206) (mean ± SEM, N=6, paired Friedman test). **f** Log of murine organoid diameter with decreasing numbers of lung fibroblasts (CCL206) (median is shown, N=6, Kolmogorov-Smirnov test (after Bonferroni correction: α = 0.0025)). **g** Colony forming efficiency of murine organoids with decreasing numbers of lung fibroblasts (CCL) treated with EVs (10^9^ EVs/mL) (mean ± SEM, N=6, paired Friedman test). **h** Log of murine organoid diameter with decreasing numbers of lung fibroblasts (CCL) treated with EVs (mean ± SEM, N=6, Friedman). **i** Colony forming efficiency of murine organoids with decreasing numbers of lung fibroblasts (CCL) treated with SFs (30 µg/mL) (mean ± SEM, N=6, paired Friedman test). **j** Log of murine organoid diameter with decreasing numbers of lung fibroblasts (CCL) treated with SFs (30 µg/mL) (mean ± SEM, N=6, Friedman). **k** Immunohistochemistry quantification for ACT^+^, SPC^+^/ACT^+^, and SPC^-^/ACT^-^ organoids (mean ± SEM, N=6, paired one-way ANOVA with Tukey test for multiple testing). Statistically significant comparisons are represented by *p < 0.05, **p < 0.01, and ***p < 0.001.

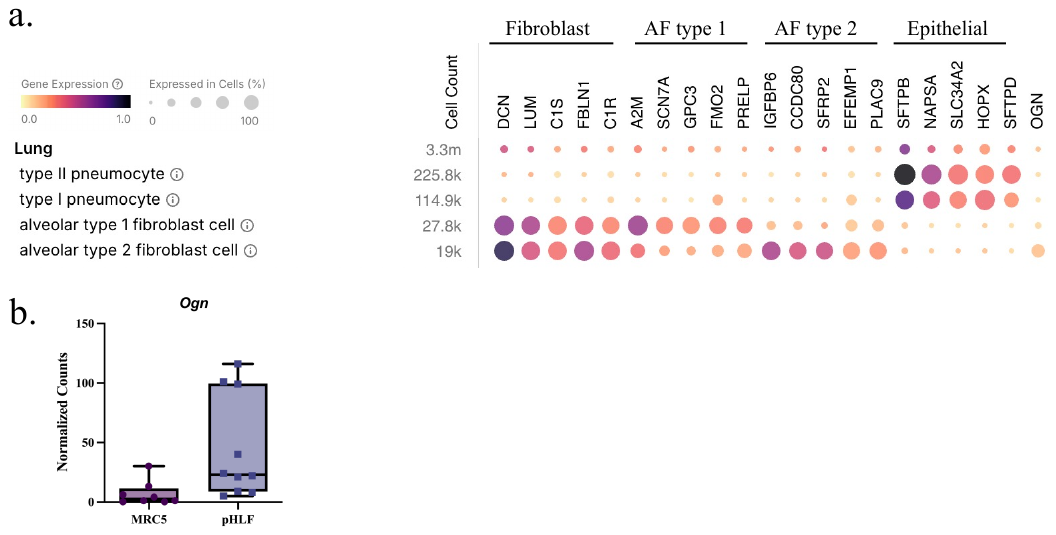

**Fig. S3. OGN expression in lung fibroblasts.**

**a** The expression of markers for fibroblasts (*DCN*, *LUM*, *C1S*, *FBLN1,* and *C1R*), alveolar fibroblast (AF) type 1 (*SCN7A*, *GPC3*, *FMO2*, *PRELP*), AF type 2 (*IGFBP6*, *CCDC80*, *SFRP2*, *EFEMP1*, *PLAC9*), epithelial cells (*SFTPB*, *NAPSA*, *SLC34A2*, *HOPX*, and *SFTPD*), and *OGN*. Data were extracted from a public scRNA-Seq dataset (https://lungmap.nl/). **b** Normalized counts of *OGN* in MRC5 fibroblasts and pulmonary human lung fibroblasts (pHLF).

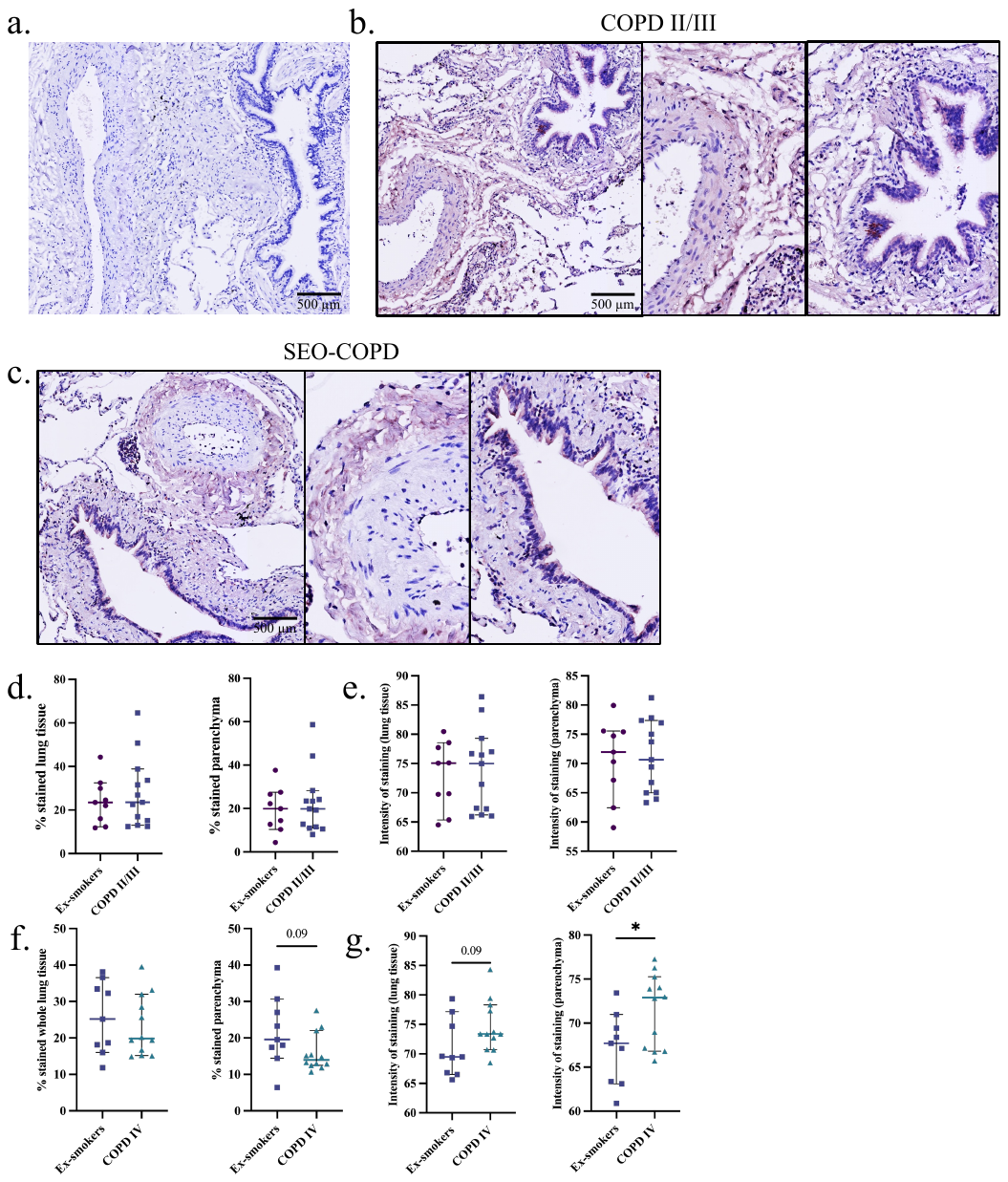

**Fig. S4. OGN expression in moderate-severe COPD lung tissue.**

**a** Example image of negative control staining in human whole lung tissue. **b-c** Example image of OGN staining in whole lung tissue in moderate-severe COPD (II/III) and SEO-COPD (scale = 500 µm). **d** Positively stained area percentage (%) for OGN in whole lung tissue and parenchyma in COPD II/III patients. **e** Intensity of staining for OGN in whole lung tissue and parenchyma in COPD II/III patients. **f** Positively stained area percentage (%) for OGN in whole lung tissue and parenchyma in SEO-COPD patients. **g** OGN staining intensity in whole lung tissue and parenchyma in SEO-COPD patients. Statistically significant comparisons are represented by *p < 0.05.

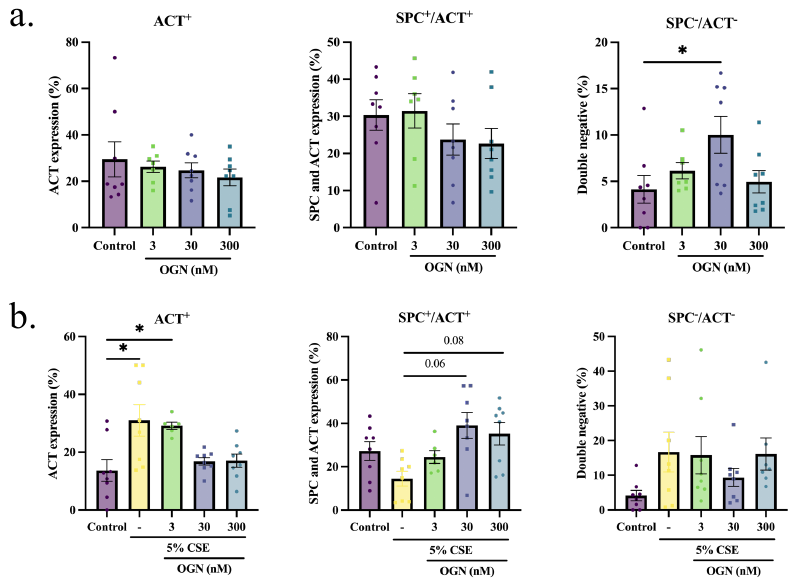

**Fig S5. Osteoglycin supports organoid formation and differentiation.**

**a** Immunohistochemistry quantification for ACT^+^, SPC^+^/ACT^+^, and SPC^-^/ACT^-^ organoids (mean ± SEM, N=8, paired one-way ANOVA with Tukey test for multiple testing). **b** Immunohistochemistry quantification for ACT^+^, SPC^+^/ACT^+^, and SPC^-^/ACT^-^ organoids in the presence of CSE (mean ± SEM, N=8-11, paired one-way ANOVA with Tukey test for multiple testing). Statistically significant comparisons are represented by *p < 0.05.

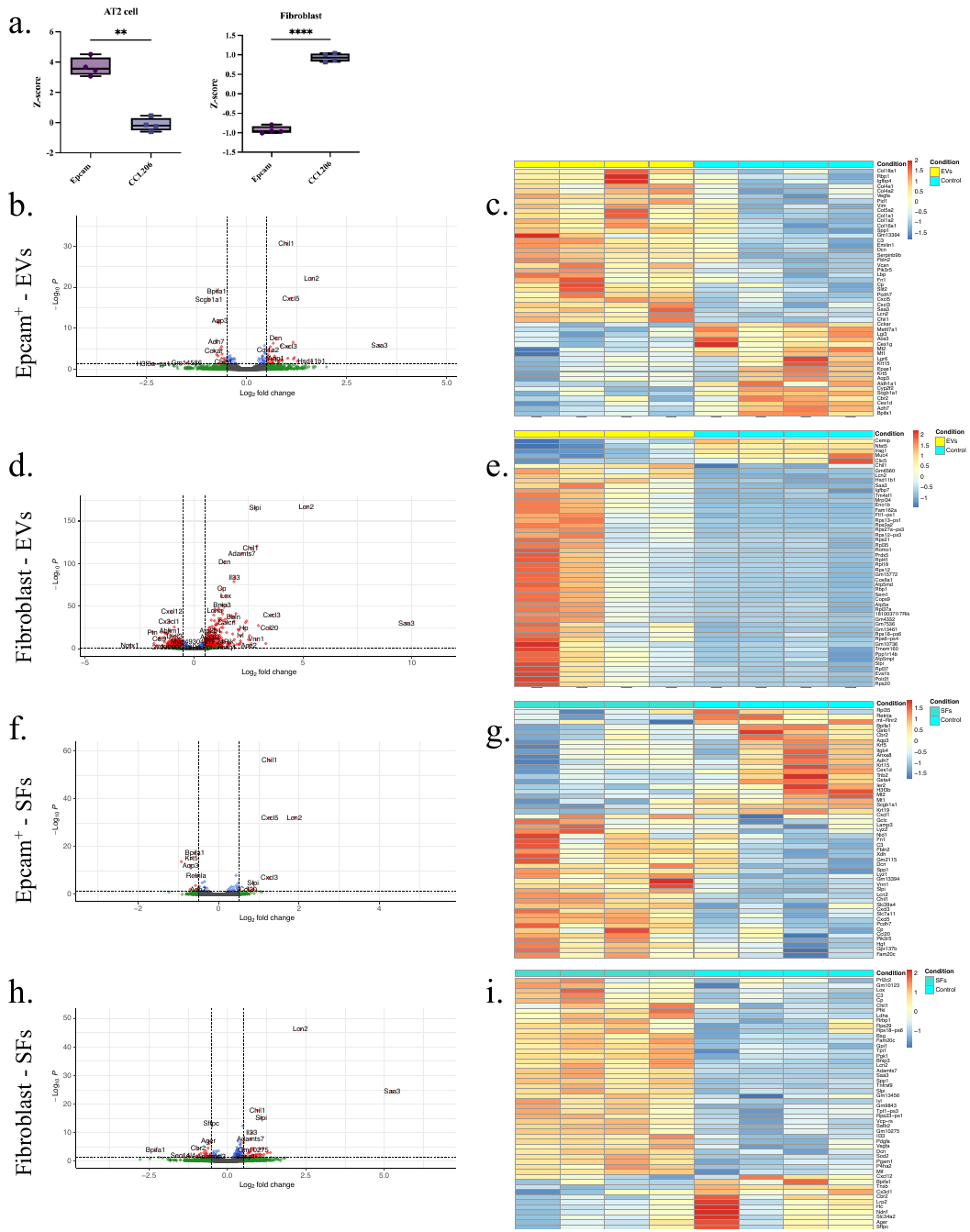

**Fig. S6. EVs and SFs induce transcriptional changes in Epcam^+^ cells and fibroblasts.**

**a** AT2 cell and fibroblast signature within control Epcam^+^ cell and fibroblast population (mean ± min/max, N=4, paired two-tailed T-test). **b** Volcano plot illustrating the response to EVs versus control in epithelial progenitor cells (cut-offs: padj<0.05 and logFold>1). **c** Heatmap with the top 50 significant genes up- or downregulated in epithelial progenitor cells with EVs. **d** Volcano plot illustrating the response to EVs versus control in fibroblasts (cut-offs: padj<0.05 and logFold>1). **e** Heatmap with the top 50 significant genes up- or downregulated in fibroblasts treated with EVs. **f** Volcano plot illustrating the response to SFs versus control in epithelial progenitor cells (cut-offs: padj<0.05 and logFold>1). **g** Heatmap with the top 50 significant genes up- or downregulated in epithelial progenitor cells with SFs. **h** Volcano plot illustrating the response to SFs versus control in fibroblasts (cut-offs: padj<0.05 and logFold>1). **i** Heatmap with the top 50 significant genes up- or downregulated in fibroblasts treated with SFs. Statistically significant comparisons are represented by **p < 0.01 and ****p < 0.0001.

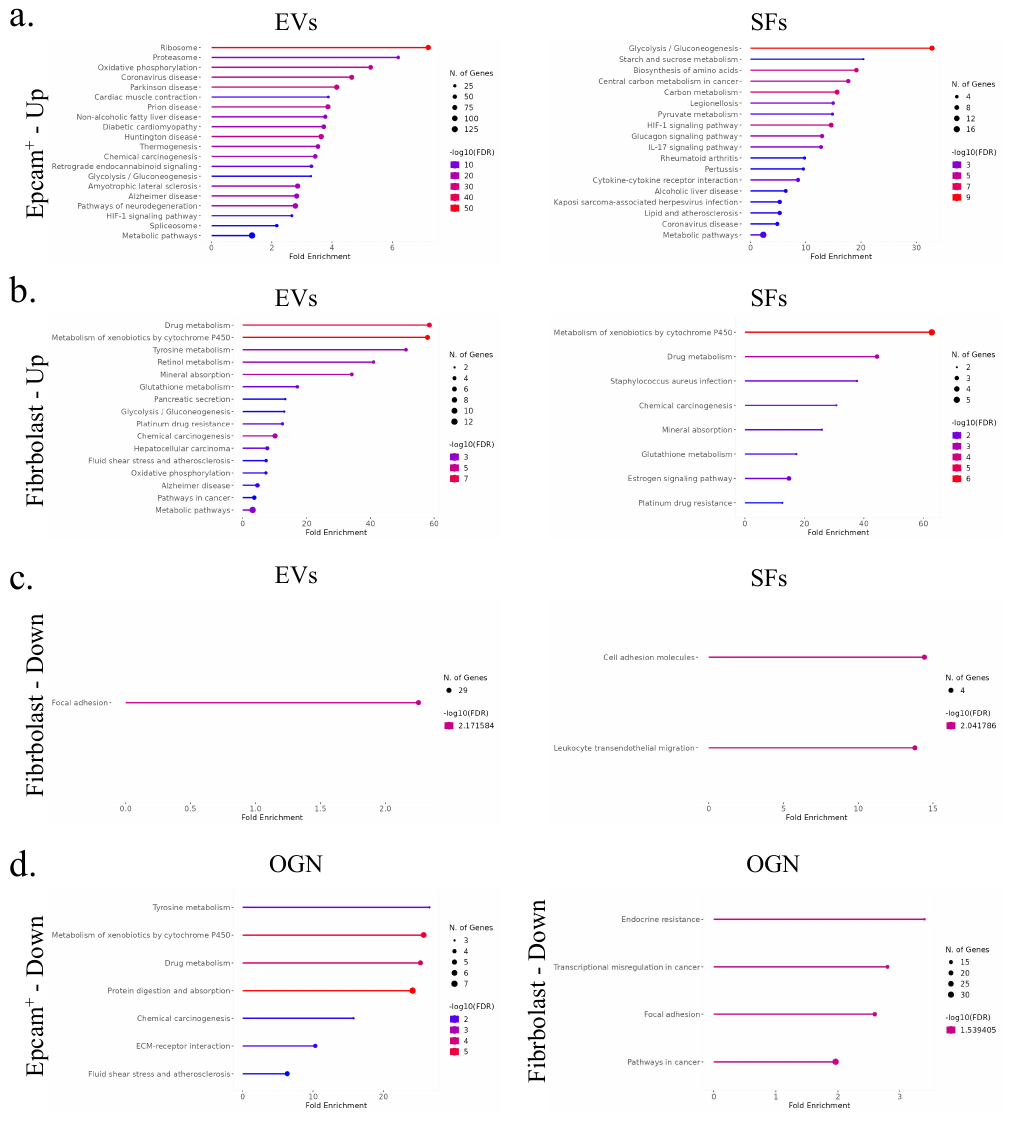

**Fig. S7. Gene set enrichment analysis in Epcam^+^ cells and fibroblasts upon treatment with EVs, SFs, or OGN.**

**a** The top upregulated pathways in epithelial progenitors treated with EVs or SFs. **b** The top upregulated pathways in fibroblasts treated with EVs or SFs. **c** The top-downregulated pathway(s) in fibroblasts treated with EVs or SFs. **d** The top downregulated pathways in epithelial progenitors or fibroblasts treated with OGN.

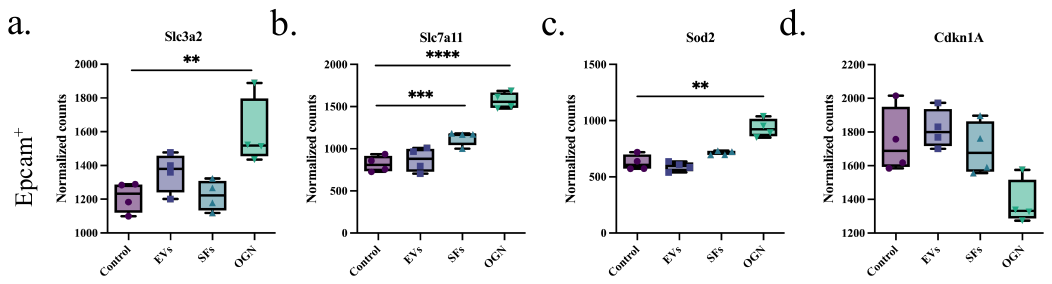

**Fig. S8. Osteoglycin increases the expression of protective genes in epithelial cells.**

**a-d** Normalized counts of heavy chain subunit of cystine and glutamine anti-transporter (*Slc3a2*), light chain subunit of cystine and glutamine anti-transporter (*Slc7a11*), superoxide dismutase 2 (*Sod2*), and senescence marker (*Cdkn1a*) upon treatment of epithelial progenitors with EVs, SFs, or OGN (mean ± min/max, N=4, paired one-way ANOVA with Tukey test for multiple testing). Statistically significant comparisons are represented by **p < 0.01, ***p < 0.001, and ****p < 0.0001.

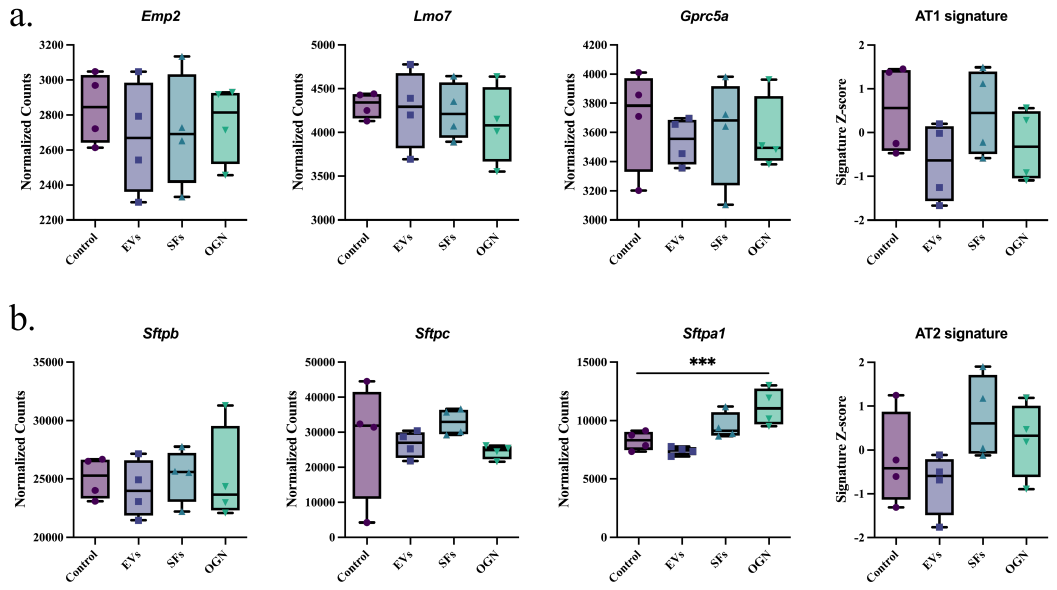

**Fig. S9. Osteoglycin increases the expression of protective genes in epithelial cells.**

**a** Normalized counts of markers for alveolar type 1 (AT1) cells (*Emp2*, *Lmo7*, and *Gprc5a*) and composite AT1 gene signature based on the top 10 AT1 marker genes upon treatment of epithelial progenitor cells with EVs, SFs, or OGN (mean ± min/max, N=4, paired one-way ANOVA with Tukey test for multiple testing). **b** Normalized counts of markers for alveolar type 2 (AT2) cells (*Sftpb*, *Sftpc*, and *Sftpa1*) and composite AT2 gene signature based on the top 10 AT2 marker genes upon treatment of epithelial progenitor cells with EVs, SFs, or OGN (mean ± min/max, N=4, paired one-way ANOVA with Tukey test for multiple testing). Statistically significant comparisons are represented by ***p < 0.001.

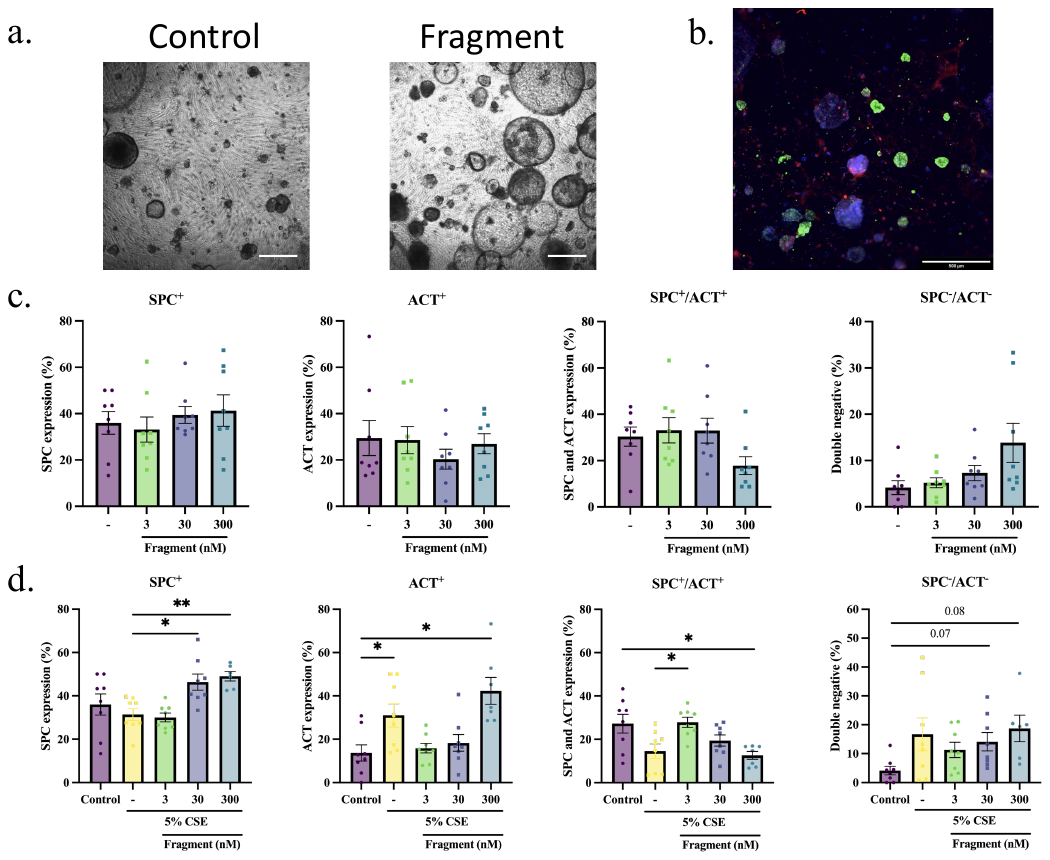

**Fig. S10. The active fragment of osteoglycin induces alveolar organoid formation.**

**a** Representative brightfield images of murine lung organoids (scale = 500 µm). **b** Representative immunofluorescence image of stained organoids treated with OGN fragment for airway-type organoids (acetylated α-tubulin, red), alveolar-type organoids (surfactant protein C, green), and Dapi (nuclei, blue) (scale = 500 µm). **c** Immunohistochemistry quantification for SPC^+^, ACT^+^, SPC^+^/ACT^+^, and SPC^-^/ACT^-^ organoids (mean ± SEM, N=8, paired one-way ANOVA with Tukey test for multiple testing). **d** Immunohistochemistry quantification for SPC^+^, ACT^+^, SPC^+^/ACT^+^, and SPC^-^/ACT^-^ organoids in the presence of CSE (mean ± SEM, N=8, paired one-way ANOVA with Tukey test for multiple testing). Statistically significant comparisons are represented by *p < 0.05 and **p < 0.01.

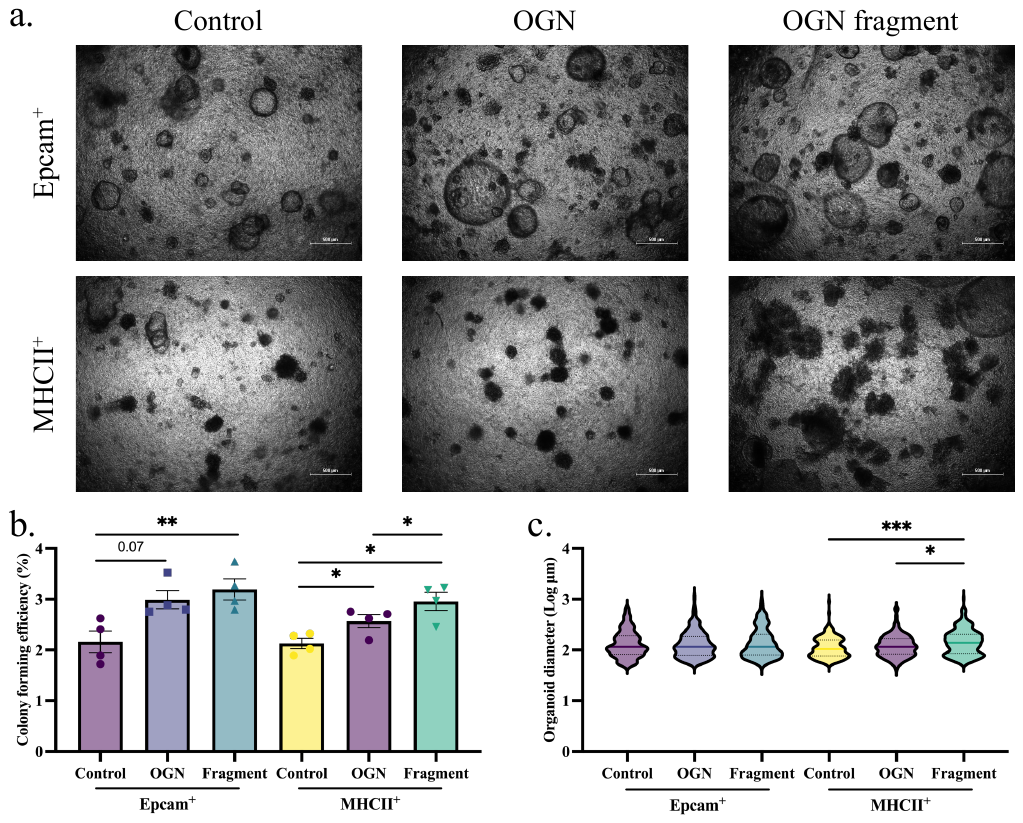

**Fig. S11. Osteoglycin and its active fragment induce organoid formation from MHCII^+^ and Epcam^+^ cells. A)** Representative brightfield images of murine lung organoids from Epcam^+^ cells and MHCII^+^ cells (scale = 500 µm). **B)** Colony forming efficiency of murine Epam^+^ and MHC^+^ cultures treated with OGN or its active fragment on day 14 (mean ± SEM, N=6, paired Friedman test). **C)** Log of organoid diameter of murine Epam^+^ and MHC^+^ cultures treated with OGN or its active fragment on day 14 (median is shown, N=6, Kolmogorov-Smirnov test (after Bonferroni correction: α = 0.025)). Statistically significant comparisons are represented by *p < 0.05, **p < 0.01, and ***p < 0.001.

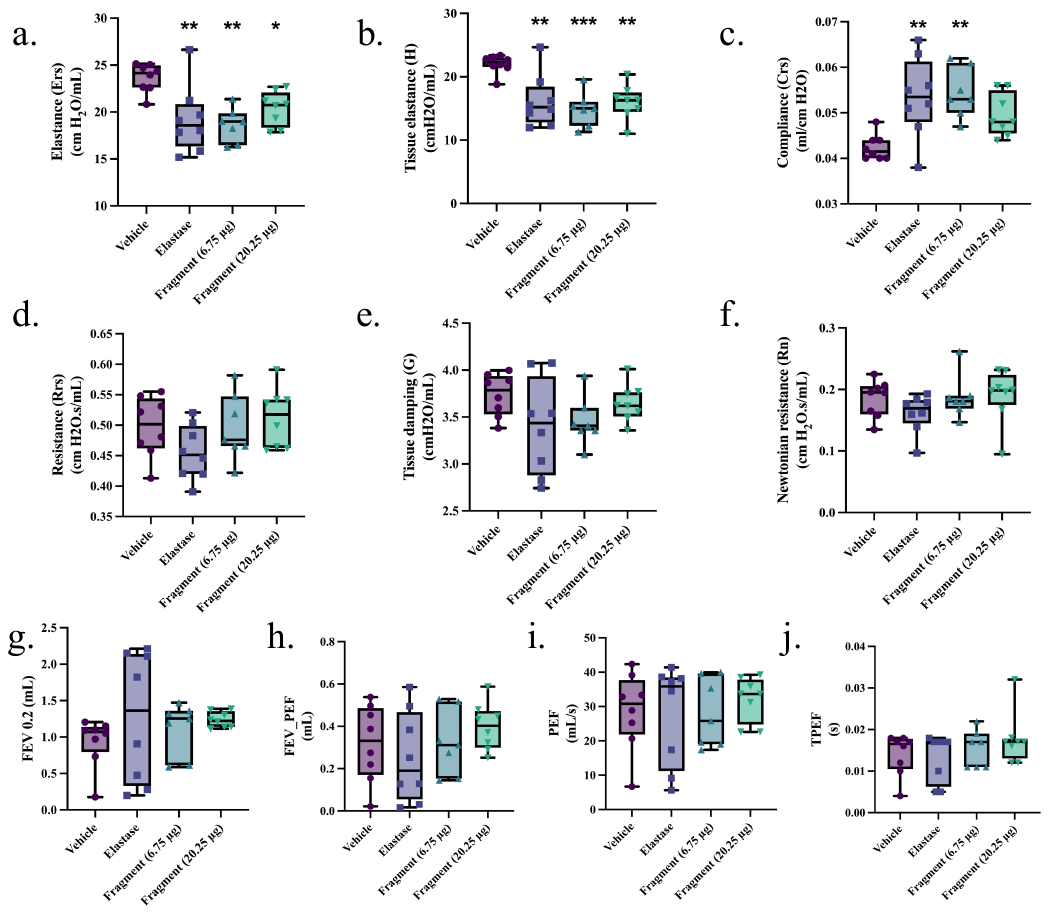

**Fig. S12. Lung tissue characteristics and lung function parameters in murine elastase-induced lung injury model.**

**a-f** Lung tissue characteristics: elastance, tissue elastance, compliance, resistance, tissue damping, and Newtonian resistance as measured with the FlexiVent (median ± min/max data point, N=7-8, One-Way ANOVA followed by Sidak’s multiple comparison). **g-j** Forced expiratory volume at 0.2 seconds (FEV0.2), forced expiratory volume at peak expiratory flow (FEV_PEF), peak expiratory flow (PEF), time to reach peak expiratory flow (TPEF) as measured with the FlexiVent (median ± min/max data point, N=7-8, One-Way ANOVA followed by Sidak’s multiple comparison). Statistically significant comparisons are represented by *p < 0.05, **p < 0.01, and ***p < 0.001.

**Table S1. Assessment of proteomics analysis based on MISEV guidelines.** Data are categorized based on the MISEV 2018 guidelines. Green-colored rows represent higher relative abundance in EV samples, blue rows represent higher relative abundance in SF samples, and red rows represent ratios that could not be calculated. N/A, not applicable.

| **Class** | **Sub-class** | **Names** | **Proteins** | **Presence in EV-samples** | **Presence in SF-samples** | **Ratio of average abundancy (EV/SF)** |
| --- | --- | --- | --- | --- | --- | --- |
| **Transmembrane or GPI-anchored proteins associated to plasma membrane and/or endosomes** | **Non-tissue specific** | Tetraspanins | CD63 | 4/4 | 0/4 | N/A |
|  |  |  | CD81 | 4/4 | 4/4 | 79,28 |
|  |  |  | CD82 | 4/4 | 4/4 | 64,59 |
|  |  | Multi-pass membrane proteins (GNA*) | CD47 | 4/4 | 1/4 | 61,88 |
|  |  |  | GNAI2 | 4/4 | 4/4 | 8,15 |
|  |  |  | GNAI3 | 1/4 | 0/4 | N/A |
|  |  |  | GNA11 | 3/4 | 1/4 | 26,10 |
|  |  |  | GNAS | 3/4 | 3/4 | 43,53 |
|  |  | MHC-class 1 | HLA-A | 4/4 | 4/4 | 40,50 |
|  |  |  | HLA-B | 3/4 | 4/4 | 7,18 |
|  |  |  | HLA-C | 3/4 | 4/4 | 1,42 |
|  |  |  | H2-K | 0/4 | 0/4 | N/A |
|  |  |  | H2-D | 0/4 | 0/4 | N/A |
|  |  |  | H2Q | 0/4 | 0/4 | N/A |
|  |  | Integrins (ITGA*/ITGB*) | ITGA1 | 4/4 | 4/4 | 24,11 |
|  |  |  | ITGA2 | 4/4 | 3/4 | 23,86 |
|  |  |  | ITGA3 | 4/4 | 4/4 | 35,90 |
|  |  |  | ITGA5 | 4/4 | 4/4 | 6,75 |
|  |  |  | ITGAV | 4/4 | 4/4 | 35,09 |
|  |  |  | ITGB1 | 4/4 | 4/4 | 34,79 |
|  |  |  | ITGB3 | 4/4 | 1/4 | 3,18 |
|  |  | Transferrin receptor | TFR2 | 0/4 | 0/4 | N/A |
|  |  | Lysosomal-associated membrane protein | LAMP1 | 4/4 | 4/4 | 20,20 |
|  |  |  | LAMP2 | 4/4 | 4/4 | 17,14 |
|  |  | Heparan sulfate proteoglycans (SDC*) | SDC1 | 4/4 | 4/4 | 32,55 |
|  |  |  | SDC2 | 4/4 | 1/4 | 8,81 |
|  |  |  | SDC4 | 4/4 | 4/4 | 9,03 |
|  |  |  | SDCBP | 4/4 | 4/4 | 51,48 |
|  |  | EMMPRIN | BSG | 4/4 | 4/4 | 23,22 |
|  |  | ADAM metallopeptidase domain 10 | ADAM10 | 4/4 | 4/4 | 7,77 |
|  |  | GPI-anchored 5’nucleotidase | CD73 / NT4E | 0/4 | 0/4 | N/A |
|  |  | Complement-binding proteins | CD55 | 4/4 | 3/4 | 6,43 |
|  |  |  | CD59 | 4/4 | 4/4 | 16,53 |
|  |  | Sonic Hedgehog | SHH | 0/4 | 0/4 | N/A |
|  | **Cell or Tissue specific** | Epithelial cell | TSPAN8 | 0/4 | 0/4 | N/A |
|  |  | Leukocytes | CD37 | 0/4 | 0/4 | N/A |
|  |  |  | CD53 | 0/4 | 0/4 | N/A |
|  |  | Absent from NK and B cells | CD9 | 4/4 | 4/4 | 46,19 |
|  |  | Endothelial cells | PECAM1 | 0/4 | 0/4 | N/A |
|  |  | Breast cancer | ERBB2 | 0/4 | 0/4 | N/A |
|  |  | Epithelial cells | EPCAM | 0/4 | 0/4 | N/A |
|  |  | Mesenchymal Stromal Cels | CD90 / THY1 | 4/4 | 3/4 | 69,01 |
|  |  | Immune cells | CD45 / PTPRC | 0/4 | 0/4 | N/A |
|  |  | Platelets | CD41 / ITGA2B | 0/4 | 0/4 | N/A |
|  |  |  | CD42a / GP9 | 0/4 | 0/4 | N/A |
|  |  | Red blood cells | GYPA | 0/4 | 0/4 | N/A |
|  |  | Monocytes | CD14 | 0/4 | 0/4 | N/A |
|  |  | MHC class II | HLA-DR | 0/4 | 0/4 | N/A |
|  |  |  | HLA-DP | 0/4 | 0/4 | N/A |
|  |  |  | HLA-DQ | 0/4 | 0/4 | N/A |
|  |  |  | H2-A* | 0/4 | 0/4 | N/A |
|  |  | T cells | CD3* | 0/4 | 0/4 | N/A |
|  |  | Neurons | AChE-S | 0/4 | 0/4 | N/A |
|  |  |  | APP | 4/4 | 4/4 | 0,53 |
|  |  | Erythrocytes | AChE-E | 0/4 | 0/4 | N/A |
|  |  | Multi-drug resistance-associated protein | ABCC1 | 1/4 | 0/4 | N/A |
| **Cytosolic proteins recovered in EVs** | **With lipid or membrane protein-binding ability** | ESCRT-I/II/III (CHMP*) | TSG101 | 4/4 | 2/4 | 12,10 |
|  |  |  | CHMP1B | 3/4 | 1/4 | 44,63 |
|  |  |  | CHMP2A | 3/4 | 0/4 | N/A |
|  |  |  | CHMP4B | 3/4 | 0/4 | N/A |
|  |  |  | CHMP5 | 0/4 | 1/4 | N/A |
|  |  | Accessory proteins | ALIX | 0/4 | 0/4 | N/A |
|  |  |  | VPS4A | 0/4 | 0/4 | N/A |
|  |  |  | VPS4B | 0/4 | 0/4 | N/A |
|  |  |  | ARRDC1 | 0/4 | 1/4 | N/A |
|  |  | Flotillins | FLOT1 | 4/4 | 3/4 | 32,85 |
|  |  |  | FLOT2 | 3/4 | 2/4 | 33,23 |
|  |  | Caveolins (CAV*) | CAV1 | 0/4 | 2/4 | N/A |
|  |  |  | CAVIN1 | 0/4 | 4/4 | N/A |
|  |  | Eps15 Homology Domain (EDH*) | EDH1 | 3/4 | 4/4 | 2,83 |
|  |  |  | EDH2 | 4/4 | 4/4 | 9,07 |
|  |  |  | EHD3 | 4/4 | 4/4 | 6,93 |
|  |  |  | EDH4 | 1/4 | 3/4 | 10,83 |
|  |  | Ras Homolog Family Member A | RHOA | 2/4 | 4/4 | 0,90 |
|  |  | Annexins (ANX*) | ANXA1 | 4/4 | 4/4 | 2,45 |
|  |  |  | ANXA2 | 4/4 | 4/4 | 3,84 |
|  |  |  | ANXA3 | 0/4 | 1/4 | N/A |
|  |  |  | ANXA4 | 3/4 | 1/4 | 22,22 |
|  |  |  | ANXA5 | 4/4 | 4/4 | 0,31 |
|  |  |  | ANXA6 | 4/4 | 4/4 | 9,76 |
|  |  |  | ANXA7 | 2/4 | 1/4 | 3,82 |
|  |  |  | ANXA11 | 4/4 | 3/4 | 17,93 |
|  |  | Heat shock proteins | HSC70 / HSPA8 | 4/4 | 4/4 | 1,84 |
|  |  |  | HSP84 / HSP90AB1 | 4/4 | 4/4 | 0,25 |
|  |  | ADP-ribosylation factor 6 | ARF6 | 0/4 | 0/4 | N/A |
|  |  | Syntenin | SDCBP | 4/4 | 4/4 | 51,48 |
|  |  | Microtubule-associated Tau (neurons) | MAPT | 0/4 | 0/4 | N/A |
|  | **Promiscuous incorporation in EVs** | Heat shock protein | HSP70 / HSPA1A | 4/4 | 4/4 | 3,61 |
|  |  | Cytoskeleton (ACT* / TUB*) | ACTN1 | 4/4 | 4/4 | 0,22 |
|  |  |  | ACTN4 | 4/4 | 4/4 | 0,20 |
|  |  |  | ACTB | 4/4 | 4/4 | 17,75 |
|  |  |  | ACTBL2 | 4/4 | 4/4 | 39,31 |
|  |  |  | ACTR2 | 3/4 | 4/4 | 0,93 |
|  |  |  | ACTR3 | 4/4 | 4/4 | 0,41 |
|  |  |  | ACTR5 | 2/4 | 0/4 | N/A |
|  |  |  | ACTR1A | 4/4 | 4/4 | 1,61 |
|  |  |  | ACTR1B | 0/4 | 4/4 | N/A |
|  |  |  | ACTA2 | 4/4 | 4/4 | 1,07 |
|  |  |  | ACTG1 | 4/4 | 4/4 | 0,84 |
|  |  |  | ACTL6A | 0/4 | 4/4 | N/A |
|  |  |  | TUBA1A | 4/4 | 4/4 | 0,87 |
|  |  |  | TUBA1B | 4/4 | 4/4 | 1,51 |
|  |  |  | TUBA1C | 4/4 | 4/4 | 5,11 |
|  |  |  | TUBA4A | 4/4 | 4/4 | 8,70 |
|  |  |  | TUBAL3 | 0/4 | 3/4 | N/A |
|  |  |  | TUBB | 4/4 | 4/4 | 1,75 |
|  |  |  | TUBB1 | 1/4 | 3/4 | 1,41 |
|  |  |  | TUBB2A | 0/4 | 4/4 | N/A |
|  |  |  | TUBB2B | 3/4 | 4/4 | 3,29 |
|  |  |  | TUBB3 | 4/4 | 4/4 | 1,31 |
|  |  |  | TUBB4B | 4/4 | 4/4 | 1,62 |
|  |  |  | TUBB6 | 4/4 | 4/4 | 4,78 |
|  |  |  | TUBG1;TUBG2 | 0/4 | 1/4 | N/A |
|  |  | Enzymes | GAPDH | 4/4 | 4/4 | 1,91 |
| **Major components of non-EV co-isolated structures** | **Lipoproteins (produced by liver, abundant in plasma, serum)** | Lipoproteins | APOA1 | 4/4 | 4/4 | 0,31 |
|  |  |  | APOA2 | 0/4 | 0/4 | N/A |
|  |  |  | APOB | 4/4 | 4/4 | 1,08 |
|  |  |  | APOB100 | 0/4 | 0/4 | N/A |
|  |  | Albumin | ALB | 4/4 | 4/4 | 1,29 |
|  | **Protein and protein/nucleic acid aggregates** | Tamm-Horsfall protein | UMOD | 0/4 | 0/4 | N/A |
|  |  | Ribosomal proteins (RPL* / RPS*) | RPL10 | 3/4 | 4/4 | 1,02 |
|  |  |  | RPL10A | 3/4 | 4/4 | 0,43 |
|  |  |  | RPL11 | 4/4 | 4/4 | 1,70 |
|  |  |  | RPL12 | 2/4 | 4/4 | 0,17 |
|  |  |  | RPL13 | 0/4 | 4/4 | N/A |
|  |  |  | RPL13A | 0/4 | 4/4 | N/A |
|  |  |  | RPL14 | 0/4 | 3/4 | N/A |
|  |  |  | RPL15 | 4/4 | 4/4 | 2,71 |
|  |  |  | RPL17 | 1/4 | 4/4 | 1,04 |
|  |  |  | RPL18 | 3/4 | 4/4 | 2,76 |
|  |  |  | RPL18A | 3/4 | 4/4 | 7,12 |
|  |  |  | RPL19 | 1/4 | 4/4 | 0,35 |
|  |  |  | RPL22 | 0/4 | 4/4 | N/A |
|  |  |  | RPL23 | 1/4 | 4/4 | 2,57 |
|  |  |  | RPL23A | 0/4 | 2/4 | N/A |
|  |  |  | RPL24 | 0/4 | 3/4 | N/A |
|  |  |  | RPL26 | 0/4 | 4/4 | N/A |
|  |  |  | RPL27 | 2/4 | 4/4 | 4,56 |
|  |  |  | RPL27A | 1/4 | 4/4 | 14,20 |
|  |  |  | RPL28 | 3/4 | 4/4 | 0,89 |
|  |  |  | RPL29 | 2/4 | 4/4 | 3,64 |
|  |  |  | RPL3 | 4/4 | 4/4 | 3,01 |
|  |  |  | RPL30 | 0/4 | 4/4 | N/A |
|  |  |  | RPL31 | 0/4 | 4/4 | N/A |
|  |  |  | RPL32 | 0/4 | 2/4 | N/A |
|  |  |  | RPL34 | 0/4 | 3/4 | N/A |
|  |  |  | RPL35 | 0/4 | 1/4 | N/A |
|  |  |  | RPL35A | 4/4 | 4/4 | 3,57 |
|  |  |  | RPL36 | 2/4 | 4/4 | 0,69 |
|  |  |  | RPL36A | 2/4 | 1/4 | 3,95 |
|  |  |  | RPL37A | 1/4 | 2/4 | 3,07 |
|  |  |  | RPL38 | 0/4 | 4/4 | N/A |
|  |  |  | RPL4 | 1/4 | 4/4 | 1,42 |
|  |  |  | RPL5 | 3/4 | 4/4 | 0,74 |
|  |  |  | RPL6 | 4/4 | 4/4 | 1,63 |
|  |  |  | RPL7 | 0/4 | 3/4 | N/A |
|  |  |  | RPL7A | 1/4 | 4/4 | 1,18 |
|  |  |  | RPL8 | 4/4 | 4/4 | 2,18 |
|  |  |  | RPL9 | 0/4 | 3/4 | N/A |
|  |  |  | RPLP0 | 4/4 | 4/4 | 0,48 |
|  |  |  | RPLP1 | 0/4 | 4/4 | N/A |
|  |  |  | RPLP2 | 0/4 | 3/4 | N/A |
|  |  |  | RPS10 | 0/4 | 4/4 | N/A |
|  |  |  | RPS11 | 4/4 | 4/4 | 2,83 |
|  |  |  | RPS13 | 1/4 | 4/4 | 1,83 |
|  |  |  | RPS14 | 4/4 | 4/4 | 3,32 |
|  |  |  | RPS15A | 1/4 | 4/4 | 0,52 |
|  |  |  | RPS16 | 3/4 | 4/4 | 2,97 |
|  |  |  | RPS17 | 0/4 | 3/4 | N/A |
|  |  |  | RPS18 | 4/4 | 4/4 | 2,54 |
|  |  |  | RPS19 | 0/4 | 1/4 | N/A |
|  |  |  | RPS2 | 4/4 | 4/4 | 2,66 |
|  |  |  | RPS20 | 2/4 | 4/4 | 2,63 |
|  |  |  | RPS21 | 0/4 | 3/4 | N/A |
|  |  |  | RPS23 | 2/4 | 4/4 | 0,84 |
|  |  |  | RPS25 | 3/4 | 4/4 | 2,24 |
|  |  |  | RPS26 | 3/4 | 4/4 | 7,64 |
|  |  |  | RPS27L | 4/4 | 4/4 | 1,68 |
|  |  |  | RPS28 | 0/4 | 4/4 | N/A |
|  |  |  | RPS3 | 4/4 | 4/4 | 2,17 |
|  |  |  | RPS3A | 1/4 | 4/4 | 4,86 |
|  |  |  | RPS4X | 4/4 | 4/4 | 4,49 |
|  |  |  | RPS5 | 3/4 | 4/4 | 3,13 |
|  |  |  | RPS6 | 3/4 | 4/4 | 3,13 |
|  |  |  | RPS7 | 1/4 | 4/4 | 11,85 |
|  |  |  | RPS8 | 4/4 | 4/4 | 2,94 |
|  |  |  | RPS9 | 3/4 | 4/4 | 3,28 |
|  |  |  | RPSA | 4/4 | 4/4 | 1,42 |
| **Transmembrane, lipid bound and soluble proteins associated to other intracellular compartments than PM/endosomes** | **Nucleus** | Histones | HIST1H2BK;H2BFS;HIST1H2BD;H2BC4;  HIST2H2BF;HIST1H2BH;HIST1H2BN;  HIST1H2BM;HIST1H2BL | 3/4 | 4/4 | 4,04 |
|  |  |  | HIST1H2BJ;HIST1H2BO;  HIST1H2BB;HIST2H2BE | 4/4 | 4/4 | 3,31 |
|  |  |  | HIST1H1B | 2/4 | 2/4 | 11,31 |
|  |  |  | HIST1H1C | 2/4 | 3/4 | 2,12 |
|  |  |  | HIST1H1E | 3/4 | 4/4 | 2,47 |
|  |  | Lamin A | LMNA | 4/4 | 4/4 | 0,32 |
|  | **Mitochondria** | Inner Membrane Mitochondrial Protein | IMMT | 0/4 | 0/4 | N/A |
|  |  | Cytochrome C1 | CYC1 | 0/4 | 0/4 | N/A |
|  |  | Translocase Of Outer Mitochondrial Membrane 20 | TOMM20 | 0/4 | 0/4 | N/A |
|  | **Secretory pathway** | Calnexin | CANX | 4/4 | 4/4 | 1,56 |
|  |  | Heat Shock Protein 90 Beta Family Member 1 | HSP90B1 | 4/4 | 4/4 | 0,11 |
|  |  | Heat Shock Protein Family A Member 5 | HSPA5 | 4/4 | 4/4 | 0,76 |
|  |  | Golgin A2 | GOLGA2 | 0/4 | 0/4 | N/A |
|  | **Other** | Autophagy related 9A | ATG9A | 0/4 | 0/4 | N/A |
|  |  | Actins | ACTN1 | 4/4 | 4/4 | 0,22 |
|  |  |  | ACTN4 | 4/4 | 4/4 | 0,20 |
|  |  | Cytokeratin 18 | KRT18 | 1/4 | 4/4 | 5,05 |
| **Secreted proteins recovered with EVs** | **Cytokines and growth factors** | Transforming Growth Factor Beta 1 | TGFB1 | 4/4 | 4/4 | 0,60 |
|  |  | Transforming Growth Factor Beta 2 | TGFB2 | 0/4 | 4/4 | N/A |
|  |  | Interferon Gamma | IFNG | 0/4 | 0/4 | N/A |
|  |  | Vascular Endothelial Growth Factor A | VEGFA | 0/4 | 0/4 | N/A |
|  |  | Fibroblast Growth Factor 1 | FGF1 | 0/4 | 0/4 | N/A |
|  |  | Fibroblast Growth Factor 2 | FGF2 | 0/4 | 0/4 | N/A |
|  |  | Platelet-Derived Growth Factor | PDGFC | 2/4 | 4/4 | 0,16 |
|  |  |  | PDGFD | 1/4 | 4/4 | 0,45 |
|  |  | Epidermal Growth Factor | EGF | 0/4 | 0/4 | N/A |
|  |  | Interleukin-6 | IL6 | 0/4 | 3/4 | N/A |
|  | **Adhesion and extracellular matrix proteins** | Fibronectin 1 | FN1 | 4/4 | 4/4 | 0,68 |
|  |  | Collagen (COL*) | COL1A1 | 4/4 | 4/4 | 15,96 |
|  |  |  | COL1A2 | 4/4 | 4/4 | 8,60 |
|  |  |  | COL2A1 | 4/4 | 4/4 | 10,22 |
|  |  |  | COL3A1 | 4/4 | 4/4 | 10,62 |
|  |  |  | COL4A1 | 4/4 | 4/4 | 1,96 |
|  |  |  | COL4A2 | 4/4 | 4/4 | 1,43 |
|  |  |  | COL5A1 | 4/4 | 4/4 | 3,29 |
|  |  |  | COL5A2 | 4/4 | 4/4 | 2,13 |
|  |  |  | COL6A1 | 4/4 | 4/4 | 4,84 |
|  |  |  | COL6A2 | 4/4 | 4/4 | 6,59 |
|  |  |  | COL6A3 | 4/4 | 4/4 | 6,29 |
|  |  |  | COL7A1 | 4/4 | 4/4 | 9,20 |
|  |  |  | COL8A1 | 0/4 | 4/4 | N/A |
|  |  |  | COL10A1 | 0/4 | 4/4 | N/A |
|  |  |  | COL14A1 | 1/4 | 3/4 | 0,50 |
|  |  |  | COL15A1 | 4/4 | 4/4 | 17,70 |
|  |  |  | COL18A1 | 4/4 | 4/4 | 16,45 |
|  |  | Milk Fat Globule EGF | MFGE8 | 4/4 | 4/4 | 29,79 |
|  |  | Galectin3-binding protein | LGALS3BP | 4/4 | 4/4 | 1,96 |
|  |  | CD5 molecule like | CD5L | 0/4 | 0/4 | N/A |
|  |  | Fetuin-A | AHSG | 4/4 | 4/4 | 0,15 |

**Table S2. Relative abundance of proteins present in all three replicates of lung fibroblast-derived EVs and SFs.**

Data are organized alphabetically.

| **Protein ID** | **EV1** | **EV2** | **EV3** | **EV4** | **SF1** | **SF2** | **SF3** | **SF4** |
| --- | --- | --- | --- | --- | --- | --- | --- | --- |
| A2M | 2099630,25 | 3561911,25 | 3887590,75 | 11037333,00 | 4325883,50 | 8218129,50 | 2444535,00 | 5136352,50 |
| AARS | 45520,28 | 29067,80 | 44199,16 | 219898,33 | 64920,82 | 67642,19 | 54782,34 | 41569,97 |
| ABI3BP | 131090,08 | 333473,84 | 198256,55 | 108285,95 | 11211,62 | 10576,46 | 9391,09 | 16823,61 |
| ACAN | 78561,59 | 236119,98 | 139433,00 | 264260,13 | 4769,55 | 30119,34 | 6633,27 | 15694,53 |
| ACLY | 14380,67 | 16095,07 | 22420,34 | 8043,20 | 23555,51 | 24984,97 | 27074,42 | 17996,67 |
| ACO1 | 18221,31 | 15541,75 | 21846,88 | 14676,95 | 18126,09 | 10717,63 | 15789,28 | 14670,48 |
| ACTA2 | 132489,91 | 50613,29 | 219055,69 | 47841,07 | 117933,25 | 96085,38 | 101586,84 | 104588,90 |
| ACTB | 627145,81 | 257649,64 | 1237549,75 | 281830,84 | 32894,02 | 42098,53 | 30355,16 | 30079,81 |
| ACTBL2 | 274252,28 | 880207,63 | 5528283,50 | 1249316,63 | 49245,09 | 75310,79 | 67726,20 | 9521,23 |
| ACTG1 | 5996162,50 | 3408193,75 | 5867516,00 | 2469540,50 | 5051828,00 | 5256116,50 | 4822049,00 | 5995818,50 |
| ACTN1 | 78284,71 | 60523,37 | 49780,89 | 130528,66 | 379590,56 | 317903,84 | 337940,84 | 391891,47 |
| ACTN4 | 51512,30 | 38289,74 | 17795,38 | 85878,77 | 265604,44 | 201501,61 | 245240,31 | 233918,88 |
| ACTR1A | 23130,74 | 28878,35 | 43511,00 | 17324,40 | 19486,26 | 17619,81 | 19169,27 | 13764,12 |
| ACTR3 | 29861,41 | 40719,37 | 77481,86 | 42643,46 | 126379,69 | 131583,19 | 105974,74 | 99698,53 |
| ADAM10 | 128270,05 | 130491,84 | 161467,98 | 56783,53 | 11366,87 | 19437,78 | 14558,20 | 16030,28 |
| ADAM9 | 355547,38 | 385293,28 | 1114406,25 | 178499,39 | 30161,77 | 11447,38 | 20862,21 | 19568,21 |
| AEBP1 | 27343,83 | 35479,04 | 30423,61 | 22162,75 | 84475,73 | 85787,25 | 78279,95 | 98195,94 |
| AFP | 47162,96 | 52656,71 | 122633,81 | 220912,84 | 81355,23 | 418369,72 | 126401,63 | 429515,19 |
| AGRN | 1276232,63 | 1640076,25 | 3225285,25 | 754915,38 | 45298,28 | 46000,69 | 58997,86 | 48876,46 |
| AHCY | 35546,10 | 50453,98 | 80891,75 | 57181,03 | 37167,09 | 64313,14 | 47092,06 | 40797,64 |
| AHNAK | 48755,07 | 40597,71 | 113925,09 | 78192,22 | 13226,15 | 11903,38 | 19972,17 | 6700,69 |
| AHSG | 383627,03 | 1022947,75 | 1474925,13 | 1212249,88 | 3481528,00 | 11545314,00 | 5963862,50 | 6660407,50 |
| AKR1B1 | 27487,03 | 21843,71 | 15901417,00 | 12191,29 | 41160,56 | 24742,81 | 35355,12 | 27756,29 |
| ALB | 5115764,00 | 5285184,50 | 12791583,00 | 6074637,50 | 1859396,88 | 6415930,50 | 2538101,25 | 11819574,00 |
| ALDOA | 118329,01 | 79962,72 | 119398,11 | 143480,64 | 1102568,00 | 1039462,50 | 892007,94 | 915898,44 |
| ALDOB | 17017,42 | 14425,96 | 28331,81 | 20482,96 | 18384,94 | 55172,79 | 25894,12 | 8205,29 |
| ALDOC | 36886,67 | 64406,65 | 38053,58 | 77837,77 | 281794,44 | 347672,59 | 270233,13 | 262954,81 |
| ANGPTL2 | 115664,31 | 58111,42 | 59166,39 | 47775,11 | 122112,41 | 93218,85 | 108917,36 | 79463,13 |
| ANXA1 | 28896,60 | 25486,69 | 54253,58 | 15028,87 | 8032,30 | 6507,61 | 30019,41 | 5888,68 |
| ANXA2 | 194037,83 | 177526,31 | 346407,03 | 221130,92 | 46079,71 | 55998,19 | 112232,04 | 30052,87 |
| ANXA5 | 158199,55 | 198359,81 | 185022,47 | 101664,46 | 603451,88 | 340482,94 | 710597,94 | 394433,91 |
| ANXA6 | 78632,91 | 95104,44 | 194642,77 | 60279,82 | 2670,23 | 33453,84 | 3578,16 | 4216,66 |
| AP2B1 | 20365,63 | 26987,98 | 7314,20 | 29918,89 | 104224,44 | 117721,96 | 92789,34 | 38535,96 |
| APOA1 | 106938,38 | 70101,27 | 415628,72 | 136515,98 | 214081,95 | 1253856,25 | 326339,59 | 526419,13 |
| APOB | 40671,66 | 66536,41 | 75052,77 | 108421,72 | 25302,57 | 152690,03 | 18894,65 | 71595,28 |
| APOE | 102735,37 | 108564,34 | 111410,98 | 92323,16 | 107627,71 | 34174,58 | 27694,89 | 58710,93 |
| APOM | 43072,67 | 66114,14 | 270346,69 | 125625,44 | 29204,08 | 161480,94 | 43000,00 | 81011,75 |
| APP | 39950,48 | 53437,28 | 40860,22 | 54862,21 | 69624,56 | 83923,70 | 137897,17 | 64812,29 |
| ARF3;ARF1 | 30331,58 | 31775,98 | 42744,58 | 13214,75 | 11493,16 | 15731,01 | 7812,85 | 19606,56 |
| ARG1 | 60142,30 | 25196,48 | 32991,57 | 96158,14 | 1670,99 | 6439,30 | 2491,21 | 680,91 |
| ATP1A1 | 40019,53 | 37751,00 | 57878,75 | 24951,81 | 227,14 | 1726,25 | 1492,54 | 747,04 |
| ATP5F1A | 28711,89 | 35563,91 | 100558,95 | 46475,95 | 9348,22 | 24675,52 | 12367,00 | 2805,69 |
| BASP1 | 19610,62 | 11027,12 | 22614,49 | 7723,15 | 7979,57 | 3845,31 | 9922,34 | 3460,37 |
| BGN | 412704,06 | 518113,09 | 764441,44 | 1146070,50 | 547848,56 | 668927,13 | 613515,25 | 895828,38 |
| BLVRA | 36308,30 | 27180,73 | 69995,20 | 32212,17 | 11205,36 | 6349,54 | 10851,09 | 8331,17 |
| BSG | 53060,43 | 90242,89 | 126127,32 | 42007,57 | 3182,31 | 4052,23 | 3334,41 | 2846,40 |
| C1R | 144288,08 | 117217,05 | 66795,02 | 222262,98 | 806167,81 | 598402,69 | 841374,13 | 783461,56 |
| C1RL | 43801,36 | 52309,96 | 38800,45 | 99892,75 | 375756,50 | 174762,13 | 307418,84 | 240119,75 |
| C1S | 92881,80 | 78341,88 | 75461,33 | 186908,94 | 526956,88 | 383660,56 | 408760,56 | 613400,44 |
| C3 | 89909,84 | 252100,36 | 284447,91 | 499385,34 | 154483,92 | 626021,25 | 209112,08 | 450459,97 |
| CALM1;CALM2;CALM3 | 56121,35 | 30160,42 | 44534,13 | 21378,73 | 20940,58 | 10706,06 | 18701,11 | 3393,69 |
| CALML5 | 543621,56 | 284966,56 | 763198,81 | 1714569,75 | 26120,99 | 15316,84 | 11391,38 | 1421,58 |
| CALR | 55967,93 | 30251,35 | 68457,56 | 48135,84 | 390050,97 | 259630,80 | 429670,59 | 307939,59 |
| CANX | 6641,90 | 1299,83 | 12451,93 | 8144,98 | 4734,23 | 4049,17 | 6372,16 | 3151,37 |
| CAPN1 | 11559,10 | 1235342,75 | 211486,34 | 45473,26 | 24791,98 | 32891,52 | 19860,18 | 20945,51 |
| CAPNS1 | 579490,38 | 17538,51 | 137237,83 | 97248,30 | 29354,30 | 11775,50 | 15984,69 | 12633,37 |
| CAPZA1 | 18720,80 | 23547,39 | 53307,68 | 83759,90 | 49354,26 | 27919,26 | 53452,10 | 32478,54 |
| CAPZA2 | 17578,53 | 19362,60 | 28612,97 | 17303,70 | 44300,00 | 18327,34 | 43472,78 | 28985,74 |
| CAT | 43948,85 | 17938,30 | 90662,11 | 137600,42 | 9123,92 | 13491,42 | 8374,29 | 6746,52 |
| CCDC47 | 98395,98 | 223335,41 | 314415,75 | 363696,97 | 29875,66 | 220518,52 | 37130,98 | 82929,30 |
| CCT2 | 46705,27 | 42673,78 | 40106,69 | 41215,00 | 29124,88 | 29132,95 | 32698,25 | 29116,57 |
| CCT4 | 38569,10 | 50231,02 | 38909,50 | 25985,79 | 24034,78 | 27845,27 | 27570,77 | 24404,82 |
| CCT5 | 188364,14 | 192472,22 | 263385,81 | 161538,72 | 204904,69 | 220463,63 | 219519,33 | 198592,13 |
| CCT8 | 21736,65 | 36532,71 | 19268,88 | 14557,32 | 34100,48 | 31984,26 | 39993,15 | 23334,64 |
| CD109 | 66194,95 | 50917,79 | 329478,56 | 71712,05 | 113279,11 | 96232,84 | 100887,94 | 73790,51 |
| CD248 | 20963,45 | 16774,25 | 62706,45 | 32071,71 | 78819,77 | 34447,41 | 144641,80 | 55340,82 |
| CD44 | 89674,76 | 49144,51 | 547172,56 | 42798,00 | 27993,18 | 45772,50 | 63466,15 | 9820,17 |
| CD59 | 41139,88 | 64615,59 | 135329,20 | 30171,29 | 2674,93 | 6126,45 | 4051,76 | 3554,47 |
| CD81 | 1280355,25 | 1268773,13 | 2866425,00 | 563304,69 | 14498,07 | 27377,63 | 16486,23 | 17055,03 |
| CD82 | 827687,75 | 686888,94 | 1245914,75 | 288717,00 | 12566,49 | 10936,79 | 4657,29 | 19047,70 |
| CD9 | 225275,16 | 144800,31 | 416171,19 | 68152,88 | 5607,64 | 5076,84 | 4992,26 | 2819,10 |
| CD99 | 58936,92 | 41330,28 | 142814,98 | 12425,84 | 7220,84 | 4502,41 | 4227,41 | 2695,55 |
| CDC42 | 17198,74 | 25250,71 | 42076,22 | 13209,15 | 6658,03 | 9043,06 | 8643,26 | 7093,07 |
| CEMIP | 66286,58 | 37323,29 | 51075,29 | 31499,89 | 20366,97 | 30349,80 | 20847,42 | 19561,28 |
| CFAP100 | 38875,37 | 445070,03 | 785934,88 | 1187317,63 | 803276,88 | 2079396,50 | 500514,44 | 414348,06 |
| CFB | 84186,34 | 535687,13 | 685193,63 | 1363399,88 | 11724,26 | 19318,26 | 11145,28 | 15048,31 |
| CFL1 | 142875,39 | 62559,58 | 330214,75 | 56550,39 | 33611,13 | 15325,85 | 53924,62 | 20164,65 |
| CLIC1 | 55449,84 | 84791,42 | 65655,36 | 54314,70 | 57518,38 | 68338,98 | 61366,55 | 37979,84 |
| CLIC4 | 32989,30 | 42131,51 | 52625,24 | 15598,60 | 43035,75 | 16609,31 | 53985,89 | 43639,36 |
| CLTC | 174101,03 | 68319,63 | 218531,38 | 104028,30 | 29568,29 | 34909,11 | 28645,12 | 31742,38 |
| CLU | 413635,25 | 395382,34 | 1114507,75 | 288821,66 | 573745,75 | 459629,78 | 527010,38 | 569109,19 |
| COL12A1 | 1094462,38 | 964789,31 | 948742,00 | 755230,44 | 314804,75 | 543582,63 | 391256,78 | 410281,31 |
| COL15A1 | 454091,47 | 431581,69 | 588319,13 | 247094,34 | 25087,62 | 22339,44 | 23680,45 | 26145,33 |
| COL18A1 | 131287,50 | 148876,11 | 153396,81 | 68367,52 | 7425,69 | 7458,78 | 6475,80 | 9151,23 |
| COL1A1 | 1570860,00 | 4860649,50 | 8336601,50 | 7344409,00 | 85802,27 | 806543,06 | 225965,25 | 267396,84 |
| COL1A2 | 2094262,88 | 5514503,00 | 10602997,00 | 7579273,50 | 478105,03 | 977359,06 | 855385,06 | 686465,88 |
| COL2A1 | 316514,84 | 992220,00 | 1575176,25 | 1330038,88 | 21477,02 | 224432,00 | 87818,33 | 78466,34 |
| COL3A1 | 704849,25 | 1375128,63 | 2178305,75 | 1928364,38 | 75266,02 | 238795,33 | 108530,25 | 160031,48 |
| COL4A1 | 63330,21 | 78471,52 | 68788,70 | 98006,02 | 18383,91 | 42096,56 | 44589,04 | 52172,61 |
| COL4A2 | 237466,52 | 181589,17 | 550479,50 | 104970,49 | 128929,30 | 145714,27 | 280975,41 | 197108,69 |
| COL5A1 | 142501,78 | 162475,80 | 214381,55 | 270161,47 | 36989,10 | 89228,76 | 52781,50 | 61069,50 |
| COL5A2 | 75375,58 | 76278,28 | 80892,70 | 180744,09 | 30213,39 | 43579,47 | 66449,00 | 53818,16 |
| COL6A1 | 1285580,75 | 1293903,50 | 1747473,25 | 1188580,50 | 224514,97 | 198698,95 | 253139,09 | 463955,66 |
| COL6A2 | 1669595,50 | 1735331,38 | 2436262,50 | 1771532,13 | 243307,66 | 270112,81 | 216294,44 | 425988,56 |
| COL6A3 | 1694321,38 | 1700654,13 | 2181224,50 | 1540616,75 | 261184,41 | 267911,75 | 333278,34 | 268975,44 |
| COL7A1 | 77388,77 | 56308,56 | 51303,91 | 33363,46 | 7031,42 | 3711,67 | 6421,79 | 6559,98 |
| COMP | 4813,10 | 12129,90 | 41961,83 | 48429,84 | 12029,91 | 57051,62 | 23471,96 | 29806,15 |
| COPA | 25361,85 | 16953,65 | 26780,23 | 20140,69 | 17560,38 | 17004,41 | 17222,70 | 15553,79 |
| COPB1 | 10458,61 | 5137,13 | 12485,74 | 5082,27 | 12190,74 | 10014,47 | 11521,73 | 7491,86 |
| CPA4 | 15364,50 | 12061,95 | 21723,04 | 50794,54 | 15582,45 | 19204,19 | 41609,34 | 24349,48 |
| CPS1 | 12008,39 | 7301,40 | 22691,46 | 4945,61 | 6094,49 | 33391,28 | 15037,33 | 354,91 |
| CS | 38880,07 | 34772,05 | 39743,92 | 16238,96 | 46380,14 | 29478,12 | 42740,98 | 26569,40 |
| CSE1L | 6525,83 | 18592,81 | 4535,23 | 41542,59 | 18666,83 | 36176,43 | 31830,60 | 15451,11 |
| CSPG4 | 32969,81 | 22211,89 | 80908,65 | 26418,38 | 38928,35 | 40737,22 | 53552,14 | 32919,46 |
| CSTF2 | 373993,41 | 1270796,25 | 1939383,88 | 1817040,00 | 31109,09 | 129620,78 | 34331,92 | 68722,63 |
| CTHRC1 | 124875,95 | 33999,51 | 45826,66 | 59669,66 | 72108,93 | 41924,60 | 102979,05 | 100984,76 |
| CTSB | 48657,32 | 50801,88 | 52858,13 | 74782,59 | 706886,81 | 253018,80 | 691535,63 | 464844,78 |
| CTSD | 131003,55 | 45581,77 | 172582,06 | 167502,31 | 67556,40 | 37359,67 | 84501,45 | 50231,37 |
| DARS | 14105,56 | 16262,64 | 23439,86 | 15413,62 | 12801,74 | 11264,28 | 10211,36 | 12473,31 |
| DCD | 2665199,00 | 1532498,25 | 4156831,00 | 1789930,38 | 4333,24 | 19529,79 | 8378,77 | 9963,13 |
| DCN | 3275972,25 | 3915195,75 | 4844601,00 | 9506612,00 | 5220813,00 | 5800789,00 | 6675430,50 | 3889585,00 |
| DDX39B | 21644,65 | 30270,10 | 4359,35 | 55779,08 | 41806,51 | 46658,73 | 82313,54 | 30433,38 |
| DPYSL2 | 30915,59 | 19327,49 | 19824,83 | 17751,47 | 47841,91 | 52216,01 | 33340,27 | 39249,57 |
| DPYSL3 | 25865,20 | 10854,22 | 27636,51 | 13434,86 | 45224,25 | 33571,56 | 29463,13 | 30722,71 |
| DSTN | 103060,15 | 52413,09 | 309093,59 | 222357,70 | 6801,04 | 10036,58 | 9340,85 | 8336,87 |
| DYNC1H1 | 23436,62 | 19553,19 | 36899,11 | 11001,67 | 5095,60 | 2012,13 | 5119,47 | 5580,03 |
| EDIL3 | 971079,00 | 1381724,88 | 3006195,00 | 1360499,75 | 17204,79 | 62448,78 | 40040,74 | 52929,73 |
| EEA1 | 66264,48 | 38025,14 | 5118,63 | 41355,65 | 2523,19 | 9,39 | 5591,50 | 8223,13 |
| EEF1A1 | 106635,42 | 107287,06 | 167839,94 | 52987,05 | 95366,70 | 65096,64 | 98567,15 | 68290,41 |
| EEF1G | 57035,46 | 62202,27 | 54325,66 | 39001,96 | 75240,77 | 68798,53 | 81550,91 | 57835,85 |
| EEF2 | 171420,97 | 206040,77 | 250462,72 | 127614,92 | 336791,44 | 313304,34 | 318865,69 | 302743,28 |
| EFEMP1 | 39531,97 | 64601,81 | 113342,81 | 79662,96 | 575533,13 | 340693,44 | 749467,75 | 631943,38 |
| EFEMP2 | 12547,29 | 32694,80 | 64431,28 | 20301,38 | 148304,50 | 170797,50 | 112541,91 | 173032,25 |
| EFTUD2 | 128400,44 | 147350,20 | 113751,36 | 138184,78 | 254314,61 | 243484,86 | 218760,30 | 209963,83 |
| EHD2 | 94948,50 | 124370,74 | 168017,28 | 22261,29 | 15192,18 | 8364,64 | 12394,53 | 9203,51 |
| EHD3 | 46108,46 | 31810,27 | 167470,59 | 12832,79 | 9535,95 | 10215,17 | 9152,67 | 8372,72 |
| EIF3B | 15499,09 | 7912,16 | 10221,45 | 4345,15 | 21376,95 | 22029,11 | 23010,72 | 18744,88 |
| EIF4A2 | 47316,85 | 48271,30 | 45253,91 | 22997,73 | 32965,83 | 24709,84 | 40272,36 | 40058,87 |
| EIF4A3 | 10907,35 | 13298,34 | 12493,38 | 9181,42 | 4009,77 | 13444,54 | 7153,07 | 9726,15 |
| EIF5A | 18195,50 | 10238,87 | 21382,14 | 41347,07 | 5481,71 | 4216,47 | 3748,12 | 2215,49 |
| EMILIN1 | 794475,50 | 736932,31 | 600789,44 | 481149,16 | 143117,23 | 104617,51 | 78556,27 | 153586,03 |
| ENO1 | 89368,66 | 93737,95 | 152249,56 | 99861,70 | 547509,56 | 190141,64 | 691059,81 | 239164,36 |
| ENO3 | 32037,56 | 45093,79 | 87331,39 | 49687,10 | 262915,56 | 108692,95 | 341921,47 | 114913,54 |
| EPHB2 | 8001,02 | 4670,19 | 14622,24 | 2607,25 | 1475,95 | 1441,70 | 3291,98 | 2816,94 |
| EPRS | 12154,36 | 11655,13 | 7505,80 | 12470,96 | 7407,37 | 4888,96 | 4893,24 | 5580,36 |
| EXT2 | 4546947,00 | 52277,36 | 84844,10 | 68663,61 | 77487,86 | 220934,27 | 105120,38 | 97157,39 |
| EZR | 28472,17 | 16127,53 | 12370,75 | 6049,37 | 22587,21 | 12024,52 | 35993,79 | 11850,85 |
| F2 | 44175,21 | 61339,96 | 284570,16 | 72238,96 | 11237,01 | 54395,91 | 17241,60 | 47673,21 |
| FABP5 | 190044,11 | 180692,05 | 764186,50 | 2325147,00 | 12696,00 | 11356,96 | 11527,36 | 5684,47 |
| FARP1 | 28879,69 | 29404,96 | 10917,52 | 6209,83 | 2858,35 | 1925,24 | 2797,84 | 4505,18 |
| FASN | 34356,81 | 17811,76 | 117942,73 | 140475,72 | 5906,16 | 5260,79 | 7477,05 | 5124,95 |
| FAT1 | 35419,70 | 37828,08 | 59941,52 | 38872,27 | 13979,01 | 34035,21 | 20167,20 | 19760,70 |
| FBLN1 | 144740,58 | 159505,38 | 320387,78 | 169053,89 | 471998,28 | 364096,28 | 297924,94 | 573284,19 |
| FBLN2 | 14202,24 | 9814,47 | 7480,14 | 2633,20 | 11805,63 | 15484,12 | 11291,14 | 18218,00 |
| FBN2 | 10140,93 | 5866,15 | 2536,76 | 5198,83 | 35123,19 | 31117,79 | 59345,73 | 27698,88 |
| FERMT2 | 13305,84 | 5366,81 | 35732,68 | 8974,69 | 21074,69 | 19147,57 | 21581,45 | 19589,54 |
| FGA | 7797,06 | 18079,55 | 4242,24 | 34649,53 | 4882,00 | 31818,73 | 5602,13 | 11877,73 |
| FGB | 38051,73 | 43183,68 | 83228,72 | 75381,37 | 12763,12 | 45858,12 | 27922,10 | 29213,32 |
| FGG | 13484,51 | 21268,66 | 69665,17 | 37445,46 | 5963,23 | 15677,31 | 10876,85 | 12532,39 |
| FLNA | 226313,20 | 157816,61 | 191115,42 | 181497,69 | 473592,00 | 328375,06 | 487501,63 | 374479,22 |
| FLNB | 47996,84 | 36564,25 | 44261,59 | 39399,29 | 90218,10 | 63166,30 | 89211,66 | 73202,02 |
| FLNC | 195031,44 | 108754,93 | 133353,22 | 110793,86 | 326809,16 | 167401,95 | 359026,16 | 228704,81 |
| FN1 | 607109,63 | 1155777,38 | 934549,63 | 1282576,75 | 925198,50 | 2424881,50 | 738819,31 | 1767742,50 |
| FSCN1 | 57917,05 | 44386,89 | 93247,33 | 38576,43 | 132077,16 | 87238,77 | 161765,11 | 95362,52 |
| G6PD | 58875,02 | 34785,24 | 54999,32 | 22358,62 | 49364,61 | 35125,59 | 47086,77 | 30036,01 |
| GALNT2 | 14362,63 | 40816,86 | 78006,48 | 14999,42 | 17794,44 | 6944,83 | 44523,59 | 13961,21 |
| GANAB | 41295,88 | 27968,90 | 23676,38 | 15090,02 | 75085,96 | 58121,84 | 136109,23 | 50894,37 |
| GAPDH | 374496,16 | 516997,97 | 612935,88 | 556657,50 | 259610,44 | 295892,22 | 273372,00 | 251428,31 |
| GAS6 | 36891,55 | 36466,63 | 50933,24 | 37881,24 | 241961,22 | 322304,34 | 335080,22 | 250091,53 |
| GC | 10793,40 | 23549,86 | 16958,73 | 51533,71 | 57802,95 | 208440,41 | 87473,41 | 242726,59 |
| GDI2 | 41349,77 | 33940,79 | 66400,44 | 35013,04 | 53168,11 | 29739,69 | 89045,34 | 36283,05 |
| GLG1 | 15064,88 | 13561,43 | 11325,44 | 6401,63 | 1690,13 | 1965,81 | 2846,48 | 2027,77 |
| GNAI2 | 102068,83 | 138231,89 | 193766,06 | 40459,32 | 2091,60 | 30643,59 | 18282,58 | 7195,21 |
| GNB1 | 65815,13 | 48350,04 | 175601,00 | 31391,51 | 9656,18 | 7725,41 | 7892,00 | 4430,69 |
| GNB2 | 51119,62 | 79072,42 | 131545,53 | 26897,04 | 23952,93 | 12942,75 | 17048,25 | 16251,05 |
| GPC1 | 470172,25 | 533266,31 | 2073172,63 | 699521,50 | 106493,05 | 325765,06 | 213738,47 | 210489,48 |
| GPC6 | 66724,05 | 59024,51 | 114906,33 | 68502,39 | 5259,21 | 13173,84 | 8453,75 | 10162,20 |
| GPI | 11289,72 | 6422,06 | 22077,27 | 6219,31 | 76715,84 | 41793,63 | 80338,23 | 43144,75 |
| GREM1 | 203492,17 | 27041,04 | 53146,47 | 49888,80 | 106178,07 | 92079,24 | 138895,61 | 297624,50 |
| GSN | 27741,79 | 34478,99 | 80940,99 | 96399,97 | 133764,75 | 211526,84 | 172798,50 | 132986,58 |
| GSTA5 | 70240,60 | 105119,39 | 133921,94 | 160999,45 | 28458,53 | 142559,61 | 44173,10 | 36668,82 |
| GSTM5 | 16331,17 | 25225,97 | 28363,49 | 11983,25 | 28893,43 | 42800,55 | 35957,55 | 15240,47 |
| H4C1 | 274558,38 | 94150,71 | 211951,75 | 144403,19 | 27082,46 | 47644,54 | 17714,93 | 11905,47 |
| HBA1 | 309767,31 | 445924,56 | 705340,50 | 532828,63 | 145669,03 | 720985,00 | 192127,13 | 344245,44 |
| HBB | 351531,56 | 793248,00 | 618060,50 | 1143160,00 | 612507,13 | 3453485,00 | 854198,19 | 1438930,63 |
| HGF | 118039,80 | 24640,37 | 37244,22 | 21814,23 | 33276,82 | 26219,20 | 36817,22 | 38128,84 |
| HIST1H2BJ;HIST1H2BO; | 240870,17 | 85990,39 | 167001,73 | 108849,90 | 40542,61 | 86423,49 | 37196,02 | 18122,38 |
| HIST2H2AC;HIST2H2AA3 | 742614,56 | 236818,19 | 626127,19 | 297475,00 | 167298,86 | 224675,11 | 84691,59 | 78206,22 |
| HIST2H3A | 896228,38 | 265656,66 | 803099,38 | 344520,06 | 61577,84 | 95268,52 | 36084,99 | 36831,45 |
| HLA-A | 57714,41 | 63350,04 | 150824,02 | 28090,03 | 19989,62 | 17394,87 | 27675,32 | 10237,05 |
| HNRNPA1 | 17341,85 | 5032,77 | 4919,98 | 5375,94 | 5760,75 | 9443,65 | 7002,67 | 5855,62 |
| HNRNPM | 372405,50 | 246069,66 | 213085,89 | 13246,61 | 3575,49 | 2940,18 | 6473,31 | 545,80 |
| HNRNPU | 12001,93 | 17703,72 | 15131,17 | 7772,93 | 3717,38 | 7136,20 | 2307,35 | 1897,95 |
| HP | 44492,77 | 28413,10 | 676827,38 | 26118,45 | 16943,29 | 56905,26 | 16882,25 | 2648,61 |
| HSP90AA1 | 75413,80 | 95237,72 | 139863,27 | 89736,17 | 408294,78 | 414261,53 | 434175,72 | 354294,81 |
| HSP90AB1 | 282554,72 | 333189,88 | 193137,41 | 224441,44 | 1078226,25 | 903743,25 | 1224935,75 | 884800,50 |
| HSP90B1 | 26101,55 | 16001,83 | 14006,12 | 17340,32 | 175641,05 | 145605,50 | 206732,92 | 112481,64 |
| HSPA1A;HSPA1B | 276113,97 | 298560,03 | 357226,69 | 177962,98 | 79238,76 | 79925,55 | 76953,98 | 71123,74 |
| HSPA5 | 41190,00 | 33469,93 | 112108,14 | 65117,74 | 93437,39 | 59967,19 | 123681,25 | 53651,13 |
| HSPA8 | 240368,86 | 187865,14 | 187256,83 | 121549,74 | 104060,71 | 73997,77 | 116648,95 | 106722,80 |
| HSPB1 | 54097,34 | 35461,80 | 76839,01 | 41706,59 | 34527,54 | 25662,72 | 36797,93 | 16151,90 |
| HSPD1 | 15189,58 | 10429,34 | 26607,13 | 20056,04 | 63699,20 | 64809,27 | 59482,70 | 35188,29 |
| HSPG2 | 435656,44 | 549823,75 | 619976,38 | 235107,97 | 17665,63 | 34888,23 | 7988,06 | 32253,07 |
| HTRA1 | 67344,40 | 60335,58 | 83145,50 | 46925,20 | 91494,38 | 55710,86 | 103447,66 | 68608,32 |
| IDH1 | 28036,69 | 27768,02 | 40137,13 | 28533,84 | 77612,30 | 70018,02 | 79189,41 | 49136,18 |
| IGF2R | 16837,14 | 21149,54 | 26147,71 | 19355,02 | 20010,76 | 34410,00 | 21297,83 | 28208,07 |
| IGFBP3 | 9471,64 | 8791,69 | 22067,91 | 11380,77 | 23091,15 | 50137,56 | 169027,89 | 56203,80 |
| IGHG1 | 74876,32 | 10539,87 | 61978,59 | 71356,81 | 10101,85 | 6642,72 | 11468,76 | 50134,46 |
| IGSF8 | 68294,73 | 89209,66 | 113284,43 | 57807,66 | 1278,30 | 2197,63 | 1277,85 | 1837,03 |
| IQGAP1 | 174386,64 | 35255,82 | 1171902,63 | 251366,13 | 42551,19 | 24771,72 | 34320,50 | 41954,66 |
| ISYNA1 | 90824,79 | 72087,78 | 168159,70 | 105478,94 | 2888,10 | 6794,68 | 3008,73 | 5918,74 |
| ITGA1 | 35273,92 | 53461,39 | 96317,38 | 14338,83 | 3493,63 | 1261,33 | 1775,20 | 1738,32 |
| ITGA3 | 127346,20 | 149743,02 | 263198,00 | 76339,69 | 1211,88 | 5502,58 | 8439,97 | 2023,04 |
| ITGA5 | 34280,68 | 40681,94 | 31953,23 | 9510,95 | 2881,01 | 3109,14 | 6784,83 | 4472,86 |
| ITGAV | 206155,22 | 229916,34 | 531746,31 | 108137,57 | 7257,09 | 10474,05 | 2148,24 | 10780,90 |
| ITGB1 | 115057,80 | 136066,75 | 324838,75 | 74425,63 | 3638,34 | 5930,63 | 4456,92 | 4667,30 |
| ITIH1 | 157387,91 | 310258,13 | 446505,63 | 508333,63 | 43507,91 | 146274,88 | 43479,74 | 95916,55 |
| ITIH2 | 5569703,00 | 14570614,00 | 21079388,00 | 12028043,00 | 77944,19 | 654234,75 | 124162,61 | 362247,66 |
| ITIH3 | 202069,66 | 541723,00 | 336690,50 | 627877,63 | 59418,63 | 241345,19 | 102217,79 | 216079,03 |
| ITIH4 | 20030,20 | 46973,05 | 13898,42 | 101998,23 | 104248,67 | 444019,28 | 197559,41 | 380951,22 |
| JUP | 58325,77 | 37274,51 | 132950,36 | 170432,83 | 578,42 | 1518,34 | 3620,38 | 715,37 |
| KPNB1 | 23366,42 | 16683,52 | 19220,04 | 13238,14 | 68650,11 | 68847,00 | 53191,64 | 60730,56 |
| KPRP | 31186,47 | 45663,43 | 71798,51 | 263468,56 | 6423,21 | 9753,62 | 690,49 | 1090,68 |
| KRT1 | 14072378,00 | 5294913,00 | 24573272,00 | 37199816,00 | 36866,58 | 114267,36 | 51848,38 | 145400,66 |
| KRT10 | 18194604,00 | 7014236,00 | 28457528,00 | 14374153,00 | 28523,91 | 88342,56 | 47092,32 | 95053,45 |
| KRT14 | 11326415,00 | 3810483,75 | 17642314,00 | 16968082,00 | 49301,19 | 102625,51 | 43829,61 | 133977,77 |
| KRT16 | 569814,25 | 142613,83 | 891831,31 | 708446,69 | 6884,69 | 4362,44 | 1970,97 | 14550,64 |
| KRT17 | 177475,36 | 41395,01 | 215070,50 | 407875,91 | 2989,57 | 6599,89 | 2228,84 | 2297,01 |
| KRT19 | 1383103,88 | 1083571,38 | 1314269,75 | 403385,81 | 12885,16 | 38972,48 | 25064,00 | 15103,51 |
| KRT2 | 13736436,00 | 4914892,00 | 21080910,00 | 11580109,00 | 32362,71 | 104694,73 | 53234,24 | 104384,52 |
| KRT5 | 2639798,75 | 1022018,50 | 3992585,00 | 4336944,00 | 9069,25 | 24274,07 | 8594,01 | 23134,44 |
| KRT6A | 3575214,00 | 1440562,50 | 4864597,50 | 5264088,50 | 16142,77 | 43582,34 | 19934,10 | 52895,11 |
| KRT6B | 15358256,00 | 4809123,50 | 21678234,00 | 43449604,00 | 36073,40 | 82161,65 | 47217,04 | 133343,98 |
| KRT77 | 62247,18 | 21934,23 | 76582,55 | 42621,77 | 2803,85 | 5180,20 | 1975,16 | 1124,29 |
| KRT9 | 12902477,00 | 5725966,00 | 27047976,00 | 73616528,00 | 33135,27 | 98953,10 | 41333,26 | 154350,31 |
| LAMA1 | 370265,06 | 338625,78 | 196348,59 | 152368,23 | 17058,69 | 24061,45 | 11483,91 | 23284,48 |
| LAMA2 | 21187,89 | 10766,98 | 17474,45 | 19336,67 | 1456,72 | 1971,48 | 1484,92 | 3607,01 |
| LAMA4 | 542989,38 | 372030,34 | 648137,25 | 252052,47 | 23936,91 | 26706,59 | 15806,63 | 28584,69 |
| LAMA5 | 357380,19 | 231533,86 | 213534,78 | 86913,75 | 11722,38 | 7256,01 | 6926,22 | 9947,18 |
| LAMB1 | 1117242,38 | 779169,50 | 1147238,50 | 523012,22 | 57977,38 | 55992,70 | 40663,92 | 69202,13 |
| LAMB2 | 146872,13 | 143474,42 | 143040,06 | 67850,22 | 14328,14 | 14747,54 | 15841,51 | 17994,53 |
| LAMC1 | 1264674,50 | 990923,56 | 1396385,00 | 636907,56 | 64691,68 | 73992,14 | 55600,71 | 80785,90 |
| LAMP1 | 90394,17 | 63330,54 | 141969,45 | 67056,19 | 4796,48 | 5538,25 | 4737,33 | 2889,13 |
| LAMP2 | 49418,76 | 54183,73 | 141054,02 | 44968,57 | 4872,69 | 5080,96 | 3830,52 | 3109,85 |
| LDHA | 184849,78 | 266769,50 | 292612,25 | 197269,06 | 2938097,25 | 2160623,00 | 2890375,75 | 1886078,75 |
| LDHB | 13168,83 | 31636,81 | 3003,80 | 11857,07 | 366898,53 | 284961,47 | 337232,03 | 158669,22 |
| LGALS1 | 80787,20 | 60364,69 | 66320,91 | 23081,63 | 29721,13 | 28027,49 | 29111,39 | 25204,16 |
| LGALS3BP | 272222,53 | 185262,58 | 308742,69 | 193237,48 | 109051,66 | 128846,90 | 124698,23 | 127432,94 |
| LMNA | 39551,98 | 22378,40 | 37876,37 | 50910,84 | 146070,34 | 101733,47 | 127040,41 | 98278,12 |
| LOXL2 | 39083,66 | 58258,60 | 83338,95 | 52976,43 | 144079,92 | 181992,72 | 314181,66 | 210996,78 |
| LRP1 | 44079,66 | 25465,79 | 60035,90 | 41819,54 | 26893,49 | 39953,17 | 24986,06 | 26690,09 |
| LRRC17 | 31865,26 | 47799,67 | 39566,90 | 23553,63 | 4020,98 | 3560,73 | 684,57 | 6128,55 |
| LTBP1 | 103785,88 | 90762,91 | 99485,02 | 97574,77 | 109337,94 | 106365,18 | 89132,23 | 143460,05 |
| LTBP2 | 68933,92 | 29045,45 | 34225,17 | 68077,11 | 131178,34 | 173842,72 | 188656,66 | 130242,78 |
| LTF | 170816,30 | 102058,45 | 650867,75 | 184080,31 | 367106,06 | 690837,50 | 1009312,00 | 1297936,63 |
| LUM | 15449,67 | 38442,45 | 41744,80 | 134395,25 | 198798,30 | 89947,58 | 391510,63 | 85688,39 |
| MAMDC2 | 337776,69 | 363723,94 | 849155,63 | 300578,41 | 7117,86 | 17880,94 | 9033,89 | 9391,24 |
| MAPK1 | 76366,88 | 19998,73 | 135233,33 | 74796,41 | 14414,90 | 17176,05 | 16859,99 | 3337,98 |
| MARCKS | 44185,22 | 43493,00 | 89347,34 | 24131,35 | 19504,97 | 7232,75 | 27833,21 | 15983,46 |
| MCAM | 5905,77 | 17697,53 | 25818,40 | 9516,76 | 3071,69 | 2466,27 | 3999,40 | 2095,23 |
| MDH2 | 47873,82 | 51771,48 | 60162,51 | 69590,09 | 57315,03 | 25278,73 | 77164,84 | 28158,08 |
| MFAP4 | 101623,45 | 23545,09 | 57789,72 | 17786,73 | 194427,47 | 72747,21 | 199956,03 | 62453,01 |
| MFGE8 | 821966,63 | 1240407,88 | 2395417,75 | 1386315,50 | 32499,33 | 56940,84 | 53546,68 | 53176,21 |
| MMP2 | 17009,52 | 28277,46 | 22153,37 | 62160,55 | 807829,69 | 1100465,38 | 568548,81 | 752849,94 |
| MSN | 157189,77 | 116638,48 | 402821,19 | 84511,65 | 307475,38 | 111030,57 | 472123,06 | 186977,44 |
| MTHFD1 | 142322,20 | 2976,32 | 81586,88 | 35257,43 | 3055,47 | 1977,56 | 2930,31 | 1627,27 |
| MYH9 | 175350,52 | 120423,05 | 108211,73 | 110034,99 | 102719,57 | 81316,86 | 63235,59 | 94813,20 |
| NIBAN2 | 18517,39 | 16362,75 | 10319,76 | 10047,92 | 21998,96 | 14489,70 | 21715,98 | 14623,03 |
| NID1 | 1387527,38 | 1186337,25 | 1903387,88 | 963453,31 | 118385,18 | 124298,10 | 123109,22 | 165447,56 |
| NID2 | 281577,22 | 183363,84 | 212897,09 | 128031,98 | 51768,36 | 30013,33 | 59488,29 | 30210,19 |
| NME2 | 8116,10 | 4068,72 | 5610,66 | 6558,52 | 130066,91 | 69438,49 | 150268,80 | 50834,11 |
| NOTCH3 | 3333,40 | 2240,62 | 1736,39 | 8468,63 | 7254,53 | 15638,55 | 12529,27 | 12511,01 |
| NQO1 | 42358,75 | 20693,96 | 40514,40 | 14306,06 | 29670,20 | 27036,09 | 21212,87 | 22302,40 |
| NRP1 | 17975,53 | 29194,49 | 56660,93 | 33838,43 | 24408,08 | 38013,63 | 51374,46 | 23211,81 |
| NT5E | 266782,56 | 450096,25 | 676312,38 | 196795,25 | 7053,12 | 7615,48 | 10536,62 | 12183,13 |
| NTN4 | 57180,57 | 37580,46 | 125385,26 | 34852,66 | 5041,63 | 18908,88 | 7551,40 | 5500,23 |
| NUMA1 | 97907,58 | 63060,59 | 35086,15 | 51709,61 | 7250,57 | 7491,08 | 5647,84 | 7737,14 |
| P4HB | 41047,25 | 27519,45 | 30494,57 | 33414,71 | 95419,83 | 48875,96 | 84387,68 | 66826,88 |
| PACSIN3 | 21281,44 | 9622,43 | 49356,72 | 12462,89 | 1605,18 | 3306,63 | 2739,80 | 3267,54 |
| PAICS | 64771,80 | 52226,25 | 4004,55 | 55308,75 | 25214,22 | 22653,81 | 22350,36 | 18095,02 |
| PAPPA | 155513,73 | 100610,66 | 136039,67 | 120052,85 | 32184,92 | 49527,19 | 49959,07 | 47711,87 |
| PARD3 | 89327,96 | 249032,30 | 323487,16 | 455470,38 | 127881,93 | 558347,00 | 156233,30 | 270752,25 |
| PCBP2 | 16855,57 | 27295,64 | 16541,95 | 13660,56 | 17468,11 | 10309,27 | 14678,93 | 15427,28 |
| PCOLCE | 28513,50 | 43993,95 | 48844,12 | 54754,97 | 370230,19 | 373916,41 | 358889,09 | 383170,19 |
| PDCD6IP | 115378,45 | 98798,85 | 130926,28 | 59166,65 | 9506,84 | 6942,70 | 9850,63 | 8515,44 |
| PDIA3 | 35720,59 | 19983,85 | 38553,43 | 28981,79 | 85372,54 | 32939,21 | 149561,70 | 47092,01 |
| PFN1 | 90810,33 | 83280,63 | 77826,56 | 77278,65 | 43310,69 | 41645,14 | 33480,33 | 51820,81 |
| PFN2 | 20404,11 | 18206,65 | 4155,02 | 13030,87 | 16754,26 | 12316,09 | 9802,45 | 15299,69 |
| PGD | 31585,70 | 21072,32 | 7271,27 | 5359,47 | 58293,11 | 41960,56 | 58015,15 | 43155,76 |
| PGK1 | 29463,84 | 44375,36 | 70705,68 | 31967,55 | 107185,34 | 43100,75 | 147789,13 | 91451,43 |
| PIP | 97780,02 | 87776,70 | 142581,67 | 44320,35 | 6729,83 | 5303,01 | 6382,30 | 431,13 |
| PKLR | 266388,34 | 290166,09 | 168659,53 | 218406,89 | 173548,73 | 197353,06 | 167697,42 | 89335,26 |
| PKM | 468003,06 | 494081,22 | 403203,91 | 305959,06 | 717120,44 | 542645,81 | 627482,50 | 597649,63 |
| PLAT | 23249,62 | 22627,65 | 5003,76 | 16007,67 | 101882,84 | 47972,75 | 32625,26 | 75924,04 |
| PLEC | 71283,62 | 33685,68 | 46533,51 | 53202,72 | 87981,93 | 51756,84 | 63574,48 | 64411,56 |
| PLG | 36024,04 | 64102,74 | 46797,34 | 109321,07 | 83585,38 | 322136,56 | 116309,84 | 173711,89 |
| PLOD1 | 298341,69 | 346142,53 | 442409,50 | 397643,63 | 760530,69 | 842842,31 | 759580,94 | 1006718,75 |
| PLOD3 | 57618,00 | 36273,18 | 29200,19 | 57416,62 | 145383,13 | 107484,07 | 95720,77 | 94554,85 |
| PLSCR3 | 237610,81 | 193373,84 | 497540,59 | 75555,54 | 2825,37 | 4636,30 | 2271,96 | 2727,30 |
| PLTP | 19593,54 | 63680,57 | 36301,43 | 31368,19 | 305186,38 | 225315,53 | 242124,92 | 335731,19 |
| POSTN | 87033,20 | 164086,66 | 82998,34 | 60549,18 | 264949,78 | 82823,65 | 85343,28 | 295430,56 |
| POTEJ | 60320,21 | 74396,73 | 77817,27 | 27095,80 | 121933,87 | 127373,05 | 125712,73 | 80447,38 |
| PPIA | 14039,76 | 28941,38 | 76815,88 | 16973,49 | 13020,00 | 22654,31 | 12516,41 | 12448,52 |
| PPP1CC | 41328,73 | 30866,31 | 1244,01 | 7099,72 | 33248,30 | 27381,68 | 33015,22 | 24112,08 |
| PPP2R1A | 13501,81 | 17215,74 | 8502,45 | 9754,21 | 31476,29 | 38535,75 | 39077,57 | 19640,41 |
| PRDX1 | 40804,61 | 35031,60 | 83309,88 | 105725,30 | 40016,56 | 28543,51 | 53958,87 | 31810,68 |
| PRDX2 | 26465,48 | 14664,92 | 70670,84 | 81655,16 | 13695,75 | 9340,88 | 16446,37 | 12958,65 |
| PRDX6 | 32392,54 | 42272,32 | 135715,61 | 23209,09 | 19500,75 | 20813,58 | 22831,75 | 24593,56 |
| PRKDC | 6870,47 | 72874,79 | 13223,70 | 19967,16 | 2269,49 | 15529,20 | 3766,38 | 6288,28 |
| PRSS1;PRSS2;PRSS3P2 | 1343874,00 | 11629502,00 | 117135192,00 | 554847,31 | 385418,59 | 539852,75 | 284730,13 | 35768,57 |
| PRSS23 | 11944,48 | 6269,03 | 9335,29 | 8565,09 | 3925,82 | 11610,38 | 7935,11 | 5935,96 |
| PSMA1 | 37250,16 | 22738,87 | 39646,40 | 48032,41 | 96572,84 | 122400,54 | 85073,04 | 82604,41 |
| PSMA3 | 65634,23 | 36784,79 | 80568,19 | 52196,56 | 111038,51 | 130764,32 | 99720,93 | 59833,30 |
| PSMA5 | 36009,65 | 32338,38 | 30811,76 | 31344,87 | 102552,57 | 118347,08 | 85122,95 | 82798,88 |
| PSMA7 | 21491,84 | 15235,60 | 726900,44 | 37778,34 | 63076,18 | 82993,62 | 52901,62 | 45119,36 |
| PSMB6 | 63658,42 | 44721,78 | 43230,85 | 50650,41 | 184248,16 | 240848,95 | 176424,41 | 123909,59 |
| PSMC2 | 27543,83 | 27275,91 | 18984,62 | 4295,71 | 9391,50 | 5170,85 | 7406,32 | 8017,96 |
| PSMC5 | 49720,79 | 11875,65 | 7662,11 | 20184,49 | 11047,13 | 12505,38 | 12010,40 | 6170,72 |
| PSMC6 | 16812,71 | 11549,22 | 93283,75 | 6217,77 | 6775,50 | 4894,26 | 5538,21 | 3872,33 |
| PSMD11 | 334212,63 | 130180,19 | 107028,45 | 137895,31 | 79498,60 | 55304,05 | 62715,71 | 37827,54 |
| PSMD13 | 10516,33 | 26135,61 | 26711,91 | 11108,46 | 12500,03 | 8792,98 | 9593,58 | 7376,64 |
| PSMD2 | 9582,79 | 17208,22 | 12772,09 | 5065,95 | 17688,35 | 10585,88 | 14100,92 | 10428,80 |
| PTK7 | 57114,18 | 46496,68 | 69988,98 | 35025,08 | 28872,37 | 24931,23 | 48335,23 | 74479,43 |
| PTX3 | 12634336,00 | 19734224,00 | 49123000,00 | 18489482,00 | 397805,91 | 413449,34 | 522247,22 | 549026,50 |
| PXDN | 28780,65 | 51834,27 | 49102,76 | 27695,08 | 140984,48 | 190635,28 | 161721,48 | 177483,06 |
| PYGL | 20373,21 | 23495,79 | 415355,81 | 8208,34 | 22623,54 | 27205,16 | 22799,80 | 22019,04 |
| QSOX1 | 426490,38 | 626534,88 | 818537,19 | 628714,56 | 7134007,00 | 4165606,00 | 9298124,00 | 6473022,50 |
| RAB10 | 20504,31 | 17842,95 | 23440,24 | 13067,73 | 5371,28 | 4466,34 | 4683,04 | 5496,18 |
| RAB11A;RAB11B | 20968,27 | 42110,28 | 31294,85 | 34433,79 | 10678,77 | 11988,30 | 8988,55 | 10990,64 |
| RAB14 | 30318,03 | 20698,00 | 46203,58 | 14208,55 | 3666,27 | 7822,11 | 5348,07 | 8156,76 |
| RAB21 | 16164,96 | 8439,54 | 32692,93 | 6079,58 | 2384,35 | 1867,96 | 2700,57 | 3113,56 |
| RAB5B | 50141,57 | 32048,38 | 14987,02 | 30567,71 | 6614,48 | 9063,73 | 5564,00 | 8731,19 |
| RAB5C | 150574,63 | 69385,00 | 190521,97 | 36521,39 | 18686,85 | 14633,35 | 15317,36 | 9173,31 |
| RAB7A | 81507,51 | 77559,66 | 134206,09 | 106715,51 | 2382,30 | 6843,34 | 3360,38 | 2570,03 |
| RAC1 | 87195,79 | 137075,45 | 218380,33 | 84536,16 | 28057,27 | 26269,18 | 38182,48 | 17481,86 |
| RACK1 | 126097,73 | 94918,98 | 132384,39 | 37617,83 | 55259,96 | 64751,97 | 56454,67 | 46772,18 |
| RAN | 52664,29 | 61498,49 | 48543,63 | 38979,41 | 32674,19 | 43133,78 | 24464,16 | 33891,39 |
| RANGAP1 | 32023,27 | 7931,65 | 32353,03 | 8483,43 | 27311,58 | 22903,47 | 26274,38 | 25730,92 |
| RAP1A | 69394,77 | 105306,41 | 215230,64 | 64249,81 | 5733,04 | 16294,99 | 3973,61 | 9767,96 |
| RDX | 83185,49 | 64104,16 | 154404,30 | 39542,50 | 164735,03 | 68314,02 | 251715,52 | 117902,12 |
| RECK | 7765,35 | 14133,28 | 29593,88 | 5071,71 | 1194,64 | 5440,17 | 4077,77 | 4362,20 |
| RELN | 37692,08 | 27453,55 | 45155,92 | 20909,60 | 4576,36 | 26992,77 | 11021,01 | 5712,03 |
| RHOC | 33957,86 | 20360,57 | 88402,37 | 19344,29 | 16199,86 | 16705,28 | 15339,10 | 8253,24 |
| RNH1 | 7614,31 | 5249,56 | 5807,45 | 2494,13 | 46494,40 | 29202,92 | 54794,02 | 35016,08 |
| ROCK1 | 46077,84 | 93256,14 | 238076,78 | 368649,16 | 88606,46 | 183256,48 | 105585,12 | 102692,84 |
| RPL11 | 22362,59 | 27161,12 | 7006,12 | 11786,76 | 9187,23 | 15813,64 | 10978,94 | 4098,42 |
| RPL15 | 17385,12 | 15380,84 | 14571,94 | 18293,87 | 5658,45 | 11700,67 | 3624,92 | 3250,92 |
| RPL3 | 38511,56 | 21177,88 | 83541,84 | 10953,62 | 2912,76 | 28987,03 | 7553,79 | 11730,05 |
| RPL35A | 39472,66 | 35414,24 | 48889,06 | 27919,62 | 9949,84 | 14111,20 | 12815,46 | 5557,11 |
| RPL6 | 2347,43 | 9969,22 | 7129,18 | 10103,35 | 6825,34 | 3181,90 | 4808,56 | 3298,01 |
| RPL8 | 29278,95 | 31382,90 | 27059,13 | 35353,36 | 15812,97 | 19082,61 | 19951,22 | 1697,59 |
| RPLP0 | 7076,39 | 13443,10 | 41418,80 | 5481,46 | 43868,46 | 28593,51 | 39330,40 | 28022,74 |
| RPS11 | 16871,25 | 18068,63 | 39255,74 | 8927,64 | 6010,73 | 7665,30 | 9290,21 | 6359,90 |
| RPS14 | 16993,30 | 16411,09 | 20131,16 | 19861,79 | 4828,60 | 8801,59 | 5896,57 | 2609,09 |
| RPS18 | 26934,40 | 34615,68 | 44440,17 | 18886,65 | 11915,48 | 11835,56 | 16278,80 | 9222,13 |
| RPS2 | 37237,39 | 47009,93 | 13010,33 | 35698,88 | 12450,44 | 14397,25 | 12174,10 | 10995,41 |
| RPS27L | 21189,12 | 19052,94 | 15596,98 | 34251,28 | 2561,95 | 24749,34 | 17698,18 | 8711,20 |
| RPS3 | 51711,57 | 55866,70 | 72499,48 | 33741,71 | 24752,37 | 15789,63 | 30548,25 | 27372,73 |
| RPS4X | 81134,40 | 41357,52 | 44247,31 | 18704,54 | 8937,63 | 13629,83 | 12127,46 | 6647,40 |
| RPS8 | 43693,07 | 18936,69 | 38209,61 | 27858,90 | 5272,96 | 27191,69 | 8396,38 | 2974,22 |
| RPSA | 74433,83 | 91508,77 | 32513,90 | 50670,80 | 52825,20 | 41029,59 | 40312,58 | 40793,40 |
| RRBP1 | 4267756,50 | 4535982,00 | 1547685,75 | 1735881,00 | 211886,69 | 186566,36 | 137041,77 | 233609,27 |
| RTN4 | 487426,69 | 321520,41 | 588247,56 | 269235,69 | 20199,13 | 39983,75 | 20830,68 | 18030,40 |
| S100A11 | 45474,23 | 23905,99 | 61085,44 | 36219,32 | 12013,34 | 21187,22 | 23655,67 | 6943,81 |
| S100A6 | 57913,87 | 54421,91 | 136000,88 | 35560,34 | 168603,41 | 80979,11 | 108174,48 | 79616,11 |
| S100A8 | 80569,79 | 55815,52 | 116939,44 | 115573,95 | 1028,07 | 6450,50 | 1863,25 | 161,13 |
| SDC1 | 2265998,50 | 2319404,25 | 698671,56 | 1585009,88 | 49375,34 | 67533,48 | 62836,07 | 31261,56 |
| SDC4 | 495175,66 | 369045,94 | 731425,50 | 540720,81 | 39563,97 | 91762,13 | 34222,30 | 70946,20 |
| SDCBP | 693067,75 | 674077,31 | 742682,56 | 366069,44 | 11347,65 | 14546,67 | 7891,48 | 14311,44 |
| SEC23A | 18616,73 | 8832,53 | 15190,25 | 21890,14 | 33194,95 | 45026,70 | 28876,89 | 23751,19 |
| SEPTIN2 | 33923,79 | 27722,63 | 18890,41 | 53755,80 | 22068,03 | 16887,26 | 19834,54 | 19372,07 |
| SEPTIN7 | 24906,47 | 18861,40 | 277283,34 | 24463,13 | 16499,36 | 15247,01 | 17086,30 | 15140,88 |
| SERPINA1 | 393788,81 | 3223,47 | 39824,10 | 29113,11 | 36848,33 | 135966,80 | 61838,29 | 34066,59 |
| SERPINC1 | 68612,09 | 110493,88 | 180121,44 | 227980,72 | 193707,55 | 561568,31 | 396874,59 | 506453,75 |
| SERPINE1 | 26587,92 | 37859,31 | 33924,49 | 31937,15 | 261020,83 | 389733,22 | 2149854,75 | 427063,63 |
| SERPINE2 | 8838,97 | 14979,04 | 35299,67 | 11009,70 | 11998,27 | 25320,93 | 51788,57 | 23116,04 |
| SERPINF1 | 55831,05 | 143280,95 | 164595,56 | 389200,28 | 161764,58 | 124599,30 | 151403,80 | 130510,25 |
| SERPINF2 | 51747,04 | 55145,80 | 26514,95 | 135038,88 | 202019,11 | 758161,31 | 350057,81 | 536864,31 |
| SLC16A3 | 219132,98 | 223352,84 | 509987,56 | 105973,27 | 1822,63 | 2434,59 | 971,17 | 1479,12 |
| SLC1A5 | 422805,16 | 319358,22 | 758663,75 | 66883,86 | 719,10 | 1567,63 | 1165,23 | 627,54 |
| SLC25A10 | 878874,88 | 302814,31 | 72202,68 | 114074,66 | 3032,60 | 15038,21 | 22865,50 | 2588,84 |
| SLC25A5 | 22300,37 | 14878,50 | 57705,39 | 23469,87 | 2296,30 | 13911,10 | 5712,09 | 451,83 |
| SLC3A2 | 140928,42 | 155118,48 | 254984,94 | 59662,41 | 5992,68 | 13203,22 | 10677,87 | 7841,13 |
| SND1 | 39520,07 | 30612,12 | 48697,10 | 13815,77 | 48730,62 | 43175,51 | 46382,29 | 24537,71 |
| SPTAN1 | 41665,38 | 14571,68 | 12754,64 | 23858,02 | 29472,11 | 21103,91 | 29816,16 | 30500,08 |
| SPTBN1 | 26633,01 | 14485,56 | 32634,48 | 16454,89 | 20546,00 | 16609,79 | 18063,56 | 22076,96 |
| SRP14 | 43934,38 | 83171,43 | 276141,09 | 67377,61 | 2120,02 | 2402,24 | 2652,31 | 1689,56 |
| SSC5D | 317717,66 | 418495,53 | 370738,38 | 419256,72 | 10269,76 | 11300,50 | 13002,51 | 14398,54 |
| STAM | 82926,94 | 214568,63 | 72632,98 | 866343,38 | 169260,69 | 814864,69 | 397369,31 | 104238,98 |
| STC1 | 49746,04 | 23820,32 | 16619,63 | 48548,97 | 10927,26 | 12312,70 | 5549,44 | 18503,70 |
| STC2 | 36203,50 | 39670,84 | 19662,50 | 45205,89 | 28526,13 | 28560,41 | 73395,88 | 25996,01 |
| STK17A | 1251375,50 | 397344,09 | 623721,44 | 293396,00 | 118607,41 | 166205,91 | 198035,08 | 109091,43 |
| SVEP1 | 46593,63 | 39463,57 | 24154,80 | 34005,99 | 17177,01 | 12801,50 | 10912,01 | 16202,57 |
| TAGLN | 14101,15 | 21850,80 | 42526,25 | 12756,16 | 6281,04 | 4784,08 | 7498,33 | 22913,89 |
| TAGLN2 | 14424,57 | 29946,94 | 41627,01 | 22542,29 | 7776,24 | 18968,24 | 5942,03 | 17722,25 |
| TCP1 | 16425,54 | 17952,58 | 44192,32 | 12970,17 | 21665,56 | 23406,73 | 28369,39 | 21757,98 |
| TF | 87673,30 | 444905,16 | 109598,60 | 133612,36 | 162321,84 | 679748,13 | 300348,28 | 112462,61 |
| TFRC | 20353,44 | 33188,78 | 54093,82 | 1446,96 | 10369,20 | 20829,56 | 10503,01 | 10681,32 |
| TGFBI | 415058,13 | 408472,00 | 525681,00 | 407773,72 | 953303,81 | 542699,75 | 564374,63 | 868893,00 |
| THBS1 | 4441583,50 | 3224823,00 | 4177554,00 | 3188926,00 | 1027930,44 | 2891196,50 | 1222502,50 | 1161346,13 |
| THBS2 | 210504,72 | 296401,50 | 330225,84 | 255075,45 | 101784,16 | 402082,91 | 97896,50 | 226970,91 |
| TLN1 | 31441,52 | 45542,36 | 70226,35 | 45397,48 | 61507,84 | 66960,99 | 64922,38 | 61817,62 |
| TNC | 199242,63 | 349332,94 | 222012,31 | 214566,28 | 42491,52 | 34810,35 | 25921,51 | 53052,11 |
| TNFAIP6 | 38140,31 | 49328,31 | 67932,87 | 27124,92 | 8168,91 | 16429,58 | 9381,93 | 6594,65 |
| TNPO1 | 6485,62 | 9200,95 | 13435,39 | 1861,92 | 61864,41 | 60250,91 | 55751,69 | 50812,13 |
| TPI1 | 29203,94 | 26902,55 | 47980,05 | 38995,19 | 293405,59 | 72682,47 | 341464,88 | 224187,39 |
| TRIM28 | 25939,46 | 48187,96 | 178298,69 | 70554,07 | 18292,86 | 7866,03 | 14431,94 | 14936,41 |
| TSPAN4 | 460705,03 | 417739,00 | 1075294,38 | 235233,80 | 4209,01 | 5339,08 | 7213,52 | 5070,16 |
| TSPAN6 | 172515,45 | 139644,70 | 291528,91 | 98872,73 | 2569,64 | 4584,54 | 2691,49 | 3273,10 |
| TTYH3 | 112060,20 | 120213,86 | 203120,61 | 74929,80 | 454,65 | 2902,08 | 1865,24 | 3456,89 |
| TUBA1A | 101279,76 | 306187,72 | 76494,80 | 27831,31 | 177058,73 | 142971,55 | 113542,50 | 152813,67 |
| TUBA1B | 529399,25 | 498238,09 | 638122,69 | 299282,47 | 383722,34 | 282507,69 | 281855,69 | 356083,38 |
| TUBA1C | 16336,95 | 16727,94 | 28091,99 | 10961,79 | 4348,72 | 2456,43 | 3009,89 | 4295,00 |
| TUBB | 304968,63 | 166048,66 | 359765,34 | 103777,95 | 173040,28 | 102049,98 | 137077,73 | 115758,26 |
| TUBB3 | 11618,12 | 3882,40 | 25025,60 | 5194,71 | 13061,49 | 9728,31 | 6470,81 | 5596,87 |
| TUBB4B | 387484,97 | 289096,00 | 375068,31 | 207256,05 | 228519,98 | 147561,19 | 172649,45 | 226400,02 |
| TUBB6 | 144637,80 | 103589,29 | 334996,84 | 290739,91 | 51704,09 | 51317,88 | 41744,60 | 38122,19 |
| TXN | 44866,99 | 34529,51 | 48189,67 | 105419,21 | 10086,47 | 7335,28 | 6814,99 | 839,28 |
| TXNRD1 | 27649,68 | 12709,73 | 15210,90 | 14977,71 | 152913,94 | 100657,41 | 131067,69 | 51794,61 |
| UBA1 | 36022,02 | 46150,09 | 99065,54 | 18983,21 | 93426,36 | 88629,96 | 81405,60 | 63590,60 |
| UBB;UBC;RPS27A;UBA52 | 2857177,75 | 2545014,50 | 4385605,50 | 1943774,25 | 115097,72 | 147110,91 | 112783,48 | 122634,48 |
| UGDH | 66400,88 | 27425,67 | 369078,00 | 13989,78 | 19203,67 | 11134,25 | 18771,24 | 15027,78 |
| USP5 | 13367,50 | 8102,78 | 11722,97 | 7489,21 | 25670,29 | 25924,92 | 23619,90 | 18589,55 |
| VCAN | 1430074,75 | 2836173,00 | 3856858,25 | 1692010,63 | 5293,17 | 8871,71 | 4927,82 | 7880,72 |
| VCL | 24925,88 | 19412,60 | 40039,95 | 27709,86 | 177665,94 | 159384,98 | 172711,78 | 126299,43 |
| VCP | 8902,49 | 24298,20 | 25976,24 | 35663,01 | 37820,55 | 85661,39 | 65491,42 | 50918,41 |
| VIM | 34627,30 | 27473,02 | 39206,67 | 30903,25 | 188024,83 | 71654,20 | 127254,21 | 80925,25 |
| VTN | 127914,48 | 404835,91 | 209954,34 | 594266,88 | 69814,57 | 344741,16 | 100557,43 | 111408,28 |
| VWA1 | 4864,11 | 5025,81 | 7595,47 | 4049,39 | 889,39 | 1460,45 | 1008,31 | 1070,58 |
| WDR1 | 94275,28 | 71574,84 | 119566,52 | 50274,34 | 111940,66 | 70990,76 | 139327,05 | 103340,55 |
| XRCC5 | 43469,06 | 24333,74 | 17980,04 | 8905,65 | 27448,78 | 20900,96 | 14131,90 | 24457,14 |
| YWHAB | 92018,63 | 72650,60 | 100676,75 | 67069,79 | 580056,88 | 487982,19 | 554679,75 | 668841,50 |
| YWHAE | 66648,79 | 40452,29 | 96472,20 | 48120,46 | 289014,22 | 233115,78 | 248284,45 | 225383,23 |
| YWHAG | 56230,41 | 32144,46 | 20502,10 | 28987,95 | 102480,69 | 97205,55 | 95610,26 | 67250,09 |
| YWHAH | 9101,19 | 1534,02 | 29554,42 | 14313,00 | 39572,31 | 31241,71 | 35722,66 | 22218,07 |
| YWHAQ | 32597,34 | 64698,77 | 68616,48 | 33185,25 | 194343,41 | 168678,88 | 188978,25 | 137420,88 |
| YWHAZ | 26143,84 | 24252,16 | 54275,20 | 24594,42 | 160155,47 | 143355,33 | 159260,16 | 145598,02 |

**Table S3.** **Detailed proteomics-guided drug discovery strategy.** Factors are present in all biological replicates (N=3) of both EVs and SFs. N/A is not applicable.

| **EV-proteins** | **Growth Factors / Cytokines** | **Annotation** | **Receptor** | **Expression in AT1 cells** | **Expression in AT2 cells** | **Signal IP** | **Phobius** | **Conclusion** |
| --- | --- | --- | --- | --- | --- | --- | --- | --- |
|  | **PDGFRB** | Platelet Derived Growth Factor Receptor Beta | N/A | N/A | N/A | 0.9896 | Signal peptide region | Don't investigate, protein is a receptor |
|  | **TNFSF4** | TNF superfamily member 4 | TNFRSF4 | Yes | Yes | 0.0005 | Non-cytoplasmic | Don't investigate, protein is not secreted |

| **Shared proteins** | **Growth Factors / Cytokines** | **Annotation** | **Receptor** | **Expression in AT1 cells** | **Expression in AT2 cells** | **Signal IP** | **Phobius** | **Conclusion** |
| --- | --- | --- | --- | --- | --- | --- | --- | --- |
|  | **C3** | Complement C3 | C3AR1 | No | Low expression | 0.9867 | Signal peptide region | Don't investigate, no receptor expression |
|  | **CAT** | Catalase | N/A | N/A | N/A | 0.0004 | Non- cytoplasmic | Don't investigate, protein is an enzyme |
|  | **GREM1** | Gremlin-1 | EGFR | Yes | Yes | 0.9903 | Signal peptide region | Investigate |
|  | **HGF** | Hepatocyte Growth Factor | MET | Yes | Yes | 0.7707 | Signal peptide region | Investigate |
|  | **LTBP1** | Latent-transforming growth factor beta-binding protein 1 | N/A | N/A | N/A | 0.9446 | Signal peptide region | Don't investigate, is a binding protein |
|  | **LTBP2** | Latent-transforming growth factor beta-binding protein 2 | N/A | N/A | N/A | 0.9175 | Signal peptide region | Don't investigate, is a binding protein |
|  | **OGN** | Osteoglycin / Mimecan | Unknown | N/A | N/A | 0.9916 | Signal peptide region | Investigate, receptor unknown. |
|  | **S100A6** | Protein S100-A6 | AGER | Yes | Yes | 0.0043 | Non- cytoplasmic | Don't investigate, protein is not secreted |
|  | **STC1** | Stanniocalcin-1 | Unknown | N/A | N/A | 0.9763 | Signal peptide region | Investigate, receptor unknown. |
|  | **STC2** | Stanniocalcin-2 | Unknown | N/A | N/A | 0.9868 | Signal peptide region | Investigate, receptor unknown. |
|  | **TNC** | Tenascin-C | VTN | No | No | 0.9765 | Signal peptide region | Don't investigate, no receptor expression |

| **SF-proteins** | **Growth Factors / Cytokines** | **Annotation** | **Receptor** | **Expression in AT1 cells** | **Expression in AT2 cells** | **Signal IP** | **Phobius** | **Conclusion** |
| --- | --- | --- | --- | --- | --- | --- | --- | --- |
|  | **BMP1** | Bone morphogenetic protein 1 | ACVR1 | Yes | Yes | 0.9874 | Signal peptide region | Investigate |
|  |  |  | BMPR1A | Yes | Yes |  |  |  |
|  |  |  | BMPR1B | Low expression | No |  |  |  |
|  | **C5** | Complement C5 | C5AR1 | Low expression | Yes | 0.9977 | Signal peptide region | Don't investigate, no receptor expression |
|  |  |  | C5AR2 | No | Low expression |  |  |  |
|  | **CCN1** | Cellular Communication Network Factor 1 | ITGAV | Yes | Yes | 0.9984 | Signal peptide region | Investigate |
|  | **CCN2** | Cellular communication network factor 2 | FGFR2 | Yes | Yes | 0.9989 | Signal peptide region | Investigate |
|  |  |  | FGFR3 | Yes | Yes |  |  |  |
|  | **CLEC11A** | C-type lectin domain family 11 member A | LEPR | Low expression | Low expression | 0.8789 | Signal peptide region | Don't investigate, no receptor expression |
|  |  |  | Itga11 | No | No |  |  |  |
|  | **CSF1** | Macrophage colony-stimulating factor 1 | CSF1R | No | Low expression | 0.9651 | Signal peptide region | Don't investigate, no receptor expression |
|  | **GDF15** | Growth differentiation factor 15 | GFRAL | No | No | 0.5821 | Signal peptide region | Don't investigate, no receptor expression |
|  | **GPI** | Glucose-6-Phosphate Isomerase | AMFR | Yes | Yes | 0.0005 | Non- cytoplasmic | Don't investigate, protein is not secreted |
|  | **HDGF** | Hepatoma-derived growth factor | Unknown | N/A | N/A | 0.0005 | Non- cytoplasmic | Don't investigate, protein is not secreted |
|  | **INHBA** | Inhibin beta A chain | ACVR2A | Yes | Yes | 0.9704 | Signal peptide region | Investigate |
|  | **LTBP3** | Latent-transforming growth factor beta-binding protein 3 | N/A | N/A | N/A | 0.7515 | Signal peptide region | Don't investigate, is a binding protein |
|  | **LTBP4** | Latent-transforming growth factor beta-binding protein 4 | N/A | N/A | N/A | 0.9955 | Signal peptide region | Don't investigate, is a binding protein |
|  | **MIF** | Macrophage migration inhibitory factor | CD74 | Yes | Yes | 0.0019 | Non- cytoplasmic | Don't investigate, protein is not secreted |
|  |  |  | CXCR2 | No | No |  |  |  |
|  |  |  | CXCR4 | Low expression | Low expression |  |  |  |
|  | **NAMPT** | Nicotinamide phosphoribosyltransferase | TLR4 | No | Low expression | 0.0019 | Non-cytoplasmic | Don't investigate, protein is not secreted |
|  | **PDGFC** | Platelet-derived growth factor C | KDR | No | No | 0.9361 | Signal peptide region | Don't investigate, no receptor expression |
|  |  |  | FLT4 | No | No |  |  |  |
|  |  |  | PDGFRA | N/A | N/A |  |  |  |
|  | **PDGFD** | Platelet-derived growth factor D | PDGFRB | No | No | 0.9414 | Signal peptide region | Don't investigate, no receptor expression |
|  | **PLAU** | Urokinase-type plasminogen activator | PLAUR | Yes | Yes | 0.9993 | Signal peptide region | Don't investigate, protein is an enzyme |
|  | **PTN** | Pleiotrophin | SDC3 | Yes | Yes | 0.9893 | Signal peptide region | Investigate |
|  |  |  | RPTPß | No | Low expression |  |  |  |
|  |  |  | ALK | No | No |  |  |  |
|  |  |  | NRP1 | No | Yes |  |  |  |
|  |  |  | ITGB3 | No | No |  |  |  |
|  | **SBDS** | Ribosome maturation protein SBDS | N/A | N/A | N/A | 0.0002 | Non- cytoplasmic | Don't investigate, protein is not secreted |
|  | **SEMA3A** | Semaphorin-3A | CLCP1 | N/A | N/A | 0.9797 | Signal peptide region | Investigate, receptor expression unknown. |
|  | **SEMA4B** | Semaphorin-4B | Neuropilin-1 | No | Low expression | 0.9964 | Signal peptide region | Don't investigate, no receptor expression |
|  | **SEMA5A** | Semaphorin-5A | PLXNB3 | No | No | 0.9753 | Signal peptide region | Don't investigate, no receptor expression |
|  | **SEMA5B** | Semaphorin-5B | PLXNA1 | Yes | Yes | 0.0147 | Cytoplasma and non- cytoplasmic | Don't investigate, protein is not secreted |
|  | **SEMA7A** | Semaphorin 7a | PLXNC1 | No | Low expression | 0.8971 | Signal peptide region | Investigate |
|  |  |  | ITGB1 | Yes | Yes |  |  |  |
|  | **TGFB1** | Transforming growth factor beta-1 | TGFBR1 | Yes | Yes | 0.9872 | Signal peptide region | Don't investigate, is a COPD-related protein |
|  |  |  | TGFBR2 | Yes | Yes |  |  |  |
|  |  |  | TGFBR3 | Yes | Yes |  |  |  |
|  | **TGFB2** | Transforming growth factor beta-2 | TGFBR1 | Yes | Yes | 0.8802 | Signal peptide region | Don't investigate, is a COPD-related protein |
|  |  |  | TGFBR2 | Yes | Yes |  |  |  |
|  |  |  | TGFBR3 | Yes | Yes |  |  |  |
|  | **TNFRSF11B** | Tumor necrosis factor receptor superfamily member 11B | N/A | N/A | N/A | 0.9468 | Signal peptide region | Don't investigate, protein is a receptor |
|  | **TXLNA** | Alpha-taxilin / IL-14 | IL14R | N/A | N/A | 0.0014 | Non- cytoplasmic | Don't investigate, protein is not secreted |

**Table S4. Information of recombinant proteins screened in organoid assay.**

| **Growth factor / Cytokine** | **Annotation** | **Final concentration in organoid assay** | **Article No.** | **Company** | **Reference for chosen final concentration** |
| --- | --- | --- | --- | --- | --- |
| BMP1 | Bone morphogenetic protein 1 | 50 ng/mL | 1927-ZN | Biotechne | 10.1016/S0896-6273(00)00148-3 |
| CCN1 | Cellular communication network 1 | 4 µg/mL | 120-25 | Peprotech | 10.1038/s41467-022-30851-1 |
| CCN2 | Cellular communication network 2 | 50 ng/mL | 120-19 | Peprotech | 10.3390/ijms23010375 |
| GREM1 | Gremlin-1 | 100 ng/mL | 120-42 | Peprotech | 10.1053/j.gastro.2021.03.052 |
| HGF | Hepatocyte growth factor | 10 ng/mL | 100-39H | Peprotech | 10.1016/j.cellsig.2017.09.001 |
| INHBA | Inhibin | 5 ng/mL | ORB1676755 | Biorbyt | 10.1084/jem.20211887 |
| OGN | Osteoglycin | 10 µg/mL | ORB383003 | Biorbyt | 10.1074/jbc.M111.292193 |
| PTN | Pleiotrophin | 25 ng/mL | 450-15 | Peprotech | 10.1038/s41380-021-01316-6 |
| SEMA3A | Semaphorin 3a | 1 µg/mL | 150-17H | Peprotech | 10.1038/s41598-021-81335-z |
| SEMA7A | Semaphorin 7a | 250 ng/mL | CSB-YP021000HU | Cusabio | 10.1152/ajplung.00400.2018 |
| STC1 | Stanniocalcin 1 | 100 ng/mL | CSB-MP022821HU | Cusabio | 10.1074/jbc.M506667200 |
| STC2 | Stanniocalcin 2 | 500 ng/mL | 9405-SO | Bio Techne | 10.18632/oncotarget.12147 |

**Table S5A. Patient characteristics of immunohistochemistry OGN staining in control tissue.**

The FEV_1_ and FVC data were not available for lung tissue from Rochester (N=7) and for a few cases from Groningen (N=5). Data is presented as the median with the range between brackets. FEV_1%pred_ = highest measurement of forced expiratory volume in 1s for an individual; FVC = forced vital capacity; NA = not applicable; * = a significant difference in the number of pack years between current and ex-smokers (unpaired two-tailed T-test).

|  | **Never-smoker** | **Current smoker** | **Ex-smoker** |
| --- | --- | --- | --- |
| **Sex (Female / Male)** | 10 / 3 | 7 / 2 | 5 / 6 |
| **Age in years** | 60.00 (45.00 – 70.00) | 61.00 (51.00 – 72.00) | 62.00 (50.00 – 71.00) |
| **Pack years** | NA | 40.00 (20.00 – 81.00)* | 20.00 (10.00 – 51.00)* |
| **FEV_1%pred_** | 87.26 (80.00 – 114.40) | 102.50 (87.91 – 106.00) | 99.00 (84.96 – 127.00) |
| **FEV_1_/FVC_max_** | 80.60 (73.00 – 84.00) | 73.72 (70.55 – 90.13) | 79.00 (69.00 – 86.63) |

**Table S5B. Patient characteristics for immunohistochemistry OGN staining in COPD tissue.**

SEO-COPD (N=12) and moderate-severe COPD (n=14) were matched in terms of age, sex, and smoking status to respective control groups. Data is presented as the median with the range between brackets. FEV_1%pred_ = highest measurement of forced expiratory volume in 1s for an individual; FVC = forced vital capacity; NA = not applicable. Statistical significance was determined with an unpaired two-tailed T-test; p-values <0.05 were considered significant.

|  | **Younger control** | **SEO-COPD** | **p value** |
| --- | --- | --- | --- |
| **Sex (Female/Male)** | 6 / 3 | 8 / 4 | >0.99 |
| **Age in years** | 55.00 (43.00 – 62.00) | 51.50 (47.00 – 55.00) | 0.36 |
| **Pack years** | 20.00 (1.50 – 52.50) | 22.00 (5.00 – 54.00) | >0.99 |
| **FEV_1%pred_** | 102.20 (91.00 – 127.00) | 16.38 (12.00 – 29.90) | <0.0001 |
| **FEV_1_/FVC_max_** | 76.00 (69.00 – 86.63) | 27.28 (20.52 – 68.00) | <0.0001 |
|  | **Older control** | **Moderate-severe COPD** | **p value** |
| **Sex (Female/Male)** | 3 / 6 | 2 / 12 | 0.29 |
| **Age in years** | 74.00 (67.00 – 81.00) | 72.00 (67.00 – 81.00) | 0.71 |
| **Pack years** | 25.50 (5.00 – 52.50) | 41.00 (10.00 – 79.00) | 0.51 |
| **FEV_1%pred_** | 89.24 (75.85 – 133.00) | 62.03 (49.66 – 75.36) | <0.0001 |
| **FEV_1_/FVC_max_** | 72.00 (69.00 – 86.15) | 60.84 (43.40 – 67.90) | <0.0001 |

**Table S6. Overview of the genes included for the gene signatures of alveolar type II cells and fibroblasts**

| **AT2 cells** | **Fibroblasts** |
| --- | --- |
| *Sftpb* | *Dcn* |
| *Sftpc* | *Lum* |
| *Sftpa1* | *Fbln1* |
| *Napsa* | *Col1a2* |
| *Slc34a2* | *Col6a2* |
| *Sftpd* | *Col6a3* |
| *Ctsh* | *Mmp2* |
| *Lrrk2* | *Fn1* |
| *Hopx* | *Col6a1* |
| *Lamp3* | *Adh1b* |

**Table S7. Patient characteristics of lung tissue from COPD IV patients used for organoid culture**

The FEV_1_ and FVC data were unavailable for lung tissue from one donor patient. Data is presented as the median with the range between brackets. FEV1 = highest measured forced expiratory volume in 1s for an individual after bronchodilation (BD); FVC = forced vital capacity.

| **Sex**  M/F | **Age**  Median (range) | **Smoking status** | **Stop since (years)**  Median (range) | **Pack years**  Median (range) | **FEV1 (% of predicted post-BD)**  Median (range) | **FEV1/FVC (% of predicted post-BD)**  Median (range) |
| --- | --- | --- | --- | --- | --- | --- |
| 5/6 | 61 (55 – 68) | All ex-smokers | 10 (3 – 15) | 30 (18 – 45) | 16 (11 – 27)  2 unknown | 25.6 (17 – 29)  2 unknown |
